## Supplementary figures for "A wolf in another wolf’s clothing: Post-genomic regulation dictates venom profiles of medically-important cryptic kraits in India"

**Figure S1:** The Bayesian cytochrome *b* phylogeny of *Bungarus* species.

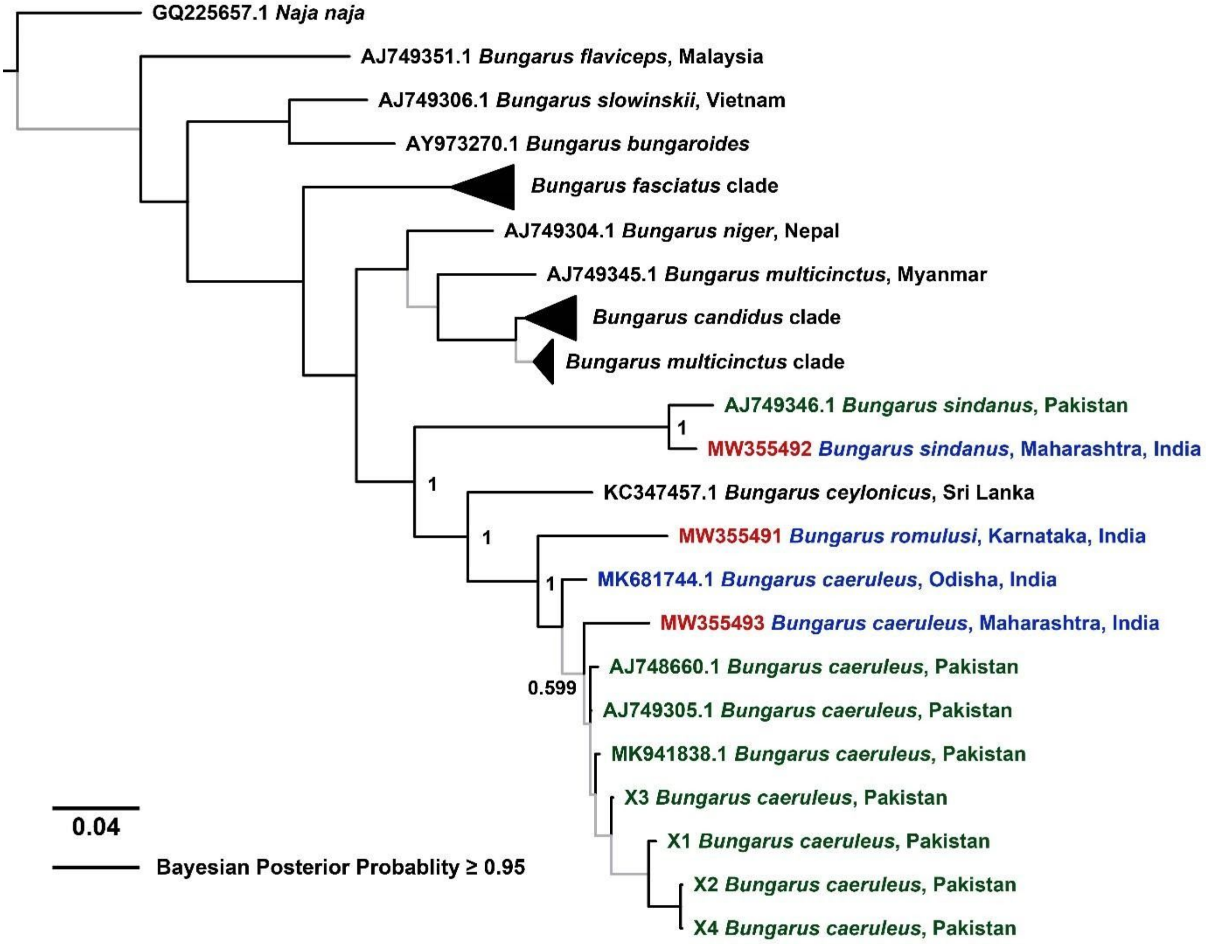


The Bayesian cytochrome *b* phylogeny of *Bungarus* species in Asia is shown here. Terminal labels coloured in blue and green represent krait lineages from India and Pakistan, respectively, and the accession numbers of individuals sequenced in this study have been highlighted in red. Well-supported branches (BPP ≥ 0.95) are shown in thick black lines, and the node support for clades of interest are indicated. The scale for branch lengths (number of nucleotide substitutions per site) is also presented.

**Figure S2A:** Maximum likelihood phylogeny (ND4 marker) for *Bungarus* species.

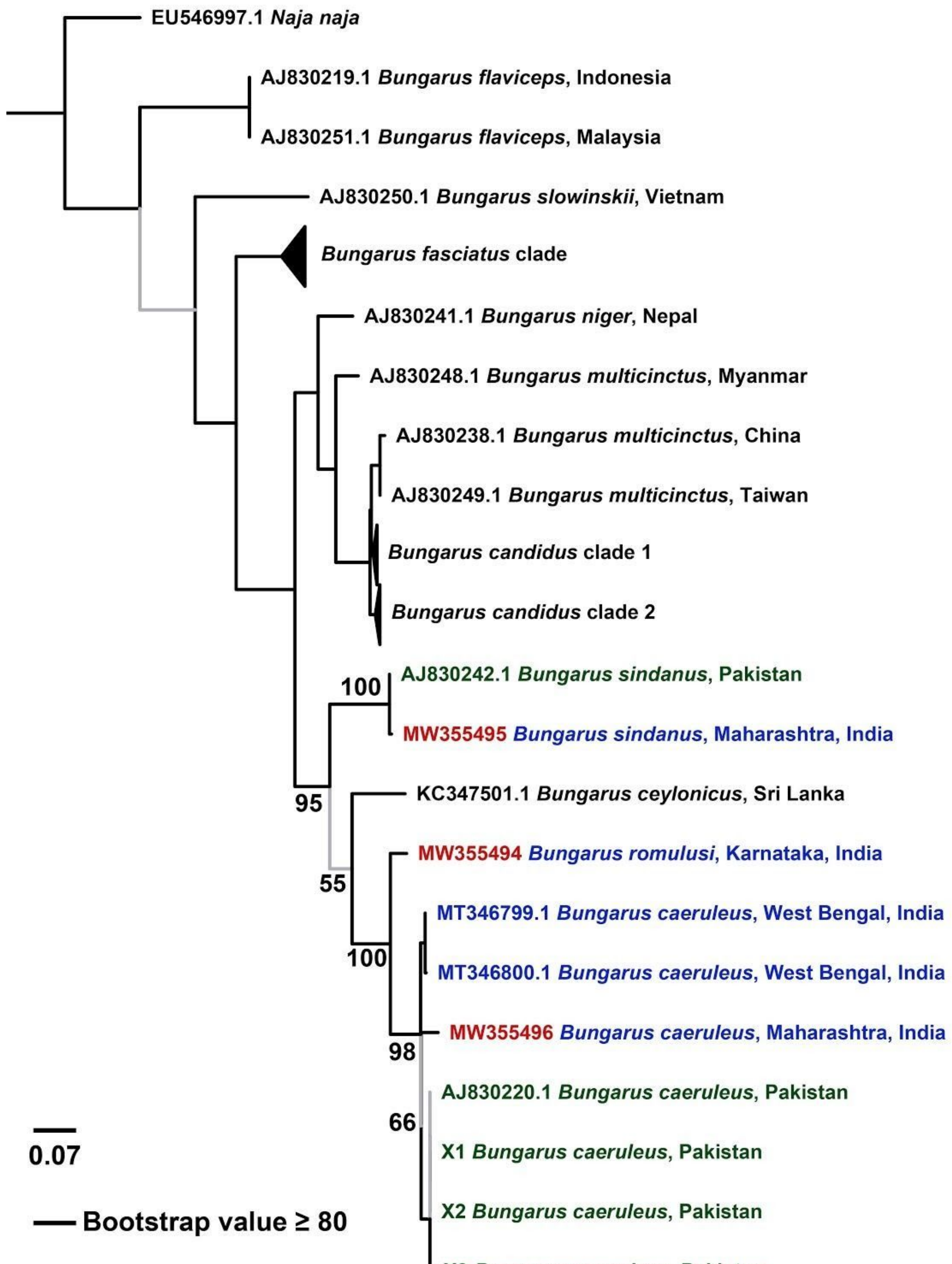


This tree depicts the relationship between various species of kraits found in Southeast Asia. Terminal labels coloured in blue and green represent krait lineages from India and Pakistan, respectively, and the accession numbers of individuals sequenced in this study have been highlighted in red. Well-supported branches (BS ≥ 80) are shown in thick black lines, and the node support for clades of interest are indicated. The scale for branch lengths (number of nucleotide substitutions per site) is also presented.

**Figure S2B:** Maximum likelihood phylogeny (cyt *b* marker) for *Bungarus* species.

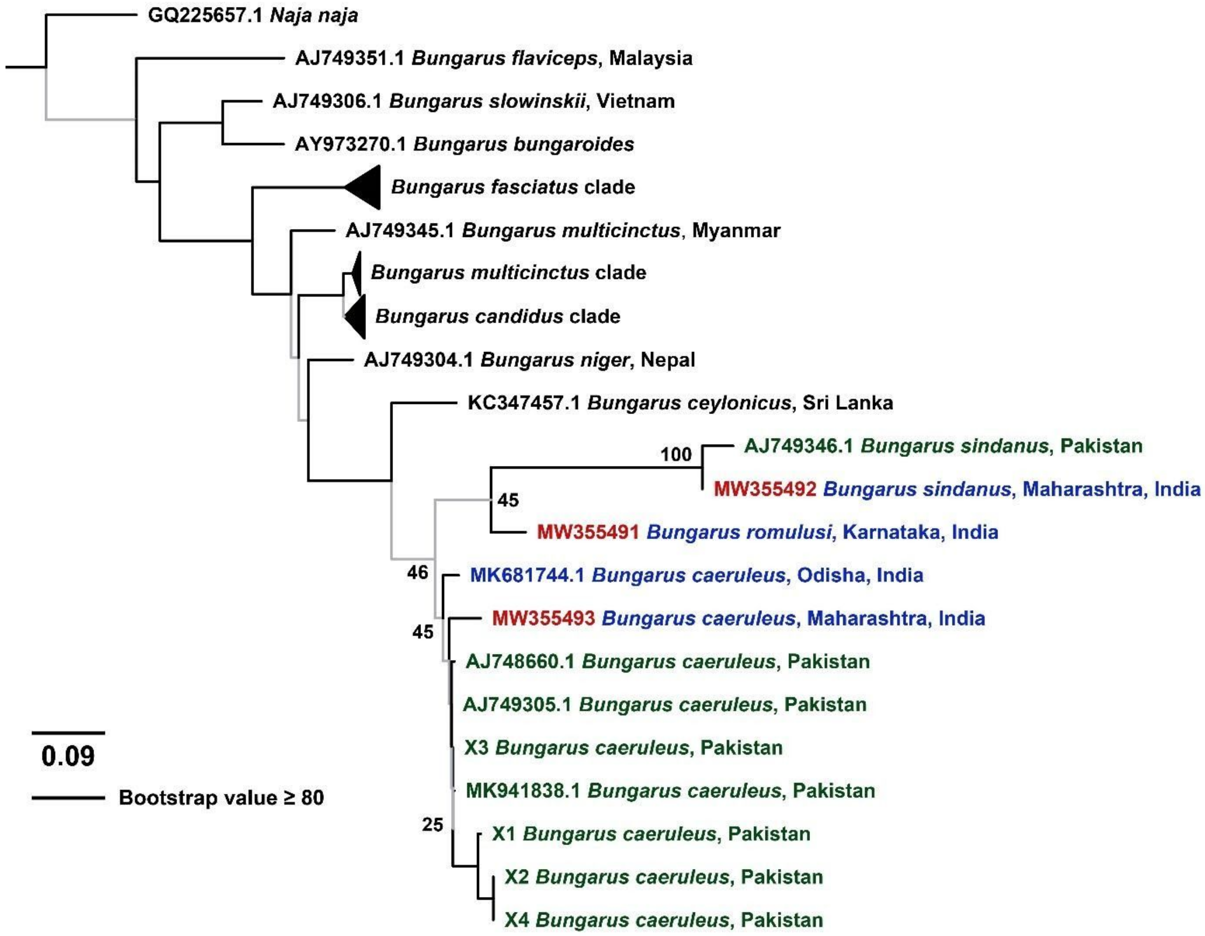


Built using cytochrome *b* sequences, this ML tree depicts the relationship between various species of kraits found in Southeast Asia. Terminal labels coloured in blue and green represent krait lineages from India and Pakistan, respectively, and the accession numbers of individuals sequenced in this study have been highlighted in red. Well-supported branches (BS ≥ 80) are shown in thick black lines, and the node support for clades of interest are indicated. The scale for branch lengths (number of nucleotide substitutions per site) is also presented.

**Figure S3:** Electrophoretic separation of *Bungarus* venoms under non-reducing and reducing conditions.

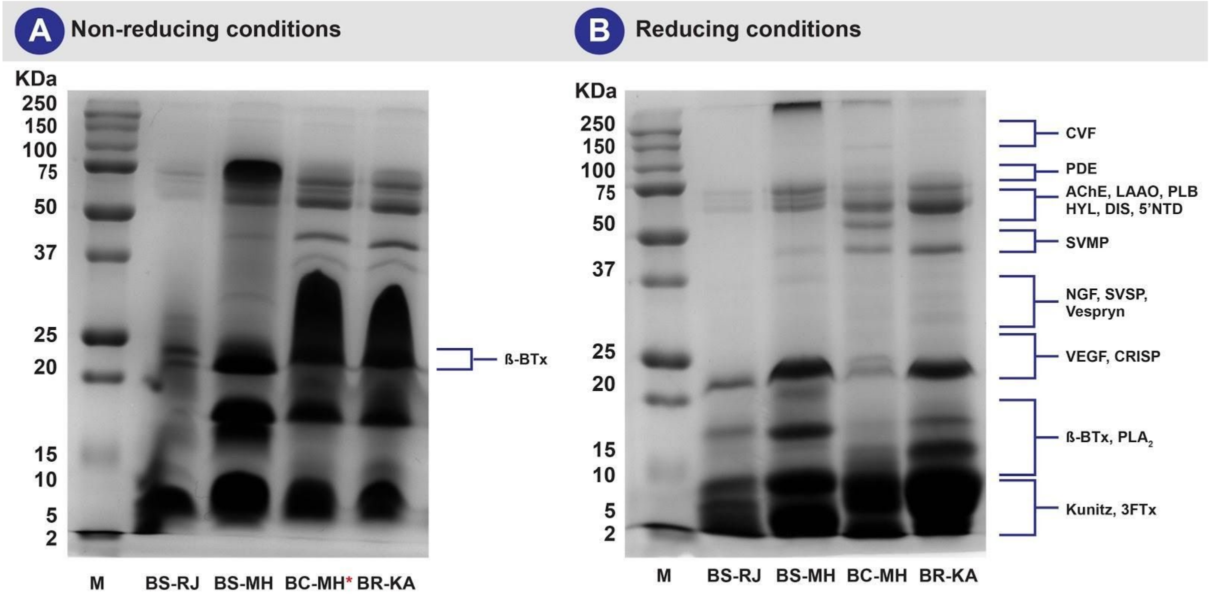


This figure depicts the SDS-PAGE profiles of *Bungarus* venoms under **(A)** non-reducing and **(B)** reducing conditions. Lanes 1 to 5 represent the marker (2 - 250 KDa), *B. sindanus* (Rajasthan and Maharashtra), *B. caeruleus* (Maharashtra) and *B. romulusi* (Karnataka).

*A pooled venom sample of *B. caeruleus* from Maharashtra has been used as a representative population

**Figure S4A-C.** Pharmacological activities of *Bungarus* venoms from Western India.

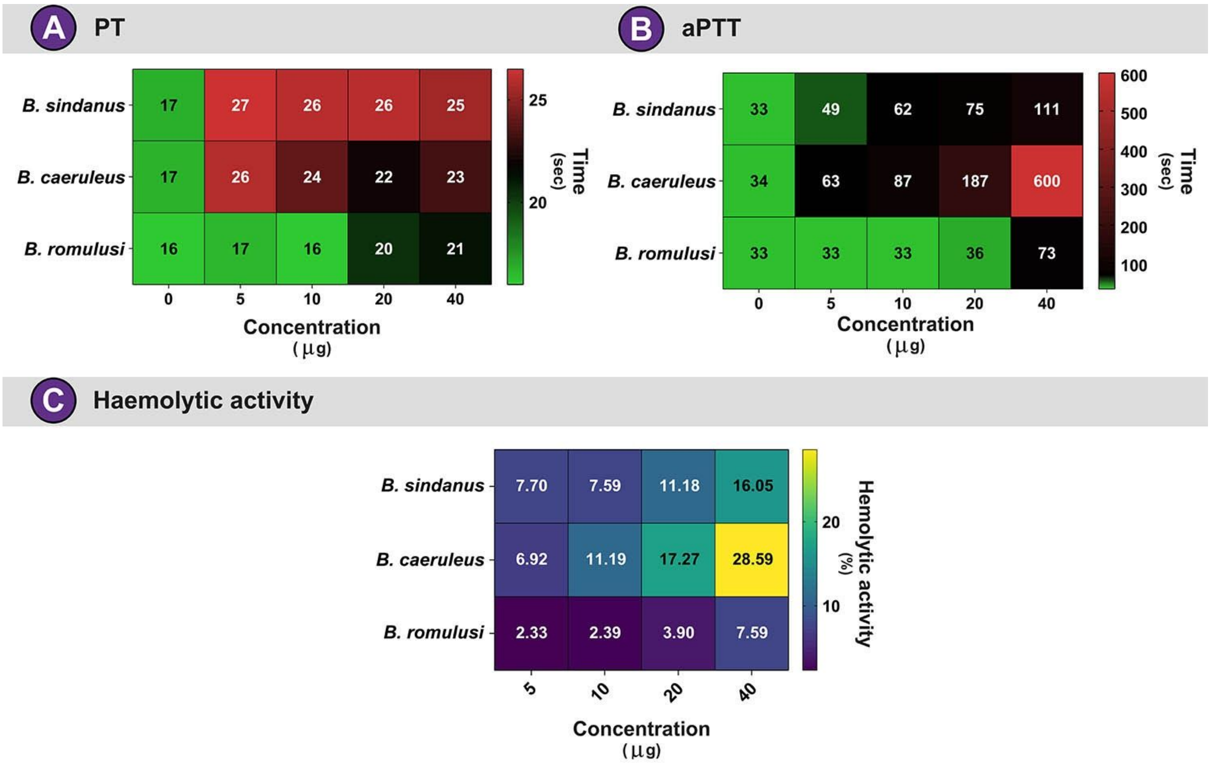


The effect of various concentrations (0, 5, 10, 20, 40 µg) of *Bungarus* venoms on the **(A)** extrinsic and **(B)** intrinsic pathways of blood coagulation, respectively, and **(C)** haemolysis are depicted in this figure as heatmaps. The time taken (in seconds) for the initial fibrin clot formation is represented by the numbers inside each square in panels A and B. The relative haemolytic activity (in %) calculated in relation to the activity of the positive control (0.5% Triton X) is represented by the numbers inside the squares in panel C.

**Figure S5.** Western blotting of Indian polyvalent antivenoms against the cryptic *Bungarus* venoms.

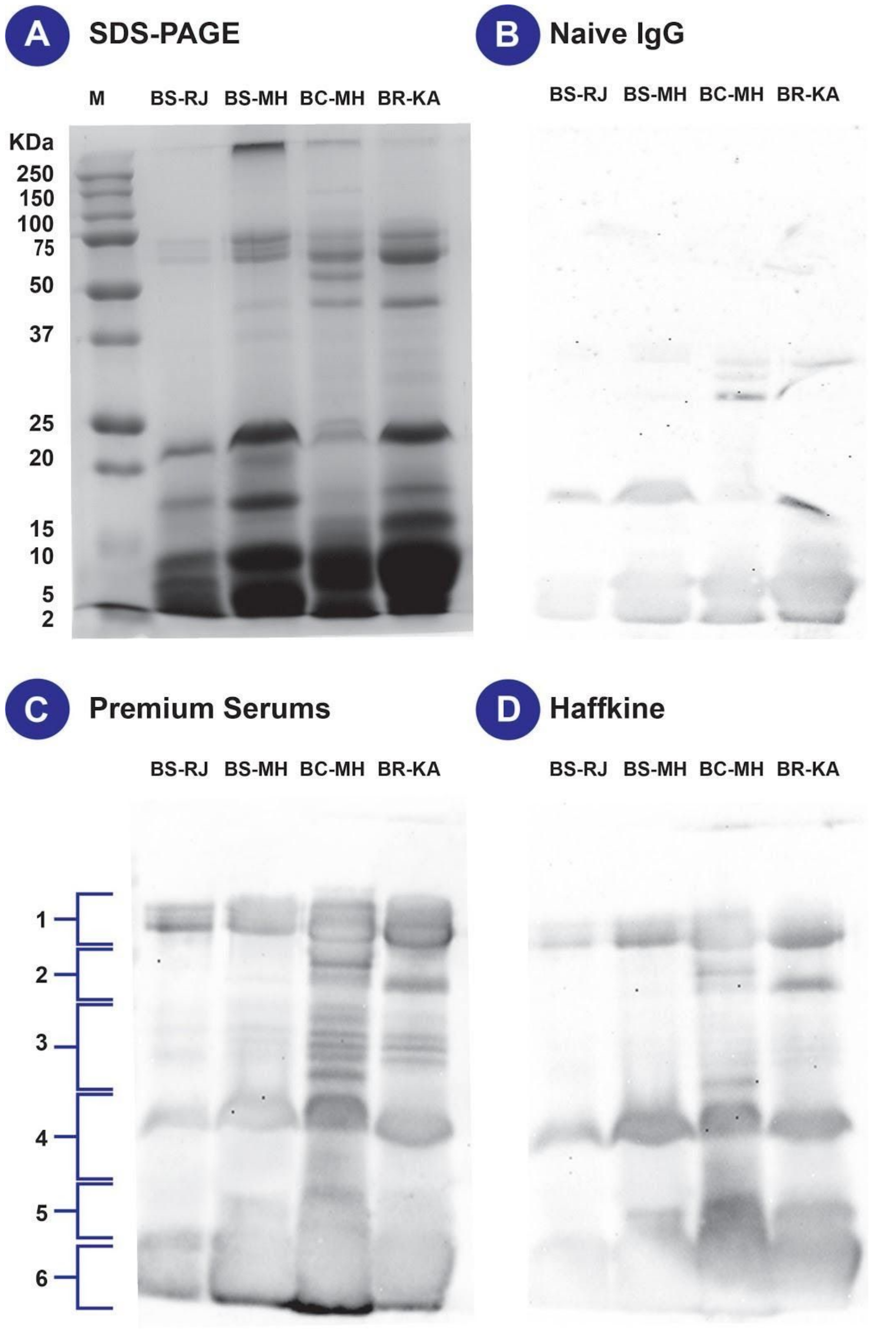


The recognition capability of commercial antivenoms against toxins (bands 1 - 6) in the venoms of *B. sindanus*, Rajasthan (BS-RJ); *B. sindanus,* Maharashtra (BS-MH); *B. caeruleus*, Maharashtra (BC-MH); and *B. romulusi*, Karnataka (BR-KA) is depicted here. Panels represent **(A)** SDS-PAGE, **(B)** Naive IgG, **(C)** Premium Serums and **(D)** Haffkine antivenoms. A standard protein marker (**M)** was run along with the venoms.
