## Supplementary tables for "A wolf in another wolf’s clothing: Post-genomic regulation dictates venom profiles of medically-important cryptic kraits in India"

**Table S1A.** Details of *Bungarus* venom samples investigated in this study.

| | No. of individuals | Protein concentration (mg/ml) | Sampling location |
|---|---|---|---|
| ***B. sindanus**** | 2 | 0.274 | Bikaner, Rajasthan |
| ***B. sindanus*** | 1 | 0.128 | Pune, Maharashtra |
| ***B. caeruleus*** | 1 | 0.099 | Pune, Maharashtra |
| ***B. romulusi*** | 1 | 0.122 | Bannerghatta, Karnataka |

*This sample was analysed previously [15].

**Table S1B.** Details of antivenom samples investigated in this study.

| **Manufacturer** | **Batch** | **Manufacture (M) and expiry (E) dates** | **Protein content** (mg/ml) | **Marketed neutralizing efficacy** (mg/ml) |
|---|---|---|---|---|
| **Haffkine BioPharmaceutical Corporation Ltd.** | AS180611 | **M:** 06/2018<br>**E:** 11/2022 | 24.7 ± 0.5 | *N. naja*: 0.60<br>*B. caeruleus*: 0.45<br>*D. russelii*: 0.60<br>*E. carinatus*: 0.45 |
| **Premium Serums & Vaccines Pvt. Ltd.** | ASVS(I) Lyo013 | **M:** 11/2018<br>**E:** 11/2022 | 26.2 ± 1.2 | |

Details of the commercial Indian polyvalent antivenoms tested in this study are provided here. Batch numbers, manufacturing and expiry dates, protein concentrations, and marketed neutralising potencies are shown.

**Table S2:** Details of primers used in the amplification of the mitochondrial markers.

| Sr. no. | Gene | Forward Primer Name | Forward Primer Sequence (5'-3') | Reverse Primer Name | Reverse Primer sequence (5'-3') | Amplicon Size | Annealing Temperature | Cycles | Reference |
|---|---|---|---|---|---|---|---|---|---|
| 1. | ND4 | ND4I | TGACTACCAAAAGC TCATGTAGAAGC | tRNA-Leu | TACTTTTACTTGGA TTTGCACCA | 820bp | 50°C | 35 | [53] |
| 2. | Cytochrome B | L14910 | GACCTGTGATMTG AAAACCAYCGTTGT | H16064 | CTTTGGTTTACAAG AACAATGCTTTA | 1117bp | 46°C | 35 | [54] |

**Table S3A: Evolutionary sequence divergence estimated using the maximum composite likelihood (MCL) model for ND4 marker.**

| | AJ830220.1 *Bungarus caeruleus* Pakistan | AJ830242.1 *Bungarus sindanus* Pakistan | *Bungarus caeruleus* Pakistan 01 | *Bungarus caeruleus* Pakistan 02 | *Bungarus caeruleus* Pakistan 03 | *Bungarus caeruleus* Pakistan 04 | MT346799.1 *Bungarus caeruleus* West Bengal India | MT346800.1 *Bungarus caeruleus* West Bengal India | MW355496 *Bungarus caeruleus* Maharashtra India | MW355494 *Bungarus romulusi* Karnataka India | MW355495 *Bungarus sindanus* Maharashtra India |
|---|---|---|---|---|---|---|---|---|---|---|---|
| AJ830220.1 *Bungarus caeruleus* Pakistan | 0 | 15.3 | 0 | 0 | 0.34 | 0.52 | 2.27 | 2.33 | 3.73 | 7.89 | 16.17 |
| AJ830242.1 *Bungarus sindanus* Pakistan | | 0 | 15.97 | 15.97 | 15.98 | 15.77 | 15.63 | 15.32 | 16.66 | 14.37 | **0.31** |
| *Bungarus caeruleus* Pakistan 01 | | | 0 | 0 | 0.34 | 0.52 | 2.37 | 2.28 | 2.35 | 7.72 | 16.42 |
| *Bungarus caeruleus* Pakistan 02 | | | | 0 | 0.34 | 0.52 | 2.37 | 2.28 | 2.35 | 7.72 | 16.42 |
| *Bungarus caeruleus* Pakistan 03 | | | | | 0 | 0.17 | 2.74 | 2.64 | 2.73 | 8.11 | 16.43 |
| *Bungarus caeruleus* Pakistan 04 | | | | | | 0 | 2.93 | 2.82 | 2.91 | 8.30 | 16.21 |
| MT346799.1 *Bungarus caeruleus* West Bengal India | | | | | | | 0 | 0.15 | 2.69 | 7.09 | 16.28 |
| MT346800.1 *Bungarus caeruleus* West Bengal India | | | | | | | | 0 | 3.25 | 6.96 | 16.02 |
| MW355496 *Bungarus caeruleus* Maharashtra India | | | | | | | | | 0 | **8.61** | 15.66 |
| MW355494 *Bungarus romulusi* Karnataka India | | | | | | | | | | 0 | **14.87** |
| MW355495 *Bungarus sindanus* Maharashtra India | | | | | | | | | | | 0 |

This table shows the evolutionary distance between *Bungarus* species in Asia for ND4 marker calculated using maximum composite likelihood (MCL) model. Distances between species of interest have been highlighted in blue. Accession numbers of individuals sequenced in this study have been marked in red.

**Table S3B: Evolutionary sequence divergence estimated using the MCL model for cytochrome b marker.**

| | AJ749305.1 *Bungarus caeruleus* Pakistan | MK941838.1 *Bungarus caeruleus* Pakistan | MK681744.1 *Bungarus caeruleus* India | *Bungarus caeruleus* Pakistan 01 | *Bungarus caeruleus* Pakistan 02 | *Bungarus caeruleus* Pakistan 03 | *Bungarus caeruleus* Pakistan 04 | AJ749346.1 *Bungarus sindanus* Pakistan | MW355493 *Bungarus caeruleus* Maharashtra India | MW355491 *Bungarus romulusi* Karnataka India | MW355492 *Bungarus sindanus* Maharashtr a India |
|---|---|---|---|---|---|---|---|---|---|---|---|
| AJ749305.1 *Bungarus caeruleus* Pakistan | 0 | 0.29 | 2.86 | 1.38 | 4.23 | 0.16 | 4.23 | 28.75 | 3.93 | 10.07 | 20.66 |
| MK941838.1 *Bungarus caeruleus* Pakistan | | 0 | 2.65 | 1.13 | 2.08 | 0.16 | 2.08 | 17.55 | 5.61 | 6.63 | 18.01 |
| MK681744.1 *Bungarus caeruleus* India | | | 0 | 2.86 | 2.86 | 2.88 | 2.86 | 18.74 | 3.42 | 5.94 | 19.56 |
| *Bungarus caeruleus* Pakistan 01 | | | | 0 | 0.83 | 0 | 0.83 | 17.6 | 3.69 | 8.83 | 18.16 |
| *Bungarus caeruleus* Pakistan 02 | | | | | 0 | 0 | 0 | 29.72 | 6.55 | 12.04 | 23.03 |
| *Bungarus caeruleus* Pakistan 03 | | | | | | 0 | 0 | 16.45 | 6.35 | 6.67 | 16.96 |
| *Bungarus caeruleus* Pakistan 04 | | | | | | | 0 | 29.72 | 6.55 | 12.04 | 23.03 |
| AJ749346.1 *Bungarus sindanus* Pakistan | | | | | | | | 0 | 32.27 | 24.08 | **3.03** |
| MW355493 *Bungarus caeruleus* Maharashtra India | | | | | | | | | 0 | **14.26** | 24.24 |
| MW355491 *Bungarus romulusi* Karnataka India | | | | | | | | | | 0 | **19.96** |

| | | | | | | | | | | | |
|---|---|---|---|---|---|---|---|---|---|---|---|
| MW355492 *Bungarus sindanus* Maharashtra India | | | | | | | | | | | 0 |

This table shows the evolutionary distance between *Bungarus* species in Asia for cytochrome b marker calculated using maximum composite likelihood (MCL) model. Distances between species of interest have been highlighted in blue. Accession numbers of individuals sequenced in this study have been marked in red.

**Table S3C: Evolutionary divergence estimated as p-distance for ND4 marker.**

| | AJ830221 *Bungarus caeruleus* Pakistan | AJ830242.1 *Bungarus sindanus* Pakistan | *Bungarus caeruleus* Pakistan 01 | *Bungarus caeruleus* Pakistan 02 | *Bungarus caeruleus* Pakistan 03 | *Bungarus caeruleus* Pakistan 04 | MT346799.1 *Bungarus caeruleus* West Bengal India | MT346801 *Bungarus caeruleus* West Bengal India | MW355496 *Bungarus caeruleus* Maharashtra India | MW355494 *Bungarus romulusi* Karnataka India | MW355495 *Bungarus sindanus* Maharashtra India |
|---|---|---|---|---|---|---|---|---|---|---|---|
| AJ830221 *Bungarus caeruleus* Pakistan | 0 | 13.48 | 0 | 0 | 0.34 | 0.51 | 2.23 | 2.29 | 0.35 | 7.42 | 14.15 |
| AJ830242.1 *Bungarus sindanus* Pakistan | | 0 | 13.98 | 13.98 | 13.98 | 13.81 | 13.71 | 13.45 | 14.39 | 12.82 | **0.31** |
| *Bungarus caeruleus* Pakistan 01 | | | 0 | 0 | 0.34 | 0.51 | 2.32 | 2.24 | 2.31 | **7.25** | 14.33 |
| *Bungarus caeruleus* Pakistan 02 | | | | 0 | 0.34 | 0.51 | 2.32 | 2.24 | 2.31 | 7.25 | 14.33 |
| *Bungarus caeruleus* Pakistan 03 | | | | | 0 | 0.17 | 2.68 | 2.59 | 2.67 | 7.59 | 14.33 |
| *Bungarus caeruleus* Pakistan 04 | | | | | | 0 | 2.86 | 2.76 | 2.85 | 7.77 | 14.16 |
| MT346799.1 *Bungarus caeruleus* West Bengal India | | | | | | | 0 | 0.15 | 2.62 | 6.69 | 14.21 |
| MT346801 *Bungarus caeruleus* West Bengal India | | | | | | | | 0 | 3.14 | 6.57 | 13.99 |
| MW355496 *Bungarus caeruleus* Maharashtra India | | | | | | | | | 0 | **7.98** | 13.75 |
| MW355494 *Bungarus romulusi* Karnataka India | | | | | | | | | | 0 | **13.20** |
| MW355495 *Bungarus sindanus* Maharashtra India | | | | | | | | | | | 0 |

This table shows the evolutionary distance between *Bungarus* species in Asia for ND4 marker estimated as p-distance. Distances between species of interest have been highlighted in blue. Accession numbers of individuals sequenced in this study have been marked in red.

**Table S3D: Evolutionary divergence estimated as p-distance for cytochrome *b* marker.**

| | AJ749305.1 *Bungarus caeruleus* Pakistan | MK681744.1 *Bungarus caeruleus* India | *Bungarus caeruleus* Pakistan 01 | *Bungarus caeruleus* Pakistan 02 | *Bungarus caeruleus* Pakistan 03 | *Bungarus caeruleus* Pakistan 04 | AJ749346.1 *Bungarus sindanus* Pakistan | MW355493 *Bungarus caeruleus* Maharashtra India | MW355491 *Bungarus romulusi* Karnataka India | MW355492 *Bungarus sindanus* Maharashtra India |
|---|---|---|---|---|---|---|---|---|---|---|
| AJ749305.1 *Bungarus caeruleus* Pakistan | 0 | 2.75 | 1.34 | 3.79 | 0.16 | 3.79 | 18.95 | 3.05 | 8.71 | 15.54 |
| MK681744.1 *Bungarus caeruleus* India | | 0 | 2.75 | 2.75 | 2.77 | 2.75 | 14.53 | 3.28 | 5.51 | 15.03 |
| *Bungarus caeruleus* Pakistan 01 | | | 0 | 0.67 | 0 | 0.67 | 13.63 | 3.47 | 7.79 | 13.97 |
| *Bungarus caeruleus* Pakistan 02 | | | | 0 | 0 | 0 | 18.92 | 5.73 | 9.98 | 16.29 |
| *Bungarus caeruleus* Pakistan 03 | | | | | 0 | 0 | 13.26 | 4.59 | 6.13 | 13.59 |
| *Bungarus caeruleus* Pakistan 04 | | | | | | 0 | 18.92 | 5.73 | 9.98 | 16.29 |
| AJ749346.1 *Bungarus sindanus* Pakistan | | | | | | | 0 | 19.36 | 17.28 | **2.87** |
| MW355493 *Bungarus caeruleus* Maharashtra India | | | | | | | | 0 | **10.92** | 16.38 |

| | | | | | | | | | | |
|---|---|---|---|---|---|---|---|---|---|---|
| MW355491 *Bungarus romulusi* Karnataka India | | | | | | | | | 0 | **15.38** |
| MW355492 *Bungarus sindanus* Maharashtra India | | | | | | | | | | 0 |

This table shows the evolutionary distance between *Bungarus* species in Asia for cytochrome b marker estimated as p-distance. Distances between species of interest have been highlighted in blue. Accession numbers of individuals sequenced in this study have been marked in red.

**Table S4A, S4B and S4C:** Venom proteomic compositions of cryptic *Bungarus* species.

Peaks Studio X Plus was used to examine raw MS/MS spectra against the National Center for Biotechnology Information's (NCBI) non-redundant (nr) database (Serpentes: 8570) for the identification of various toxin families present in the venom. The tables illustrates the key statistics of these searches, including the accession number, species name, -10lgP values, number of high confidence peptides, unique peptides, percent abundance of each toxin hit, average molecular mass (KDa), the toxin family of the matching NCBI entry.

**S4A Table.** The proteomic composition of *B. sindanus* venom.

| Sr. No. | Accession | Species | -10lgP | #Peptides | #Unique | Relative abundance of toxin hit (%) | Avg. Mass (KDa) | Family |
|---|---|---|---|---|---|---|---|---|
| **Phospholipase A2 ($PLA_2$): 52.57%** | | | | | | | | |
| 1 | ABM88801 | *Hoplocephalus stephensii* | 300.23 | 19 | 14 | 27.550% | 11.66 | PLA2 |
| 2 | ACY68711 | *Parasuta nigriceps* | 421.05 | 38 | 37 | 14.775% | 17.26 | PLA2 |
| 3 | AAB24834 | *Bungarus fasciatus* | 338.21 | 24 | 23 | 8.711% | 16.66 | PLA2 |
| 4 | ABG75909 | *Bungarus fasciatus* | 310.13 | 14 | 7 | 1.175% | 9.80 | PLA2 |
| 5 | BAC77656.1 | *Bungarus flaviceps* | 96.75 | 1 | 1 | 0.114% | 15.88 | PLA2 |
| 6 | ADF50035.1 | *Bungarus flaviceps* | 96.75 | 1 | 1 | 0.114% | 15.85 | PLA2 |
| 7 | Q7T2Q4.1 | *Bungarus flaviceps flaviceps* | 96.75 | 1 | 1 | 0.114% | 15.88 | PLA2 |
| 8 | ACY68711.1 | *Parasuta nigriceps* | 87.44 | 1 | 1 | 0.003% | 15.46 | PLA2 |
| 9 | ABU68503.1 | *Trimorphodon biscutatus* | 125.75 | 3 | 1 | 0.001% | 16.62 | PLA2 |
| 10 | JAB52761.1 | *Micrurus fulvius* | 55.04 | 1 | 1 | 0.001% | 15.82 | PLA2 |
| 11 | JAS04986.1 | *Micrurus fulvius* | 55.04 | 1 | 1 | 0.001% | 15.92 | PLA2 |
| 12 | JAS04987.1 | *Micrurus fulvius* | 55.04 | 1 | 1 | 0.001% | 15.91 | PLA2 |

| | | | | | | | | |
|---|---|---|---|---|---|---|---|---|
| 13 | JAS04982.1 | *Micrurus fulvius* | 55.04 | 1 | 1 | 0.001% | 15.92 | PLA2 |
| 14 | JAS04985.1 | *Micrurus fulvius* | 55.04 | 1 | 1 | 0.001% | 15.94 | PLA2 |
| 15 | JAI08996.1 | *Micrurus fulvius* | 55.04 | 1 | 1 | 0.001% | 15.94 | PLA2 |
| 16 | JAI08997.1 | *Micrurus fulvius* | 55.04 | 1 | 1 | 0.001% | 15.92 | PLA2 |
| 17 | JAI08998.1 | *Micrurus fulvius* | 55.04 | 1 | 1 | 0.001% | 15.91 | PLA2 |
| 18 | JAI08993.1 | *Micrurus fulvius* | 55.04 | 1 | 1 | 0.001% | 15.92 | PLA2 |
| 19 | JAB52775.1 | *Micrurus fulvius* | 55.04 | 1 | 1 | 0.001% | 15.92 | PLA2 |
| 20 | JAB52776.1 | *Micrurus fulvius* | 55.04 | 1 | 1 | 0.001% | 15.91 | PLA2 |
| 21 | JAS05103.1 | *Micrurus fulvius* | 55.04 | 1 | 1 | 0.001% | 15.81 | PLA2 |
| 22 | JAS05108.1 | *Micrurus fulvius* | 55.04 | 1 | 1 | 0.001% | 15.72 | PLA2 |
| 23 | JAS05109.1 | *Micrurus fulvius* | 55.04 | 1 | 1 | 0.001% | 15.87 | PLA2 |
| 24 | JAS05107.1 | *Micrurus fulvius* | 55.04 | 1 | 1 | 0.001% | 15.71 | PLA2 |
| 25 | JAS05106.1 | *Micrurus fulvius* | 55.04 | 1 | 1 | 0.001% | 15.77 | PLA2 |
| 26 | JAS05111.1 | *Micrurus fulvius* | 55.04 | 1 | 1 | 0.001% | 15.77 | PLA2 |
| 27 | JAS05101.1 | *Micrurus fulvius* | 55.04 | 1 | 1 | 0.001% | 15.74 | PLA2 |
| 28 | JAS05102.1 | *Micrurus fulvius* | 55.04 | 1 | 1 | 0.001% | 15.78 | PLA2 |
| 29 | JAS05104.1 | *Micrurus fulvius* | 55.04 | 1 | 1 | 0.001% | 15.78 | PLA2 |
| 30 | JAS05105.1 | *Micrurus fulvius* | 55.04 | 1 | 1 | 0.001% | 15.95 | PLA2 |
| 31 | JAS05110.1 | *Micrurus fulvius* | 55.04 | 1 | 1 | 0.001% | 15.77 | PLA2 |
| **Three-finger toxins (3FTx): 13.71%** | | | | | | | | |
| 32 | Q9YGI8.1 | *Bungarus multicinctus* | 200.58 | 5 | 5 | 1.891% | 9.52 | Type I (short) α-neurotoxin |
| 33 | CAA06887.1 | *Bungarus multicinctus multicinctus* | 200.58 | 5 | 5 | 1.891% | 9.52 | Type I (short) α-neurotoxin |
| 34 | P15816.1 | *Bungarus multicinctus* | 297.69 | 10 | 1 | 1.829% | 9.52 | Type II (long) κ-neurotoxin |

| | | | | | | | | |
|---|---|---|---|---|---|---|---|---|
| 35 | CAA35774.1 | *Bungarus multicinctus* | 297.69 | 10 | 1 | 1.829% | 9.52 | Type II (long) κ-neurotoxin |
| 36 | CAA45882 | *Bungarus multicinctus* | 197.44 | 4 | 4 | 0.822% | 8.63 | Type I (short) α-neurotoxin |
| 37 | 1IJC | *Bungarus candidus* | 159.08 | 3 | 3 | 0.680% | 7.29 | Non-conventional neurotoxin |
| 38 | 1F94 | *Bungarus candidus* | 159.08 | 3 | 3 | 0.680% | 7.29 | Non-conventional neurotoxin |
| 39 | P81782.1 | *Bungarus candidus* | 159.08 | 3 | 3 | 0.680% | 7.29 | Non-conventional neurotoxin |
| 40 | CAA06885 | *Bungarus multicinctus multicinctus* | 203.81 | 4 | 4 | 0.621% | 10.63 | Type II (long) κ-neurotoxin |
| 41 | AAL30054.1 | *Bungarus candidus* | 219.26 | 8 | 6 | 0.507% | 9.58 | Type II (long) κ-neurotoxin |
| 42 | Q8AY56.1 | *Bungarus candidus* | 219.26 | 8 | 6 | 0.507% | 9.58 | Type II (long) κ-neurotoxin |
| 43 | Q9W729.1 | *Bungarus multicinctus* | 163.69 | 2 | 2 | 0.378% | 9.66 | Type II (long) κ-neurotoxin |
| 44 | CAB46659.1 | *Bungarus multicinctus* | 163.69 | 2 | 2 | 0.378% | 9.66 | Type II (long) κ-neurotoxin |
| 45 | CAA45882 | *Bungarus multicinctus* | 167.03 | 2 | 2 | 0.275% | 8.57 | Type I (short) α-neurotoxin |
| 46 | AAL30059 | *Bungarus candidus* | 157.51 | 3 | 3 | 0.212% | 9.75 | Non-conventional neurotoxin |
| 47 | CAM11302.1 | *Bungarus caeruleus* | 117.13 | 2 | 2 | 0.048% | 8.31 | Type II (long) α-neurotoxin |
| 48 | D2N116.1 | *Bungarus caeruleus* | 117.13 | 2 | 2 | 0.048% | 8.31 | Type II (long) α-neurotoxin |
| 49 | ACR78492.1 | *Drysdalia coronoides* | 54.81 | 1 | 1 | 0.017% | 9.61 | Type II (long) α-neurotoxin |
| 50 | ACR78484.1 | *Drysdalia coronoides* | 54.81 | 1 | 1 | 0.017% | 9.61 | Type II (long) α-neurotoxin |

| | | | | | | | | |
|---|---|---|---|---|---|---|---|---|
| 51 | F8J2F2.1 | *Drysdalia coronoides* | 54.81 | 1 | 1 | 0.017% | 9.61 | Type II (long) α-neurotoxin |
| 52 | ACR78483.1 | *Drysdalia coronoides* | 54.81 | 1 | 1 | 0.017% | 9.66 | Type II (long) α-neurotoxin |
| 53 | ACR78488.1 | *Drysdalia coronoides* | 54.81 | 1 | 1 | 0.017% | 9.66 | Type II (long) α-neurotoxin |
| 54 | JAA74738.1 | *Denisonia devisi* | 54.81 | 1 | 1 | 0.017% | 10.11 | Type II (long) α-neurotoxin |
| 55 | ACR78486.1 | *Drysdalia coronoides* | 54.81 | 1 | 1 | 0.017% | 9.53 | Type II (long) α-neurotoxin |
| 56 | F8J2E6.1 | *Drysdalia coronoides* | 54.81 | 1 | 1 | 0.017% | 9.53 | Type II (long) α-neurotoxin |
| 57 | ACR78491.1 | *Drysdalia coronoides* | 54.81 | 1 | 1 | 0.017% | 9.50 | Type II (long) α-neurotoxin |
| 58 | JAA74652.1 | *Acanthophis wellsi* | 54.81 | 1 | 1 | 0.017% | 9.64 | Type II (long) α-neurotoxin |
| 59 | JAA74651.1 | *Acanthophis wellsi* | 54.81 | 1 | 1 | 0.017% | 9.64 | Type II (long) α-neurotoxin |
| 60 | JAA74655.1 | *Acanthophis wellsi* | 54.81 | 1 | 1 | 0.017% | 9.57 | Type II (long) α-neurotoxin |
| 61 | ABK63538.1 | *Tropidechis carinatus* | 54.81 | 1 | 1 | 0.017% | 10.14 | Type II (long) α-neurotoxin |
| 62 | A8HDK4.1 | *Tropidechis carinatus* | 54.81 | 1 | 1 | 0.017% | 10.14 | Type II (long) α-neurotoxin |
| 63 | ABX58163.1 | *Austrelaps labialis* | 54.81 | 1 | 1 | 0.017% | 9.59 | Type II (long) α-neurotoxin |
| 64 | ABX58162.1 | *Austrelaps labialis* | 54.81 | 1 | 1 | 0.017% | 9.57 | Type II (long) α-neurotoxin |
| 65 | ABX58156.1 | *Austrelaps labialis* | 54.81 | 1 | 1 | 0.017% | 9.59 | Type II (long) α-neurotoxin |
| 66 | ABX58159.1 | *Austrelaps labialis* | 54.81 | 1 | 1 | 0.017% | 9.57 | Type II (long) α-neurotoxin |

| | | | | | | | | |
|---|---|---|---|---|---|---|---|---|
| 67 | ABX58160.1 | *Austrelaps labialis* | 54.81 | 1 | 1 | 0.017% | 9.65 | Type II (long) α-neurotoxin |
| 68 | ABX58161.1 | *Austrelaps labialis* | 54.81 | 1 | 1 | 0.017% | 9.62 | Type II (long) α-neurotoxin |
| 69 | ABX58158.1 | *Austrelaps labialis* | 54.81 | 1 | 1 | 0.017% | 9.57 | Type II (long) α-neurotoxin |
| 70 | JAA74768.1 | *Echiopsis curta* | 97.53 | 1 | 1 | 0.011% | 11.63 | Type II (long) α-neurotoxin |
| 71 | JAA74767.1 | *Echiopsis curta* | 97.53 | 1 | 1 | 0.011% | 11.63 | Type II (long) α-neurotoxin |
| 72 | P01387.1 | *Ophiophagus hannah* | 97.53 | 1 | 1 | 0.011% | 8.11 | Type II (long) α-neurotoxin |
| 73 | Q53B58.1 | *Ophiophagus hannah* | 97.53 | 1 | 1 | 0.011% | 10.21 | Type II (long) α-neurotoxin |
| 74 | AAT97250.1 | *Ophiophagus hannah* | 97.53 | 1 | 1 | 0.011% | 10.21 | Type II (long) α-neurotoxin |
| 75 | ETE58964 | *Ophiophagus hannah* | 164.12 | 2 | 2 | 0.009% | 9.28 | Type I (short) α-neurotoxin |
| 76 | P01397.2 | *Dendroaspis polylepis polylepis* | 54.53 | 1 | 1 | 0.002% | 7.95 | Type II (long) α-neurotoxin |
| 77 | P25667.1 | *Dendroaspis polylepis polylepis* | 54.53 | 1 | 1 | 0.002% | 7.94 | Type II (long) α-neurotoxin |
| 78 | 1MR6 | *Bungarus multicinctus* | 77.2 | 1 | 1 | 0.002% | 7.54 | Non-conventional neurotoxin |
| 79 | CAA06885.1 | *Bungarus multicinctus multicinctus* | 77.2 | 1 | 1 | 0.002% | 9.83 | Non-conventional neurotoxin |
| 80 | Q9YGJ0.1 | *Bungarus multicinctus* | 77.2 | 1 | 1 | 0.002% | 9.83 | Non-conventional neurotoxin |
| 81 | AAD41806.1 | *Bungarus multicinctus* | 77.2 | 1 | 1 | 0.002% | 9.83 | Non-conventional neurotoxin |
| 82 | CAA72941.1 | *Bungarus multicinctus* | 77.2 | 1 | 1 | 0.002% | 9.76 | Non-conventional neurotoxin |

| | | | | | | | | |
|---|---|---|---|---|---|---|---|---|
| 83 | CAD01082.1 | *Bungarus multicinctus* | 77.2 | 1 | 1 | 0.002% | 9.83 | Non-conventional neurotoxin |
| 84 | O12963.1 | *Bungarus multicinctus* | 77.2 | 1 | 1 | 0.002% | 9.76 | Non-conventional neurotoxin |
| **β-bungarotoxin: 11.61%** | | | | | | | | |
| 85 | AAL87003 | *Bungarus caeruleus* | 361.32 | 24 | 10 | 10.925% | 15.49 | β-bungarotoxin |
| 86 | CAD24466 | *Bungarus multicinctus* | 343.33 | 21 | 3 | 0.683% | 14.90 | β-bungarotoxin |
| 87 | 0402253A | *Bungarus multicinctus* | 256.44 | 10 | 1 | 0.000% | 13.56 | β-bungarotoxin |
| 88 | 0412250A | *Bungarus multicinctus* | 256.44 | 10 | 1 | 0.000% | 13.49 | β-bungarotoxin |
| **Cysteine-rich secretory proteins (CRISP): 9.09%** | | | | | | | | |
| 89 | ACE73577 | *Bungarus candidus* | 429.3 | 34 | 10 | 4.522% | 26.26 | CRISP |
| 90 | ACE73577 | *Bungarus candidus* | 429.3 | 34 | 10 | 4.522% | 31.19 | CRISP |
| 91 | ACE73578.1 | *Macrovipera lebetina* | 377.32 | 19 | 1 | 0.042% | 26.44 | CRISP |
| 92 | ACY68722.1 | *Parasuta nigriceps* | 108.97 | 1 | 1 | 0.001% | 20.02 | CRISP |
| **Kunitz-type serine protease inhibitor (Kunitz): 9.02%** | | | | | | | | |
| 93 | CAP74383.1 | *Bungarus multicinctus* | 224.26 | 4 | 3 | 3.294% | 9.23 | Kunitz |
| 94 | B4ESA4.1 | *Bungarus multicinctus* | 224.26 | 4 | 3 | 3.294% | 9.23 | Kunitz |
| 95 | Q8AY42.2 | *Bungarus candidus* | 157.2 | 3 | 2 | 0.534% | 8.92 | Kunitz |
| 96 | Q8AY43.2 | *Bungarus candidus* | 157.2 | 3 | 2 | 0.534% | 9.08 | Kunitz |
| 97 | AAL30069.1 | *Bungarus candidus* | 157.2 | 3 | 2 | 0.534% | 9.39 | Kunitz |
| 98 | AAL30068.1 | *Bungarus candidus* | 157.2 | 3 | 2 | 0.534% | 9.61 | Kunitz |
| 99 | B4ESA3.1 | *Bungarus multicinctus* | 104.39 | 2 | 1 | 0.074% | 9.16 | Kunitz |
| 100 | Q8AY41.2 | *Bungarus candidus* | 104.39 | 2 | 1 | 0.074% | 9.13 | Kunitz |
| 101 | CAP74382.1 | *Bungarus multicinctus* | 104.39 | 2 | 1 | 0.074% | 9.16 | Kunitz |
| 102 | AAL30070.1 | *Bungarus candidus* | 104.39 | 2 | 1 | 0.074% | 9.66 | Kunitz |

| | | | | | | | | |
|---|---|---|---|---|---|---|---|---|
| **Vascular endothelial growth factor (VEGF): 2.26%** | | | | | | | | |
| 103 | JAA74898 | *Pseudonaja modesta* | 174.15 | 6 | 6 | 0.453% | 22.46 | VEGF |
| 104 | JAA74898 | *Pseudonaja modesta* | 174.15 | 6 | 6 | 0.453% | 22.46 | VEGF |
| 105 | JAA74688 | *Cacophis squamulosus* | 174.15 | 6 | 6 | 0.453% | 24.61 | VEGF |
| 106 | XP_026551344 | *Pseudonaja textilis* | 174.15 | 6 | 6 | 0.453% | 25.39 | VEGF |
| 107 | XP_026551344 | *Pseudonaja textilis* | 174.15 | 6 | 6 | 0.453% | 25.39 | VEGF |
| **Acetylcholinesterase (AChE): 1.12%** | | | | | | | | |
| 108 | Q92035 | *Bungarus fasciatus* | 448.54 | 49 | 3 | 0.157% | 67.89 | AChE |
| 109 | Q92035 | *Bungarus fasciatus* | 448.54 | 49 | 3 | 0.157% | 68.17 | AChE |
| 110 | Q92035 | *Bungarus fasciatus* | 448.54 | 49 | 3 | 0.157% | 68.17 | AChE |
| 111 | Q92035 | *Bungarus fasciatus* | 448.54 | 49 | 3 | 0.157% | 67.34 | AChE |
| 112 | AAC59905 | *Bungarus fasciatus* | 452.57 | 52 | 6 | 0.123% | 64.77 | AChE |
| 113 | AAC59905 | *Bungarus fasciatus* | 452.57 | 52 | 6 | 0.123% | 69.05 | AChE |
| 114 | AAC59905 | *Bungarus fasciatus* | 452.57 | 52 | 6 | 0.123% | 74.79 | AChE |
| 115 | AAC59905 | *Bungarus fasciatus* | 452.57 | 52 | 6 | 0.123% | 77.95 | AChE |
| **Snake venom metalloproteinase (Disintegrin-like): 0.49%** | | | | | | | | |
| 116 | ABN72537.1 | *Bungarus multicinctus* | 304.6 | 21 | 3 | 0.143% | 68.99 | Disintegrin-like |
| 117 | A8QL49.1 | *Bungarus multicinctus* | 304.6 | 21 | 3 | 0.143% | 68.99 | Disintegrin-like |
| 118 | LAA20517 | *Micrurus lemniscatus carvalhoi* | 255.57 | 7 | 6 | 0.110% | 21.44 | Disintegrin-like |
| 119 | AEH95531 | *Drysdalia coronoides* | 227.52 | 7 | 1 | 0.066% | 25.77 | Disintegrin-like |
| 120 | ABN72537 | *Bungarus multicinctus* | 311.2 | 21 | 2 | 0.022% | 68.84 | Disintegrin-like |
| 121 | ABN72536 | *Bungarus fasciatus* | 301.36 | 12 | 2 | 0.009% | 31.50 | Disintegrin-like |

| | | | | | | | | |
|---|---|---|---|---|---|---|---|---|
| **5′-nucleotidase: 0.04%** | | | | | | | | |
| 122 | JAB52815 | *Micrurus fulvius* | 281.36 | 13 | 3 | 0.010% | 64.59 | 5′-nucleotidase |
| 123 | JAB52815 | *Micrurus fulvius* | 281.36 | 13 | 3 | 0.010% | 64.59 | 5′-nucleotidase |
| 124 | JAB52815 | *Micrurus fulvius* | 281.36 | 13 | 3 | 0.010% | 64.59 | 5′-nucleotidase |
| 125 | JAB52815 | *Micrurus fulvius* | 281.36 | 13 | 3 | 0.010% | 64.59 | 5′-nucleotidase |
| 126 | AHJ80886.1 | *Macrovipera lebetina* | 165.14 | 2 | 1 | 0.003% | 45.03 | 5′-nucleotidase |
| **Peptidase: 0.03%** | | | | | | | | |
| 127 | XP_026571757.1 | *Pseudonaja textilis* | 223 | 5 | 5 | 0.027% | 37.55 | Peptidase C1 |
| 128 | XP_034296605.1 | *Pantherophis guttatus* | 209.17 | 6 | 1 | 0.002% | 106.08 | Peptidase M1 |
| 129 | XP_034296606.1 | *Pantherophis guttatus* | 209.17 | 6 | 1 | 0.002% | 106.08 | Peptidase M1 |
| 130 | XP_034296607.1 | *Pantherophis guttatus* | 209.17 | 6 | 1 | 0.002% | 98.37 | Peptidase M1 |
| **Nerve growth factor (NGF): 0.02%** | | | | | | | | |
| 131 | AAB25729.1 | *Bungarus multicinctus* | 104.44 | 1 | 1 | 0.008% | 27.51 | NGF |
| 132 | P34128.1 | *Bungarus multicinctus* | 104.44 | 1 | 1 | 0.008% | 27.51 | NGF |
| **Hyaluronidase: 0.02%** | | | | | | | | |
| 133 | XP_026524834 | *Notechis scutatus* | 287.68 | 16 | 5 | 0.013% | 54.56 | Hyaluronidase |
| 134 | XP_034277309.1 | *Pantherophis guttatus* | 182.52 | 6 | 1 | 0.003% | 54.30 | Hyaluronidase |
| **L-amino-acid oxidase (LAAO): 0.01%** | | | | | | | | |

| | | | | | | | | |
|---|---|---|---|---|---|---|---|---|
| 135 | ABN72539 | *Bungarus multicinctus* | 336.02 | 23 | 2 | 0.004% | 62.87 | LAAO |
| 136 | ABN72539 | *Bungarus multicinctus* | 336.02 | 23 | 2 | 0.004% | 62.87 | LAAO |
| 137 | ABN72540 | *Bungarus fasciatus* | 330.74 | 22 | 1 | 0.001% | 62.83 | LAAO |
| 138 | ABN72540 | *Bungarus fasciatus* | 330.74 | 22 | 1 | 0.001% | 62.83 | LAAO |
| **Phospholipase B (PLB): 0.005%** | | | | | | | | |
| 139 | ACR78473 | *Drysdalia coronoides* | 192.46 | 6 | 6 | 0.005% | 64.07 | PLB |
| **Snake venom serine protease: 0.004%** | | | | | | | | |
| 140 | ABN72545 | *Bungarus multicinctus* | 161.71 | 4 | 2 | 0.002% | 31.78 | Serine Protease |
| 141 | ABN72545.1 | *Bungarus multicinctus* | 158.73 | 3 | 1 | 0.001% | 31.01 | Serine Protease |
| 142 | A8QL57.1 | *Bungarus multicinctus* | 158.73 | 3 | 1 | 0.001% | 31.01 | Serine Protease |
| **C-type lectin (CTL): 0.003%** | | | | | | | | |
| 143 | ABH05181.1 | *Bungarus multicinctus* | 128.06 | 1 | 1 | 0.002% | 18.27 | CTL |
| 144 | A1XXJ9.1 | *Bungarus multicinctus* | 128.06 | 1 | 1 | 0.002% | 18.27 | CTL |
| **Vespryn: 0.002%** | | | | | | | | |
| 145 | XP_026576508 | *Pseudonaja textilis* | 118.47 | 2 | 2 | 0.002% | 24.38 | Vespryn |
| **Phosphodiesterase (PDE): 0.002%** | | | | | | | | |
| 146 | XP_026521896.1 | *Notechis scutatus* | 199.72 | 4 | 1 | 0.001% | 96.90 | PDE |
| 147 | XP_026521895.1 | *Notechis scutatus* | 199.72 | 4 | 1 | 0.001% | 100.96 | PDE |

**S4B Table.** The proteomic composition of *B. caeruleus* venom.

| Sr. No. | Accession | Species | -10lgP | #Peptides | #Unique | Relative abundance of toxin hit (%) | Avg. Mass (KDa) | Family |
|---|---|---|---|---|---|---|---|---|
| **Three-finger toxins (3FTx): 65.54%** | | | | | | | | |
| 1 | 2CTX | *Naja naja* | 158.07 | 3 | 2 | 9.260% | 7.83 | Type II (long) α-neurotoxin |
| 2 | 1LXH | *Naja kaouthia* | 158.07 | 3 | 2 | 9.260% | 7.83 | Type II (long) α-neurotoxin |
| 3 | 1LXG | *Naja kaouthia* | 158.07 | 3 | 2 | 9.260% | 7.83 | Type II (long) α-neurotoxin |
| 4 | P01391.1 | *Naja kaouthia* | 158.07 | 3 | 2 | 9.260% | 7.83 | Type II (long) α-neurotoxin |
| 5 | 4AEA | *Naja kaouthia* | 158.07 | 3 | 2 | 9.260% | 7.83 | Type II (long) α-neurotoxin |
| 6 | 1CTX | *Naja siamensis* | 158.07 | 3 | 2 | 9.260% | 7.83 | Type II (long) α-neurotoxin |
| 7 | 1YI5 | *Naja siamensis* | 158.07 | 3 | 2 | 9.260% | 7.83 | Type II (long) α-neurotoxin |
| 8 | P86538.2 | *Naja naja* | 185.04 | 3 | 3 | 0.134% | 6.71 | Cytotoxin |
| 9 | BAU24666.1 | *Naja naja* | 185.04 | 3 | 3 | 0.134% | 7.96 | Cytotoxin |

| | | | | | | | | |
|---|---|---|---|---|---|---|---|---|
| 10 | LAB55960 | *Micrurus surinamensis* | 134.85 | 3 | 3 | 0.120% | 10.09 | Non-conventiona l neurotoxin |
| 11 | 1IJC | *Bungarus candidus* | 80.02 | 1 | 1 | 0.038% | 7.29 | Non-conventiona l neurotoxin |
| 12 | 1F94 | *Bungarus candidus* | 80.02 | 1 | 1 | 0.038% | 7.29 | Non-conventiona l neurotoxin |
| 13 | P81782.1 | *Bungarus candidus* | 80.02 | 1 | 1 | 0.038% | 7.29 | Non-conventiona l neurotoxin |
| 14 | 5MG9 | *Dendroaspis angusticeps* | 82.53 | 1 | 1 | 0.026% | 7.24 | Type I (short) aminergic toxin |
| 15 | P82463.1 | *Naja kaouthia* | 82.53 | 1 | 1 | 0.026% | 7.30 | Type C muscarinic toxin |
| 16 | Q9W717.1 | *Naja atra* | 89.64 | 1 | 1 | 0.025% | 9.70 | Type I (short) orphan neurotoxin |
| 17 | APB88857.1 | *Naja naja* | 89.64 | 1 | 1 | 0.025% | 9.70 | Type I (short) orphan neurotoxin |
| 18 | CAB45156.1 | *Naja naja* | 89.64 | 1 | 1 | 0.025% | 9.70 | Type I (short) orphan neurotoxin |
| 19 | 764177A | *Naja naja* | 118.46 | 2 | 1 | 0.016% | 7.71 | Non-conventiona l neurotoxin |
| 20 | P25669.1 | P25669.1 | 118.46 | 2 | 1 | 0.016% | 7.82 | Type II (long) α-neurotoxin |
| 21 | AAL30061.1 | *Bungarus candidus* | 51.91 | 1 | 1 | 0.004% | 9.76 | Non-conventiona l neurotoxin |
| 22 | Q8AY49.1 | *Bungarus candidus* | 51.91 | 1 | 1 | 0.004% | 9.76 | Non-conventiona l neurotoxin |
| 23 | Q8AY51.1 | *Bungarus candidus* | 51.91 | 1 | 1 | 0.004% | 9.71 | Non-conventiona l neurotoxin |
| 24 | AAL30059.1 | *Bungarus candidus* | 51.91 | 1 | 1 | 0.004% | 9.71 | Non-conventiona l neurotoxin |

| | | | | | | | | |
|---|---|---|---|---|---|---|---|---|
| 25 | A2CKF7.1 | *Bungarus fasciatus* | 51.91 | 1 | 1 | 0.004% | 9.72 | Non-conventional neurotoxin |
| 26 | CAA11133.1 | *Naja atra* | 51.91 | 1 | 1 | 0.004% | 9.85 | Non-conventional neurotoxin |
| 27 | ADF50021.1 | *Bungarus flaviceps* | 51.91 | 1 | 1 | 0.004% | 9.74 | Non-conventional neurotoxin |
| 28 | ABI33871.1 | *Bungarus fasciatus* | 51.91 | 1 | 1 | 0.004% | 9.72 | Non-conventional neurotoxin |
| 29 | AAT38875.1 | *Bungarus candidus* | 51.91 | 1 | 1 | 0.004% | 9.72 | Non-conventional neurotoxin |
| 30 | AAL30060.1 | *Bungarus candidus* | 51.91 | 1 | 1 | 0.004% | 9.65 | Non-conventional neurotoxin |
| 31 | Q8AY50.1 | *Bungarus candidus* | 51.91 | 1 | 1 | 0.004% | 9.65 | Non-conventional neurotoxin |
| 32 | Q9YGI2.1 | *Naja atra* | 51.91 | 1 | 1 | 0.004% | 9.85 | Non-conventional neurotoxin |
| 33 | ADF50022.1 | *Bungarus flaviceps* | 51.91 | 1 | 1 | 0.004% | 9.76 | Non-conventional neurotoxin |
| 34 | ADF50020.1 | *Bungarus flaviceps* | 51.91 | 1 | 1 | 0.004% | 9.68 | Type I (short) α-neurotoxin |
| 35 | ADN67577.1 | *Naja atra* | 51.91 | 1 | 1 | 0.004% | 9.86 | Non-conventional neurotoxin |
| 36 | CAA11134.1 | *Naja atra* | 51.91 | 1 | 1 | 0.004% | 9.74 | Non-conventional neurotoxin |
| 37 | Q6IZ95.1 | *Bungarus candidus* | 51.91 | 1 | 1 | 0.004% | 9.72 | Non-conventional neurotoxin |
| 38 | Q9YGI1.1 | *Naja atra* | 51.91 | 1 | 1 | 0.004% | 9.74 | Non-conventional neurotoxin |
| **Phospholipase A2 ($PLA_2$): 10.5%** | | | | | | | | |
| 39 | 1U4J | *Bungarus caeruleus* | 194.87 | 5 | 1 | 1.340% | 12.97 | PLA2 |
| 40 | 1G0Z | *Bungarus caeruleus* | 194.87 | 5 | 1 | 1.340% | 12.97 | PLA2 |

| | | | | | | | | |
|---|---|---|---|---|---|---|---|---|
| 41 | Q6SLM1.1 | *Bungarus caeruleus* | 194.87 | 5 | 1 | 1.340% | 14.83 | PLA2 |
| 42 | AAR19228.1 | *Bungarus caeruleus* | 194.87 | 5 | 1 | 1.340% | 14.83 | PLA2 |
| 43 | 1PSH | *Naja naja* | 167.87 | 4 | 3 | 0.523% | 13.35 | PLA2 |
| 44 | 1A3F | *Naja naja* | 167.87 | 4 | 3 | 0.523% | 13.35 | PLA2 |
| 45 | 1A3D | *Naja naja* | 167.87 | 4 | 3 | 0.523% | 13.35 | PLA2 |
| 46 | P15445.1 | *Naja naja* | 167.87 | 4 | 3 | 0.523% | 13.35 | PLA2 |
| 47 | CAA45372.1 | *Naja naja* | 167.87 | 4 | 3 | 0.523% | 13.48 | PLA2 |
| 48 | 0702209A | *Bungarus multicinctus* | 120.69 | 2 | 2 | 0.082% | 12.83 | PLA2 |
| 49 | P00606.2 | *Bungarus multicinctus* | 120.69 | 2 | 2 | 0.082% | 15.59 | PLA2 |
| 50 | CAA37482.1 | *Bungarus multicinctus* | 120.69 | 2 | 2 | 0.082% | 15.59 | PLA2 |
| 51 | JAB52792.1 | *Micrurus fulvius* | 190.96 | 5 | 1 | 0.077% | 16.16 | PLA2 |
| 52 | AAZ29513.1 | *Micrurus fulvius* | 190.96 | 5 | 1 | 0.077% | 15.33 | PLA2 |
| 53 | JAB52790.1 | *Micrurus fulvius* | 190.96 | 5 | 1 | 0.077% | 15.98 | PLA2 |
| 54 | JAB52794.1 | *Micrurus fulvius* | 190.96 | 5 | 1 | 0.077% | 16.03 | PLA2 |
| 55 | JAB52809.1 | *Micrurus fulvius* | 190.96 | 5 | 1 | 0.077% | 16.08 | PLA2 |
| 56 | JAB52771.1 | *Micrurus fulvius* | 190.96 | 5 | 1 | 0.077% | 16.21 | PLA2 |
| 57 | JAS04998.1 | *Micrurus fulvius* | 190.96 | 5 | 1 | 0.077% | 16.16 | PLA2 |
| 58 | JAB52773.1 | *Micrurus fulvius* | 190.96 | 5 | 1 | 0.077% | 16.22 | PLA2 |
| 59 | JAB52772.1 | *Micrurus fulvius* | 190.96 | 5 | 1 | 0.077% | 16.19 | PLA2 |
| 60 | JAS04995.1 | *Micrurus fulvius* | 190.96 | 5 | 1 | 0.077% | 16.18 | PLA2 |
| 61 | JAB52770.1 | *Micrurus fulvius* | 190.96 | 5 | 1 | 0.077% | 16.16 | PLA2 |
| 62 | JAB52795.1 | *Micrurus fulvius* | 190.96 | 5 | 1 | 0.077% | 16.24 | PLA2 |
| 63 | JAB52774.1 | *Micrurus fulvius* | 190.96 | 5 | 1 | 0.077% | 16.18 | PLA2 |
| 64 | JAS04999.1 | *Micrurus fulvius* | 190.96 | 5 | 1 | 0.077% | 16.18 | PLA2 |
| 65 | JAI09010.1 | *Micrurus fulvius* | 190.96 | 5 | 1 | 0.077% | 16.18 | PLA2 |
| 66 | JAI09006.1 | *Micrurus fulvius* | 190.96 | 5 | 1 | 0.077% | 16.18 | PLA2 |
| 67 | JAI09004.1 | *Micrurus fulvius* | 190.96 | 5 | 1 | 0.077% | 16.30 | PLA2 |

| | | | | | | | | |
|---|---|---|---|---|---|---|---|---|
| 68 | JAI09005.1 | *Micrurus fulvius* | 190.96 | 5 | 1 | 0.077% | 16.24 | PLA2 |
| 69 | JAI09009.1 | *Micrurus fulvius* | 190.96 | 5 | 1 | 0.077% | 16.16 | PLA2 |
| 70 | JAS04993.1 | *Micrurus fulvius* | 190.96 | 5 | 1 | 0.077% | 16.30 | PLA2 |
| 71 | JAB52804.1 | *Micrurus fulvius* | 190.96 | 5 | 1 | 0.077% | 16.29 | PLA2 |
| 72 | JAS04994.1 | *Micrurus fulvius* | 190.96 | 5 | 1 | 0.077% | 16.24 | PLA2 |
| 73 | JAB52764.1 | *Micrurus fulvius* | 190.96 | 5 | 1 | 0.077% | 16.16 | PLA2 |
| 74 | LAA98189.1 | *Micrurus lemniscatus lemniscatus* | 190.96 | 5 | 1 | 0.077% | 11.26 | PLA2 |
| 75 | JAS04997.1 | *Micrurus fulvius* | 190.96 | 5 | 1 | 0.077% | 16.16 | PLA2 |
| 76 | JAI09008.1 | *Micrurus fulvius* | 190.96 | 5 | 1 | 0.077% | 16.16 | PLA2 |
| 77 | ABU68503.1 | *Trimorphodon biscutatus* | 190.96 | 5 | 1 | 0.077% | 16.62 | PLA2 |
| 78 | ABU63164.1 | *Bungarus fasciatus* | 192.65 | 5 | 1 | 0.034% | 15.95 | PLA2 |
| 79 | ABU63166.1 | *Bungarus fasciatus* | 192.65 | 5 | 1 | 0.034% | 16.00 | PLA2 |
| 80 | AAO84769.1 | *Bungarus candidus* | 115.37 | 2 | 2 | 0.032% | 15.64 | PLA2 |
| 81 | Q802I1.1 | *Bungarus candidus* | 115.37 | 2 | 2 | 0.032% | 15.64 | PLA2 |
| 82 | AAL54920.1 | *Hydrophis hardwickii* | 60.95 | 1 | 1 | 0.028% | 16.11 | PLA2 |
| 83 | ABN54807.1 | *Hydrophis hardwickii* | 60.95 | 1 | 1 | 0.028% | 16.08 | PLA2 |
| **β-bungarotoxin: 7.1%** | | | | | | | | |
| 84 | Q8QFW3.1 | *Bungarus caeruleus* | 237.26 | 6 | 5 | 3.246% | 16.12 | β-bungarotoxin |
| 85 | AAL87004.1 | *Bungarus caeruleus* | 237.26 | 6 | 5 | 3.246% | 16.12 | β-bungarotoxin |
| 86 | Q8QFW4.1 | *Bungarus caeruleus* | 204.23 | 4 | 3 | 0.302% | 16.36 | β-bungarotoxin |
| 87 | AAL87003.1 | *Bungarus caeruleus* | 204.23 | 4 | 3 | 0.302% | 16.36 | β-bungarotoxin |
| **Acetylcholinesterase (AChE): 5.24%** | | | | | | | | |
| 88 | Q92035 | *Bungarus fasciatus* | 432.47 | 40 | 18 | 5.092% | 68.17 | AChE |
| 89 | S68801 | *Bungarus fasciatus* | 221.06 | 5 | 1 | 0.153% | 9.79 | AChE |

| | | | | | | | | |
|---|---|---|---|---|---|---|---|---|
| **L-amino-acid oxidase (LAAO): 3.96%** | | | | | | | | |
| 90 | 5Z2G | *Naja atra* | 381.05 | 22 | 18 | 1.085% | 57.96 | LAAO |
| 91 | AVX27607.1 | *Naja atra* | 381.05 | 22 | 18 | 1.085% | 57.96 | LAAO |
| 92 | A8QL58.2 | *Naja atra* | 381.05 | 22 | 18 | 1.085% | 57.96 | LAAO |
| 93 | ABN72539.1 | *Bungarus multicinctus* | 325.23 | 17 | 5 | 0.235% | 58.81 | LAAO |
| 94 | A8QL51.1 | *Bungarus multicinctus* | 325.23 | 17 | 5 | 0.235% | 58.81 | LAAO |
| 95 | ABN72539 | *Bungarus multicinctus* | 308.62 | 15 | 2 | 0.065% | 58.92 | LAAO |
| 96 | ABN72539 | *Bungarus multicinctus* | 308.62 | 15 | 2 | 0.065% | 66.29 | LAAO |
| 97 | P0DI91.1 | *Naja oxiana* | 203.51 | 4 | 2 | 0.049% | 11.22 | LAAO |
| 98 | AXL95288.1 | *Spilotes sulphureus* | 222.07 | 6 | 1 | 0.016% | 58.59 | LAAO |
| 99 | AXL95287.1 | *Spilotes sulphureus* | 222.07 | 6 | 1 | 0.016% | 58.59 | LAAO |
| 100 | JAA74761.1 | *Echiopsis curta* | 241.21 | 9 | 1 | 0.009% | 59.21 | LAAO |
| 101 | LAA54621.1 | *Micrurus corallinus* | 228.7 | 7 | 1 | 0.008% | 58.75 | LAAO |
| 102 | JAC94991.1 | *Opheodrys aestivus* | 125.19 | 3 | 1 | 0.002% | 40.49 | LAAO |
| **Cobra venom factor (CVF): 3.14%** | | | | | | | | |
| 103 | 6I2X | *Naja kaouthia* | 387.98 | 23 | 15 | 0.449% | 184.52 | CVF |
| 104 | 3PRX | *Naja kaouthia* | 387.98 | 23 | 15 | 0.449% | 184.52 | CVF |
| 105 | I51018 | *Naja kaouthia* | 387.98 | 23 | 15 | 0.449% | 184.52 | CVF |
| 106 | 3PVM | *Naja kaouthia* | 387.98 | 23 | 15 | 0.449% | 184.52 | CVF |
| 107 | AAA68989.1 | *Naja kaouthia* | 387.98 | 23 | 15 | 0.449% | 184.52 | CVF |
| 108 | Q91132.1 | *Naja kaouthia* | 387.98 | 23 | 15 | 0.449% | 184.52 | CVF |
| 109 | CAI46845.1 | *Naja naja* | 387.98 | 23 | 15 | 0.449% | 184.52 | CVF |
| **Snake venom metalloproteinase (Disintegrin-like): 2.47%** | | | | | | | | |

| | | | | | | | | |
|---|---|---|---|---|---|---|---|---|
| 110 | ABN72537 | *Bungarus multicinctus* | 273.37 | 8 | 5 | 0.648% | 68.85 | Disintegrin-like |
| 111 | ABN72537 | *Bungarus multicinctus* | 273.37 | 8 | 5 | 0.648% | 72.61 | Disintegrin-like |
| 112 | ABN72537.1 | *Bungarus multicinctus* | 294.11 | 13 | 7 | 0.229% | 68.99 | Disintegrin-like |
| 113 | A8QL49.1 | *Bungarus multicinctus* | 294.11 | 13 | 7 | 0.229% | 68.99 | Disintegrin-like |
| 114 | P82942.1 | *Naja kaouthia* | 262.78 | 11 | 7 | 0.172% | 44.49 | Disintegrin-like |
| 115 | Q9PVK7.1 | *Naja kaouthia* | 228.65 | 7 | 3 | 0.093% | 67.66 | Disintegrin-like |
| 116 | AAF00693.1 | *Naja naja* | 228.65 | 7 | 3 | 0.093% | 67.66 | Disintegrin-like |
| 117 | ABN72537 | *Bungarus multicinctus* | 254.45 | 8 | 2 | 0.062% | 50.60 | Disintegrin-like |
| 118 | XP_013922279.1 | *Thamnophis sirtalis* | 56.85 | 1 | 1 | 0.028% | 18.22 | Disintegrin-like |
| 119 | XP_032084681.1 | *Thamnophis elegans* | 56.85 | 1 | 1 | 0.028% | 69.29 | Disintegrin-like |
| 120 | ADD14036.1 | *Naja atra* | 244.4 | 7 | 2 | 0.022% | 44.75 | Disintegrin-like |
| 121 | D6PXE8.1 | *Naja atra* | 244.4 | 7 | 2 | 0.022% | 66.25 | Disintegrin-like |
| 122 | ADG02948.1 | *Naja atra* | 244.4 | 7 | 2 | 0.022% | 66.25 | Disintegrin-like |
| 123 | 3K7N | *Naja atra* | 244.4 | 7 | 2 | 0.022% | 44.19 | Disintegrin-like |
| 124 | ACN50005.1 | *Naja atra* | 244.4 | 7 | 2 | 0.022% | 66.29 | Disintegrin-like |
| 125 | D3TTC1.1 | *Naja atra* | 244.4 | 7 | 2 | 0.022% | 66.29 | Disintegrin-like |
| 126 | D5LMJ3.1 | *Naja atra* | 116.03 | 2 | 2 | 0.017% | 68.25 | Disintegrin-like |
| 127 | ADF43026.1 | *Naja atra* | 116.03 | 2 | 2 | 0.017% | 68.25 | Disintegrin-like |
| 128 | JAC96437.1 | *Hypsiglena sp. JMG-2014* | 79.39 | 1 | 1 | 0.004% | 68.09 | Disintegrin-like |
| 129 | JAS04919.1 | *Hypsiglena sp. JMG-2014* | 79.39 | 1 | 1 | 0.004% | 68.09 | Disintegrin-like |
| 130 | JAS04923.1 | *Hypsiglena sp. JMG-2014* | 79.39 | 1 | 1 | 0.004% | 68.47 | Disintegrin-like |
| 131 | JAC96439.1 | *Hypsiglena sp. JMG-2014* | 79.39 | 1 | 1 | 0.004% | 68.51 | Disintegrin-like |
| 132 | JAS04921.1 | *Hypsiglena sp. JMG-2014* | 79.39 | 1 | 1 | 0.004% | 68.51 | Disintegrin-like |
| 133 | JAS04922.1 | *Hypsiglena sp. JMG-2014* | 79.39 | 1 | 1 | 0.004% | 68.49 | Disintegrin-like |
| 134 | JAC96441.1 | *Hypsiglena sp. JMG-2014* | 79.39 | 1 | 1 | 0.004% | 68.47 | Disintegrin-like |
| 135 | JAC96440.1 | *Hypsiglena sp. JMG-2014* | 79.39 | 1 | 1 | 0.004% | 68.49 | Disintegrin-like |
| 136 | AXL96635.1 | *Ahaetulla prasina* | 148.44 | 4 | 1 | 0.003% | 68.84 | Disintegrin-like |

| | | | | | | | | |
|---|---|---|---|---|---|---|---|---|
| 137 | AXL96667.1 | *Ahaetulla prasina* | 148.44 | 4 | 1 | 0.003% | 68.81 | Disintegrin-like |
| 138 | AXL96646.1 | *Ahaetulla prasina* | 148.44 | 4 | 1 | 0.003% | 68.78 | Disintegrin-like |
| 139 | AXL96640.1 | *Ahaetulla prasina* | 148.44 | 4 | 1 | 0.003% | 68.85 | Disintegrin-like |
| 140 | AXL96666.1 | *Ahaetulla prasina* | 148.44 | 4 | 1 | 0.003% | 68.87 | Disintegrin-like |
| 141 | AXL96626.1 | *Ahaetulla prasina* | 148.44 | 4 | 1 | 0.003% | 72.96 | Disintegrin-like |
| 142 | AXL96631.1 | *Ahaetulla prasina* | 148.44 | 4 | 1 | 0.003% | 67.77 | Disintegrin-like |
| 143 | AXL96650.1 | *Ahaetulla prasina* | 148.44 | 4 | 1 | 0.003% | 71.88 | Disintegrin-like |
| 144 | AXL96645.1 | *Ahaetulla prasina* | 148.44 | 4 | 1 | 0.003% | 71.45 | Disintegrin-like |
| 145 | AXL96653.1 | *Ahaetulla prasina* | 148.44 | 4 | 1 | 0.003% | 68.11 | Disintegrin-like |
| 146 | AXL96648.1 | *Ahaetulla prasina* | 148.44 | 4 | 1 | 0.003% | 68.24 | Disintegrin-like |
| 147 | AXL96655.1 | *Ahaetulla prasina* | 148.44 | 4 | 1 | 0.003% | 68.22 | Disintegrin-like |
| 148 | AXL96638.1 | *Ahaetulla prasina* | 148.44 | 4 | 1 | 0.003% | 68.15 | Disintegrin-like |
| 149 | AXL96643.1 | *Ahaetulla prasina* | 148.44 | 4 | 1 | 0.003% | 68.28 | Disintegrin-like |
| 150 | AXL96644.1 | *Ahaetulla prasina* | 148.44 | 4 | 1 | 0.003% | 68.22 | Disintegrin-like |
| 151 | AXL96633.1 | *Ahaetulla prasina* | 148.44 | 4 | 1 | 0.003% | 68.17 | Disintegrin-like |
| **Peptidase S1: 0.62%** | | | | | | | | |
| 152 | XP_034291085.1 | *Pantherophis guttatus* | 222.57 | 7 | 3 | 0.144% | 63.13 | Peptidase S1 |
| 153 | XP_034291087.1 | *Pantherophis guttatus* | 222.57 | 7 | 3 | 0.144% | 63.13 | Peptidase S1 |
| 154 | XP_034291086.1 | *Pantherophis guttatus* | 222.57 | 7 | 3 | 0.144% | 63.13 | Peptidase S1 |
| 155 | XP_026544671.1 | *Notechis scutatus* | 202.44 | 5 | 1 | 0.078% | 60.66 | Peptidase S1 |
| 156 | XP_026570965.1 | *Pseudonaja textilis* | 202.44 | 5 | 1 | 0.078% | 64.11 | Peptidase S1 |
| 157 | ETE66745.1 | *Ophiophagus hannah* | 144.06 | 2 | 1 | 0.026% | 24.25 | Peptidase S1 |
| **Vascular endothelial growth factor (VEGF): 0.35%** | | | | | | | | |
| 158 | XP_026551346 | *Pseudonaja textilis* | 126.27 | 2 | 2 | 0.056% | 16.57 | VEGF |

| | | | | | | | | |
|---|---|---|---|---|---|---|---|---|
| 159 | XP_026551346 | *Pseudonaja textilis* | 126.27 | 2 | 2 | 0.056% | 16.57 | VEGF |
| 160 | XP_026551346 | *Pseudonaja textilis* | 126.27 | 2 | 2 | 0.056% | 16.57 | VEGF |
| 161 | XP_026551346 | *Pseudonaja textilis* | 126.27 | 2 | 2 | 0.056% | 17.34 | VEGF |
| 162 | JAA74898 | *Pseudonaja modesta* | 126.27 | 2 | 2 | 0.056% | 22.46 | VEGF |
| 163 | XP_026551344 | *Pseudonaja textilis* | 126.27 | 2 | 2 | 0.056% | 25.39 | VEGF |
| 164 | XP_007425787.1 | *Python bivittatus* | 50.53 | 1 | 1 | 0.001% | 42.64 | VEGF |
| 165 | XP_034298589.1 | *Pantherophis guttatus* | 50.53 | 1 | 1 | 0.001% | 42.84 | VEGF |
| 166 | JAC96561.1 | *Echis coloratus* | 50.53 | 1 | 1 | 0.001% | 47.65 | VEGF |
| 167 | JAI10614.1 | *Crotalus adamanteus* | 50.53 | 1 | 1 | 0.001% | 47.70 | VEGF |
| 168 | SMD28268.1 | *Crotalus durissus terrificus* | 50.53 | 1 | 1 | 0.001% | 47.61 | VEGF |
| 169 | JAC95037.1 | *Pantherophis guttatus* | 50.53 | 1 | 1 | 0.001% | 47.51 | VEGF |
| 170 | JAV48556.1 | *Agkistrodon contortrix contortrix* | 50.53 | 1 | 1 | 0.001% | 47.65 | VEGF |
| 171 | JAA95033.1 | *Crotalus horridus* | 50.53 | 1 | 1 | 0.001% | 47.64 | VEGF |
| 172 | XP_034298588.1 | *Pantherophis guttatus* | 50.53 | 1 | 1 | 0.001% | 47.51 | VEGF |
| 173 | XP_013912093.1 | *Thamnophis sirtalis* | 50.53 | 1 | 1 | 0.001% | 47.48 | VEGF |
| 174 | ASU45066.1 | *Daboia russelii* | 50.53 | 1 | 1 | 0.001% | 47.70 | VEGF |
| 175 | JAC94939.1 | *Python regius* | 50.53 | 1 | 1 | 0.001% | 47.54 | VEGF |
| 176 | XP_029139565.1 | *Protobothrops mucrosquamatus* | 50.53 | 1 | 1 | 0.001% | 47.71 | VEGF |
| 177 | JAC94973.1 | *Opheodrys aestivus* | 50.53 | 1 | 1 | 0.001% | 40.06 | VEGF |
| 178 | XP_034298590.1 | *Pantherophis guttatus* | 50.53 | 1 | 1 | 0.001% | 40.06 | VEGF |
| 179 | JAI08988.1 | *Micrurus fulvius* | 50.53 | 1 | 1 | 0.001% | 47.56 | VEGF |
| 180 | ETE60014.1 | *Ophiophagus hannah* | 50.53 | 1 | 1 | 0.001% | 47.56 | VEGF |
| 181 | JAB52939.1 | *Micrurus fulvius* | 50.53 | 1 | 1 | 0.001% | 47.56 | VEGF |
| 182 | LAB17921 | *Micrurus spixii* | 50.53 | 1 | 1 | 0.001% | 47.59 | VEGF |
| 183 | JAS04977.1 | *Micrurus fulvius* | 50.53 | 1 | 1 | 0.001% | 47.56 | VEGF |

| | | | | | | | | |
|---|---|---|---|---|---|---|---|---|
| 184 | XP_026522979.1 | *Notechis scutatus* | 50.53 | 1 | 1 | 0.001% | 47.53 | VEGF |
| 185 | XP_026555459.1 | *Pseudonaja textilis* | 50.53 | 1 | 1 | 0.001% | 47.62 | VEGF |
| 186 | JAI09156.1 | *Micrurus fulvius* | 50.53 | 1 | 1 | 0.001% | 47.56 | VEGF |
| 187 | LAA17506.1 | *Micrurus lemniscatus carvalhoi* | 50.53 | 1 | 1 | 0.001% | 53.37 | VEGF |
| 188 | LAB17921.1 | *Micrurus spixii* | 50.53 | 1 | 1 | 0.001% | 53.46 | VEGF |
| 189 | XP_026555459 | *Pseudonaja textilis* | 50.53 | 1 | 1 | 0.001% | 32.63 | VEGF |
| 190 | LAB17920.1 | *Micrurus spixii* | 50.53 | 1 | 1 | 0.001% | 32.58 | VEGF |
| 191 | LAB17921 | *Micrurus spixii* | 50.53 | 1 | 1 | 0.001% | 40.13 | VEGF |
| **Cysteine-rich secretory proteins (CRISP): 0.32%** | | | | | | | | |
| 192 | ACE73577 | *Bungarus candidus* | 151.6 | 2 | 2 | 0.159% | 26.29 | CRISP |
| 193 | ACE73577 | *Bungarus candidus* | 151.6 | 2 | 2 | 0.159% | 26.29 | CRISP |
| **5′-nucleotidase: 0.19%** | | | | | | | | |
| 194 | JAS05143.1 | *Micrurus tener* | 290.68 | 14 | 4 | 0.021% | 63.01 | 5'-nucleotidase |
| 195 | JAI09048.1 | *Micrurus fulvius* | 290.68 | 14 | 4 | 0.021% | 63.03 | 5'-nucleotidase |
| 196 | JAS05036.1 | *Micrurus fulvius* | 290.68 | 14 | 4 | 0.021% | 62.98 | 5'-nucleotidase |
| 197 | JAI09049.1 | *Micrurus fulvius* | 290.68 | 14 | 4 | 0.021% | 63.03 | 5'-nucleotidase |
| 198 | JAS05038.1 | *Micrurus fulvius* | 290.68 | 14 | 4 | 0.021% | 63.03 | 5'-nucleotidase |
| 199 | JAS05037.1 | *Micrurus fulvius* | 290.68 | 14 | 4 | 0.021% | 63.03 | 5'-nucleotidase |
| 200 | JAI09047.1 | *Micrurus fulvius* | 290.68 | 14 | 4 | 0.021% | 62.98 | 5'-nucleotidase |
| 201 | JAB52815.1 | *Micrurus fulvius* | 290.68 | 14 | 4 | 0.021% | 62.98 | 5'-nucleotidase |
| 202 | XP_026581084 | *Pseudonaja textilis* | 143.89 | 3 | 1 | 0.010% | 17.09 | 5'-nucleotidase |
| 203 | AXL95273.1 | *Spilotes sulphureus* | 164.25 | 5 | 1 | 0.003% | 64.54 | 5'-nucleotidase |
| 204 | JAG67188.1 | *Boiga irregularis* | 164.25 | 5 | 1 | 0.003% | 64.76 | 5'-nucleotidase |
| 205 | B6EWW8.1 | *Gloydius brevicaudus* | 236.58 | 9 | 1 | 0.001% | 64.43 | 5'-nucleotidase |

| | | | | | | | | |
|---|---|---|---|---|---|---|---|---|
| 206 | BAG82602.1 | *Gloydius brevicaudus* | 236.58 | 9 | 1 | 0.001% | 64.43 | 5'-nucleotidase |
| **Snake venom serine protease: 0.18%** | | | | | | | | |
| 207 | A8QL56.1 | *Ophiophagus hannah* | 144.06 | 2 | 1 | 0.026% | 28.66 | Serine Protease |
| 208 | ABN72544.1 | *Ophiophagus hannah* | 144.06 | 2 | 1 | 0.026% | 28.66 | Serine Protease |
| 209 | ABI74694.1 | *Philodryas olfersii* | 144.06 | 2 | 1 | 0.026% | 28.45 | Serine Protease |
| 210 | Q09GK1.1 | *Philodryas olfersii* | 144.06 | 2 | 1 | 0.026% | 28.45 | Serine Protease |
| 211 | AJB84504.1 | *Philodryas chamissonis* | 144.06 | 2 | 1 | 0.026% | 28.57 | Serine Protease |
| 212 | ABN72545.1 | *Bungarus multicinctus* | 184.12 | 4 | 3 | 0.024% | 31.01 | Serine Protease |
| 213 | A8QL57.1 | *Bungarus multicinctus* | 184.12 | 4 | 3 | 0.024% | 31.01 | Serine Protease |
| **Kunitz-type serine protease inhibitor (Kunitz): 0.16%** | | | | | | | | |
| 214 | 1702215A | *Naja naja* | 93.85 | 2 | 2 | 0.067% | 6.51 | Kunitz |
| 215 | P19859.1 | *Naja naja* | 93.85 | 2 | 2 | 0.067% | 6.51 | Kunitz |
| 216 | ABN72545 | *Bungarus multicinctus* | 184.12 | 4 | 3 | 0.024% | 31.55 | Kunitz |
| **Vespryn: 0.12%** | | | | | | | | |
| 217 | P82885.1 | *Naja kaouthia* | 153.4 | 3 | 3 | 0.117% | 12.04 | Vespryn |
| **Phosphodiesterase (PDE): 0.1%** | | | | | | | | |
| 218 | 5GZ4 | *Naja atra* | 241.28 | 8 | 8 | 0.034% | 94.62 | PDE |
| 219 | 5GZ5 | *Naja atra* | 241.28 | 8 | 8 | 0.034% | 94.62 | PDE |
| 220 | A0A2D0TC04.1 | *Naja atra* | 241.28 | 8 | 8 | 0.034% | 94.62 | PDE |
| **Hyaluronidase: 0.02%** | | | | | | | | |
| 221 | XP_013928511.1 | *Thamnophis sirtalis* | 141.83 | 4 | 1 | 0.010% | 52.12 | Hyaluronidase |

| | | | | | | | | |
|---|---|---|---|---|---|---|---|---|
| 222 | XP_032077741.1 | *Thamnophis elegans* | 141.83 | 4 | 1 | 0.010% | 58.84 | Hyaluronidase |
| 223 | XP_026524834 | *Notechis scutatus* | 200.89 | 6 | 1 | 0.001% | 54.31 | Hyaluronidase |
| 224 | XP_026524834 | *Notechis scutatus* | 200.89 | 6 | 1 | 0.001% | 54.31 | Hyaluronidase |
| **Phospholipase B (PLB): 0.001%** | | | | | | | | |
| 225 | JAC94989.1 | *Opheodrys aestivus* | 86.74 | 2 | 1 | 0.001% | 63.91 | PLB |

**S4C Table.** The proteomic composition of *B. romulusi* venom.

| Sr. No. | Accession | Species | -10lgP | #Peptides | #Unique | Relative abundance of toxin hit (%) | Avg. Mass (KDa) | Family |
|---|---|---|---|---|---|---|---|---|
| **Phospholipase A2 ($PLA_2$): 42.34%** | | | | | | | | |
| 1 | 0702209A | *Bungarus multicinctus* | 331.88 | 41 | 15 | 9.055% | 11.39 | PLA2 |
| 2 | ADF50037 | *Bungarus flaviceps* | 322.68 | 30 | 4 | 5.453% | 19.18 | PLA2 |
| 3 | AAR19228 | *Bungarus caeruleus* | 312.93 | 27 | 19 | 3.512% | 13.52 | PLA2 |

| | | | | | | | | |
|---|---|---|---|---|---|---|---|---|
| 4 | ADF50037 | *Bungarus flaviceps* | 221.41 | 9 | 6 | 3.347% | 19.32 | PLA2 |
| 5 | ADF50037 | *Bungarus flaviceps* | 221.41 | 9 | 6 | 3.347% | 19.36 | PLA2 |
| 6 | 1PO8 | *Bungarus caeruleus* | 199.58 | 9 | 5 | 3.009% | 13.20 | PLA2 |
| 7 | 1TC8 | *Bungarus caeruleus* | 199.58 | 9 | 5 | 3.009% | 13.20 | PLA2 |
| 8 | AAS20530.1 | *Bungarus caeruleus* | 199.58 | 9 | 5 | 3.009% | 15.06 | PLA2 |
| 9 | Q6SLM2.1 | *Bungarus caeruleus* | 199.58 | 9 | 5 | 3.009% | 15.06 | PLA2 |
| 10 | CAA37482 | *Bungarus multicinctus* | 188.15 | 7 | 6 | 2.602% | 10.07 | PLA2 |
| 11 | 0702209A | *Bungarus multicinctus* | 328.36 | 37 | 15 | 2.341% | 11.32 | PLA2 |
| 12 | CAA37482 | *Bungarus multicinctus* | 337.39 | 36 | 9 | 0.240% | 11.18 | PLA2 |
| 13 | Q9DF52.1 | *Bungarus caeruleus* | 71.14 | 2 | 1 | 0.147% | 15.77 | PLA2 |
| 14 | AAG13412.1 | *Bungarus caeruleus* | 71.14 | 2 | 1 | 0.147% | 15.77 | PLA2 |
| 15 | ABU63164.1 | *Bungarus fasciatus* | 226.51 | 8 | 2 | 0.047% | 15.95 | PLA2 |
| 16 | ABU63166.1 | *Bungarus fasciatus* | 226.51 | 8 | 2 | 0.047% | 16.00 | PLA2 |
| 17 | P0C551 | *Bungarus candidus* | 186.46 | 8 | 4 | 0.020% | 20.10 | PLA2 |
| **Acetylcholinesterase (AChE): 36.58%** | | | | | | | | |
| 18 | AAC59905 | *Bungarus fasciatus* | 391.12 | 60 | 45 | 7.316% | 63.95 | AChE |
| 19 | AAC59905 | *Bungarus fasciatus* | 391.12 | 60 | 45 | 7.316% | 64.77 | AChE |
| 20 | AAC59905 | *Bungarus fasciatus* | 391.12 | 60 | 45 | 7.316% | 64.77 | AChE |
| 21 | AAC59905 | *Bungarus fasciatus* | 391.12 | 60 | 45 | 7.316% | 64.77 | AChE |
| 22 | AAC59905 | *Bungarus fasciatus* | 391.12 | 60 | 45 | 7.316% | 64.77 | AChE |
| 23 | XP_025032622.1 | *Python bivittatus* | 115.41 | 3 | 1 | 0.001% | 67.50 | AChE |
| 24 | XP_025032620.1 | *Python bivittatus* | 115.41 | 3 | 1 | 0.001% | 67.50 | AChE |
| 25 | XP_025032621.1 | *Python bivittatus* | 115.41 | 3 | 1 | 0.001% | 67.50 | AChE |
| 26 | JAC94932.1 | *Python regius* | 115.41 | 3 | 1 | 0.001% | 69.04 | AChE |
| 27 | XP_025032619.1 | *Python bivittatus* | 115.41 | 3 | 1 | 0.001% | 72.05 | AChE |

| | | | | | | | | |
|---|---|---|---|---|---|---|---|---|
| **Three-finger toxins (3FTx): 9.96%** | | | | | | | | |
| 28 | AAD41806 | *Bungarus multicinctus* | 274.4 | 16 | 16 | 8.604% | 9.90 | Type-X α-neurotoxin |
| 29 | ETE56933 | *Ophiophagus hannah* | 300.21 | 23 | 23 | 0.849% | 15.87 | Type-X α-neurotoxin |
| 30 | AAD41806 | *Bungarus multicinctus* | 211.53 | 9 | 9 | 0.345% | 9.83 | Type-X α-neurotoxin |
| 31 | AAL30059 | *Bungarus candidus* | 182.89 | 6 | 6 | 0.082% | 9.84 | Type II (long) α-neurotoxin |
| 32 | CAA06887 | *Bungarus multicinctus multicinctus* | 138.09 | 2 | 2 | 0.015% | 11.25 | Type I (short) α-neurotoxin |
| 33 | AAL30054.1 | *Bungarus candidus* | 67.87 | 1 | 1 | 0.011% | 9.58 | Type II (long) κ-neurotoxin |
| 34 | Q8AY56.1 | *Bungarus candidus* | 67.87 | 1 | 1 | 0.011% | 9.58 | Type II (long) κ-neurotoxin |
| 35 | AAC83997.1 | *Bungarus multicinctus* | 56.12 | 1 | 1 | 0.001% | 7.98 | Type II (long) α-neurotoxin |
| 36 | AAC83990.1 | *Bungarus multicinctus* | 56.12 | 1 | 1 | 0.001% | 8.02 | Type II (long) α-neurotoxin |
| 37 | P01385.1 | *Acanthophis antarcticus* | 56.12 | 1 | 1 | 0.001% | 8.14 | Type II (long) α-neurotoxin |
| 38 | 0512217A | *Notechis scutatus* | 56.12 | 1 | 1 | 0.001% | 8.06 | Type II (long) α-neurotoxin |
| 39 | AAC83992.1 | *Bungarus multicinctus* | 56.12 | 1 | 1 | 0.001% | 7.99 | Type II (long) α-neurotoxin |
| 40 | 1HAJ | *Bungarus multicinctus* | 56.12 | 1 | 1 | 0.001% | 7.99 | Type II (long) α-neurotoxin |
| 41 | 1IDH | *Bungarus multicinctus* | 56.12 | 1 | 1 | 0.001% | 7.99 | Type II (long) α-neurotoxin |

| | | | | | | | | |
|---|---|---|---|---|---|---|---|---|
| 42 | 2BTX | *Bungarus multicinctus* | 56.12 | 1 | 1 | 0.001% | 7.99 | Type II (long) $\alpha$-neurotoxin |
| 43 | 1IK8 | *Bungarus multicinctus* | 56.12 | 1 | 1 | 0.001% | 7.99 | Type II (long) $\alpha$-neurotoxin |
| 44 | 1BXP | *Bungarus multicinctus* | 56.12 | 1 | 1 | 0.001% | 7.99 | Type II (long) $\alpha$-neurotoxin |
| 45 | 1ABT | *Bungarus multicinctus* | 56.12 | 1 | 1 | 0.001% | 7.99 | Type II (long) $\alpha$-neurotoxin |
| 46 | 1IKC | *Bungarus multicinctus* | 56.12 | 1 | 1 | 0.001% | 7.99 | Type II (long) $\alpha$-neurotoxin |
| 47 | AAC83994.1 | *Bungarus multicinctus* | 56.12 | 1 | 1 | 0.001% | 7.99 | Type II (long) $\alpha$-neurotoxin |
| 48 | AAC83991.1 | *Bungarus multicinctus* | 56.12 | 1 | 1 | 0.001% | 7.99 | Type II (long) $\alpha$-neurotoxin |
| 49 | 1IDL | *Bungarus multicinctus* | 56.12 | 1 | 1 | 0.001% | 7.99 | Type II (long) $\alpha$-neurotoxin |
| 50 | 1HOY | *Bungarus multicinctus* | 56.12 | 1 | 1 | 0.001% | 8.00 | Type II (long) $\alpha$-neurotoxin |
| 51 | AAC83993.1 | *Bungarus multicinctus* | 56.12 | 1 | 1 | 0.001% | 7.99 | Type II (long) $\alpha$-neurotoxin |
| 52 | 720920A | *Bungarus multicinctus* | 56.12 | 1 | 1 | 0.001% | 7.99 | Type II (long) $\alpha$-neurotoxin |
| 53 | 2ABX | *Bungarus multicinctus* | 56.12 | 1 | 1 | 0.001% | 7.99 | Type II (long) $\alpha$-neurotoxin |
| 54 | CAM11311.1 | *Bungarus candidus* | 56.12 | 1 | 1 | 0.001% | 8.26 | Type II (long) $\alpha$-neurotoxin |
| 55 | ADN67593.1 | *Bungarus multicinctus* | 56.12 | 1 | 1 | 0.001% | 8.26 | Type II (long) $\alpha$-neurotoxin |
| 56 | CAM11305.1 | *Bungarus candidus* | 56.12 | 1 | 1 | 0.001% | 8.26 | Type II (long) $\alpha$-neurotoxin |
| 57 | CAM11309.1 | *Bungarus candidus* | 56.12 | 1 | 1 | 0.001% | 8.26 | Type II (long) $\alpha$-neurotoxin |

| | | | | | | | | |
|---|---|---|---|---|---|---|---|---|
| 58 | CAD92407.1 | *Bungarus candidus* | 56.12 | 1 | 1 | 0.001% | 8.26 | Type II (long) α-neurotoxin |
| 59 | CAM11312.1 | *Bungarus candidus* | 56.12 | 1 | 1 | 0.001% | 8.26 | Type II (long) α-neurotoxin |
| 60 | CAM11307.1 | *Bungarus multicinctus* | 56.12 | 1 | 1 | 0.001% | 8.26 | Type II (long) α-neurotoxin |
| 61 | CAM11310.1 | *Bungarus candidus* | 56.12 | 1 | 1 | 0.001% | 8.26 | Type II (long) α-neurotoxin |
| 62 | CAM11304.1 | *Bungarus candidus* | 56.12 | 1 | 1 | 0.001% | 8.26 | Type II (long) α-neurotoxin |
| 63 | CAM11302.1 | *Bungarus caeruleus* | 56.12 | 1 | 1 | 0.001% | 8.31 | Type II (long) α-neurotoxin |
| 64 | D2N116.1 | *Bungarus caeruleus* | 56.12 | 1 | 1 | 0.001% | 8.31 | Type II (long) α-neurotoxin |
| 65 | CAM11303.1 | *Bungarus candidus* | 56.12 | 1 | 1 | 0.001% | 8.23 | Type II (long) α-neurotoxin |
| 66 | CAM11306.1 | *Bungarus candidus* | 56.12 | 1 | 1 | 0.001% | 8.26 | Type II (long) α-neurotoxin |
| 67 | Q8UW29.1 | *Hydrophis hardwickii* | 56.12 | 1 | 1 | 0.001% | 10.22 | Type II (long) α-neurotoxin |
| 68 | AAL54892.1 | *Hydrophis hardwickii* | 56.12 | 1 | 1 | 0.001% | 10.22 | Type II (long) α-neurotoxin |
| 69 | A3FM53.1 | *Hydrophis hardwickii* | 56.12 | 1 | 1 | 0.001% | 10.41 | Type II (long) α-neurotoxin |
| 70 | ABN54806.1 | *Hydrophis hardwickii* | 56.12 | 1 | 1 | 0.001% | 10.41 | Type II (long) α-neurotoxin |
| 71 | JAA74770.1 | *Echiopsis curta* | 56.12 | 1 | 1 | 0.001% | 10.26 | Type II (long) α-neurotoxin |
| 72 | P01384.2 | *Notechis scutatus scutatus* | 56.12 | 1 | 1 | 0.001% | 10.29 | Type II (long) α-neurotoxin |
| 73 | ABK63537.1 | *Notechis scutatus* | 56.12 | 1 | 1 | 0.001% | 10.29 | Type II (long) α-neurotoxin |

| | | | | | | | | |
|---|---|---|---|---|---|---|---|---|
| 74 | JAA74654.1 | *Acanthophis wellsi* | 56.12 | 1 | 1 | 0.001% | 10.39 | Type II (long) α-neurotoxin |
| 75 | AAC83981.1 | *Bungarus multicinctus* | 56.12 | 1 | 1 | 0.001% | 10.29 | Type II (long) α-neurotoxin |
| 76 | AAL30056.1 | *Bungarus candidus* | 56.12 | 1 | 1 | 0.001% | 10.29 | Type II (long) α-neurotoxin |
| **Snake venom metalloproteinase (Disintegrin-like): 6.38%** | | | | | | | | |
| 77 | ABN72537 | *Bungarus multicinctus* | 201.08 | 11 | 8 | 2.626% | 14.13 | Disintegrin-like |
| 78 | ABN72537 | *Bungarus multicinctus* | 201.08 | 11 | 8 | 2.626% | 17.02 | Disintegrin-like |
| 79 | ABN72537 | *Bungarus multicinctus* | 364.17 | 48 | 8 | 0.773% | 66.30 | Disintegrin-like |
| 80 | ABQ01136 | *Oxyuranus scutellatus* | 180.6 | 7 | 5 | 0.239% | 37.26 | Disintegrin-like |
| 81 | ABN72536 | *Bungarus fasciatus* | 179.26 | 8 | 8 | 0.068% | 17.07 | Disintegrin-like |
| 82 | ABQ01138 | *Notechis scutatus* | 239.39 | 15 | 13 | 0.043% | 21.92 | Disintegrin-like |
| **β-bungarotoxin: 3.85%** | | | | | | | | |
| 83 | Q8QFW3.1 | *Bungarus caeruleus* | 269.83 | 18 | 7 | 1.130% | 16.12 | β-bungarotoxin |
| 84 | AAL87004.1 | *Bungarus caeruleus* | 269.83 | 18 | 7 | 1.130% | 16.12 | β-bungarotoxin |
| 85 | Q8QFW4.1 | *Bungarus caeruleus* | 306.09 | 24 | 17 | 0.724% | 16.36 | β-bungarotoxin |
| 86 | AAL87003.1 | *Bungarus caeruleus* | 306.09 | 24 | 17 | 0.724% | 16.36 | β-bungarotoxin |
| 87 | ABU63164 | *Bungarus fasciatus* | 278.54 | 17 | 2 | 0.122% | 14.06 | β-bungarotoxin |
| 88 | ABG90492.1 | *Bungarus multicinctus* | 289.92 | 21 | 1 | 0.020% | 13.16 | β-bungarotoxin |
| **L-amino-acid oxidase (LAAO): 0.38%** | | | | | | | | |
| 89 | ABN72539 | *Bungarus multicinctus* | 222.97 | 13 | 3 | 0.116% | 47.27 | LAAO |
| 90 | ABN72539 | *Bungarus multicinctus* | 222.97 | 13 | 3 | 0.116% | 47.27 | LAAO |
| 91 | ABN72539 | *Bungarus multicinctus* | 222.97 | 13 | 3 | 0.116% | 47.27 | LAAO |

| | | | | | | | | |
|---|---|---|---|---|---|---|---|---|
| 92 | ABN72539.1 | *Bungarus multicinctus* | 232.45 | 12 | 2 | 0.018% | 58.81 | LAAO |
| 93 | A8QL51.1 | *Bungarus multicinctus* | 232.45 | 12 | 2 | 0.018% | 58.81 | LAAO |
| **Vascular endothelial growth factor (VEGF): 0.26%** | | | | | | | | |
| 94 | XP_026551346 | *Pseudonaja textilis* | 178.85 | 9 | 9 | 0.041% | 16.47 | VEGF |
| 95 | XP_026551346 | *Pseudonaja textilis* | 178.85 | 9 | 9 | 0.041% | 16.47 | VEGF |
| 96 | XP_026551346 | *Pseudonaja textilis* | 178.85 | 9 | 9 | 0.041% | 16.47 | VEGF |
| 97 | XP_026551346 | *Pseudonaja textilis* | 178.85 | 9 | 9 | 0.041% | 17.24 | VEGF |
| 98 | JAA74898 | *Pseudonaja modesta* | 178.85 | 9 | 9 | 0.041% | 22.36 | VEGF |
| 99 | JAA74898 | *Pseudonaja modesta* | 178.85 | 9 | 9 | 0.041% | 22.36 | VEGF |
| 100 | XP_026555459 | *Pseudonaja textilis* | 137.59 | 5 | 5 | 0.004% | 32.63 | VEGF |
| 101 | LAB17921 | *Micrurus spixii* | 137.59 | 5 | 5 | 0.004% | 47.59 | VEGF |
| 102 | LAB17921 | *Micrurus spixii* | 137.59 | 5 | 5 | 0.004% | 47.59 | VEGF |
| **5'-nucleotidase: 0.11%** | | | | | | | | |
| 103 | JAB52815 | *Micrurus fulvius* | 343.67 | 35 | 28 | 0.105% | 64.64 | 5'-nucleotidase |
| **Peptidase S1: 0.07%** | | | | | | | | |
| 104 | XP_026555558.1 | *Pseudonaja textilis* | 189.93 | 9 | 9 | 0.063% | 61.35 | Peptidase S1 |
| 105 | ETE73401.1 | *Ophiophagus hannah* | 100.63 | 2 | 2 | 0.001% | 138.99 | Peptidase S1 |
| 106 | XP_026580326.1 | *Pseudonaja textilis* | 100.63 | 2 | 2 | 0.001% | 39.29 | Peptidase S1 |
| 107 | JAS05177.1 | *Micrurus tener* | 100.63 | 2 | 2 | 0.001% | 48.12 | Peptidase S1 |
| 108 | JAI09074.1 | *Micrurus fulvius* | 100.63 | 2 | 2 | 0.001% | 48.12 | Peptidase S1 |
| 109 | JAS05063.1 | *Micrurus fulvius* | 100.63 | 2 | 2 | 0.001% | 48.12 | Peptidase S1 |
| 110 | JAB54462.1 | *Micrurus fulvius* | 100.63 | 2 | 2 | 0.001% | 48.12 | Peptidase S1 |
| 111 | XP_026548111.1 | *Notechis scutatus* | 100.63 | 2 | 2 | 0.001% | 44.79 | Peptidase S1 |

| | | | | | | | | |
|---|---|---|---|---|---|---|---|---|
| **Hyaluronidase: 0.04%** | | | | | | | | |
| 112 | XP_026524834 | *Notechis scutatus* | 334.73 | 32 | 13 | 0.043% | 54.36 | Hyaluronidase |
| **Cysteine-rich secretory proteins (CRISP): 0.02%** | | | | | | | | |
| 113 | ACE73577.1 | *Bungarus candidus* | 69.57 | 1 | 1 | 0.001% | 26.46 | CRISP |
| 114 | ACE73578.1 | *Bungarus candidus* | 69.57 | 1 | 1 | 0.001% | 26.44 | CRISP |
| 115 | ACE73577 | *Bungarus candidus* | 69.57 | 1 | 1 | 0.001% | 26.38 | CRISP |
| 116 | ACE73577 | *Bungarus candidus* | 69.57 | 1 | 1 | 0.001% | 26.38 | CRISP |
| 117 | ACE73577 | *Bungarus candidus* | 69.57 | 1 | 1 | 0.001% | 26.38 | CRISP |
| 118 | ACE73577 | *Bungarus candidus* | 69.57 | 1 | 1 | 0.001% | 26.38 | CRISP |
| 119 | ACE73577 | *Bungarus candidus* | 69.57 | 1 | 1 | 0.001% | 27.81 | CRISP |
| 120 | ACE73577 | *Bungarus candidus* | 69.57 | 1 | 1 | 0.001% | 31.26 | CRISP |
| 121 | P81993.1 | *Bungarus candidus* | 69.57 | 1 | 1 | 0.001% | 7.75 | CRISP |
| 122 | ACE73577 | *Bungarus candidus* | 69.57 | 1 | 1 | 0.001% | 19.40 | CRISP |
| 123 | ACN93671.1 | *Ophiophagus hannah* | 69.57 | 1 | 1 | 0.001% | 26.31 | CRISP |
| 124 | ACH73168.1 | *Naja kaouthia* | 69.57 | 1 | 1 | 0.001% | 26.22 | CRISP |
| 125 | AAP20603.1 | *Naja atra* | 69.57 | 1 | 1 | 0.001% | 26.25 | CRISP |
| 126 | Q7ZZN8.1 | *Naja atra* | 69.57 | 1 | 1 | 0.001% | 26.25 | CRISP |
| 127 | P84808.2 | *Naja kaouthia* | 69.57 | 1 | 1 | 0.001% | 26.22 | CRISP |
| 128 | AHZ08822.1 | *Micropechis ikaheca* | 69.57 | 1 | 1 | 0.001% | 26.22 | CRISP |
| 129 | XP_032069593.1 | *Thamnophis elegans* | 69.57 | 1 | 1 | 0.001% | 26.60 | CRISP |
| 130 | JAC94992.1 | *Opheodrys aestivus* | 69.57 | 1 | 1 | 0.001% | 26.75 | CRISP |
| **Phospholipase B (PLB): 0.01%** | | | | | | | | |
| 131 | ACR78473 | *Drysdalia coronoides* | 104.9 | 2 | 2 | 0.004% | 64.04 | PLB |

| | | | | | | | | |
|---|---|---|---|---|---|---|---|---|
| 132 | ACR78473 | *Drysdalia coronoides* | 104.9 | 2 | 2 | 0.004% | 64.04 | PLB |

**Table S5A. Details of RNA isolation**

| Sample | Tissue type | Concentration ng/µL | RIN |
|---|---|---|---|
| ***B. sindanus*** | **Venom gland** | 188 | 8.0 |
| | **Intestine** | 7420 | 9.0 |
| ***B. caeruleus*** | **Venom gland** | 1260 | 9.6 |
| | **Intestine** | 230 | 7.0 |
| ***B. romulusi*** | **Venom gland** | 3500 | 7.7 |
| | **Intestine** | 1040 | 8.3 |

**Table S5B. Tissue transcriptome sequencing and assembly**

| | **Reads** | | | | **Assembly** | |
|---|---|---|---|---|---|---|
| **Tissue** | **Raw** | **Filtered** | **Expressed** | **Annotated** | **Transcripts** | **N50** |
| ***B. sindanus*** | | | | | | |
| Venom gland | 34,952,562 | 33,332,796 | 27,599,732 | 54,793 | 229,763 | 1,436 |
| Intestine | 12,058,000 | 10,740,803 | 9,153,810 | | | |
| ***B. caeruleus*** | | | | | | |
| Venom gland | 51,842,239 | 46,397,238 | 39,446,271 | 98,441 | 308,570 | 1,701 |
| Intestine | 55,336,796 | 51,104,996 | 43,471,869 | | | |
| ***B. romulusi*** | | | | | | |
| Venom gland | 24,343,212 | 21,256,495 | 16,583,652 | 58,007 | 225,522 | 1,863 |
| Intestine | 28,413,016 | 24,882,567 | 21,853,659 | | | |

### Figure S6A. *Bungarus sindanus*

| S. No. | Isoform | VG | Int | M | Family | Subtype | Abundance | Accession | Scientific name |
|---|---|---|---|---|---|---|---|---|---|
| 1 | TRINITY_DN526_c0_g1_i10 | 5667 | 0 | 5.2821 | 5'-nucleotidase | | 0.23% | JAB52815 | Micrurus fulvius |
| 2 | TRINITY_DN526_c0_g1_i36 | 4729 | 0 | 5.2821 | 5'-nucleotidase | | | JAB52815 | Micrurus fulvius |
| 3 | TRINITY_DN526_c0_g1_i28 | 2599 | 6 | 5.2821 | 5'-nucleotidase | | | XP_013918203 | Thamnophis sirtalis |
| 4 | TRINITY_DN526_c0_g1_i8 | 1990 | 0 | 5.2821 | 5'-nucleotidase | | | JAB52815 | Micrurus fulvius |
| 5 | TRINITY_DN526_c0_g1_i22 | 690 | 62 | 5.2821 | 5'-nucleotidase | | | JAB52815 | Micrurus fulvius |
| 6 | TRINITY_DN526_c0_g1_i9 | 356 | 11 | 5.2821 | 5'-nucleotidase | | | XP_026582322 | Pseudonaja textilis |
| 7 | TRINITY_DN1357_c0_g1_i9 | 575 | 22 | 2.2921 | 5'-nucleotidase | | | XP_026557747 | Pseudonaja textilis |
| 8 | TRINITY_DN1357_c0_g1_i15 | 194 | 0 | 2.2921 | 5'-nucleotidase | | | XP_026557747 | Pseudonaja textilis |
| 9 | TRINITY_DN3400_c0_g1_i4 | 614 | 3 | 3.9753 | 5'-nucleotidase | | | LAB37863 | Micrurus spixii |
| 10 | TRINITY_DN14672_c1_g2_i1 | 144 | 17 | 2.2259 | 5'-nucleotidase | | | XP_026539047 | Notechis scutatus |
| 11 | TRINITY_DN658_c0_g1_i17 | 25535 | 0 | 10.523 | Acetylcholinesterase | | 0.95% | AAC59905 | Bungarus fasciatus |
| 12 | TRINITY_DN658_c0_g1_i23 | 16198 | 0 | 10.523 | Acetylcholinesterase | | | AAC59905 | Bungarus fasciatus |
| 13 | TRINITY_DN658_c0_g1_i25 | 12626 | 0 | 10.523 | Acetylcholinesterase | | | Q92035 | Bungarus fasciatus |
| 14 | TRINITY_DN658_c0_g1_i26 | 10794 | 10 | 10.523 | Acetylcholinesterase | | | AAC59905 | Bungarus fasciatus |
| 15 | TRINITY_DN658_c0_g1_i11 | 3303 | 0 | 10.523 | Acetylcholinesterase | | | AAC59905 | Bungarus fasciatus |
| 16 | TRINITY_DN658_c0_g1_i20 | 3090 | 0 | 10.523 | Acetylcholinesterase | | | Q92035 | Bungarus fasciatus |
| 17 | TRINITY_DN658_c0_g1_i3 | 620 | 0 | 10.523 | Acetylcholinesterase | | | Q92035 | Bungarus fasciatus |
| 18 | TRINITY_DN658_c0_g1_i7 | 382 | 0 | 10.523 | Acetylcholinesterase | | | Q92035 | Bungarus fasciatus |
| 19 | TRINITY_DN658_c0_g1_i15 | 267 | 0 | 10.523 | Acetylcholinesterase | | | Q92035 | Bungarus fasciatus |
| 20 | TRINITY_DN658_c0_g1_i1 | 233 | 0 | 10.523 | Acetylcholinesterase | | | Q92035 | Bungarus fasciatus |
| 21 | TRINITY_DN95_c7_g3_i2 | 240 | 0 | 9.3464 | Acetylcholinesterase | | | XP_026576484 | Pseudonaja textilis |
| 22 | TRINITY_DN16_c0_g2_i2 | 188135 | 7 | 12.659 | CRISP | | 6.30% | ACE73577 | Bungarus candidus |
| 23 | TRINITY_DN16_c0_g2_i6 | 175526 | 7 | 12.659 | CRISP | | | ACE73577 | Bungarus candidus |
| 24 | TRINITY_DN16_c0_g2_i11 | 118278 | 22 | 12.659 | CRISP | | | ACE73577 | Bungarus candidus |
| 25 | TRINITY_DN16_c0_g2_i14 | 4528 | 0 | 12.659 | CRISP | | | ACE73577 | Bungarus candidus |
| 26 | TRINITY_DN16_c0_g2_i13 | 613 | 0 | 12.659 | CRISP | | | XP_026546646 | Notechis scutatus |
| 27 | TRINITY_DN16_c0_g2_i8 | 284 | 0 | 12.659 | CRISP | | | ETE62137 | Ophiophagus hannah |
| 28 | TRINITY_DN1102_c0_g1_i1 | 1485 | 72 | 3.5098 | CTL | | 0.33% | XP_026577709 | Pseudonaja textilis |
| 29 | TRINITY_DN2_c1_g2_i1 | 2138 | 1 | 7.1521 | CTL | | | ADF50042 | Bungarus flaviceps |
| 30 | TRINITY_DN2_c1_g2_i2 | 1198 | 12 | 7.1521 | CTL | | | ABP94128 | Pseudechis porphyriacus |
| 31 | TRINITY_DN111_c1_g2_i10 | 945 | 8 | 5.9946 | CTL | | | XP_026575421 | Pseudonaja textilis |
| 32 | TRINITY_DN111_c1_g2_i1 | 107 | 0 | 5.9946 | CTL | | | LAA90676 | Micrurus lemniscatus lemniscatus |
| 33 | TRINITY_DN286_c0_g1_i28 | 16976 | 0 | 11.125 | CTL | | | ACC67942 | Oxyuranus microlepidotus |
| 34 | TRINITY_DN286_c0_g1_i1 | 1735 | 0 | 11.125 | CTL | | | ABP94094 | Oxyuranus scutellatus |
| 35 | TRINITY_DN286_c0_g1_i9 | 866 | 0 | 11.125 | CTL | | | ABP94094 | Oxyuranus scutellatus |
| 36 | TRINITY_DN286_c0_g1_i7 | 135 | 0 | 11.125 | CTL | | | ABP94095 | Oxyuranus scutellatus |
| 37 | TRINITY_DN286_c0_g1_i11 | 113 | 4 | 11.125 | CTL | | | JAB52841 | Micrurus fulvius |
| 38 | TRINITY_DN12_c1_g1_i10 | 1326 | 19 | 3.8658 | Disintegrin-like | | | XP_026527224 | Notechis scutatus |
| 39 | TRINITY_DN87082_c0_g1_i1 | 2594 | 88 | 4.025 | Disintegrin-like | | | XP_034289867 | Pantherophis guttatus |
| 40 | TRINITY_DN2791_c0_g1_i2 | 247 | 23 | 2.0976 | Disintegrin-like | | | ETE74105 | Ophiophagus hannah |
| 41 | TRINITY_DN2791_c0_g1_i1 | 187 | 33 | 2.0976 | Disintegrin-like | | | LAA74418 | Micrurus lemniscatus lemniscatus |
| 42 | TRINITY_DN10_c0_g1_i3 | 16256 | 0 | 10.764 | Disintegrin-like | | | LAA20517 | Micrurus lemniscatus carvalhoi |
| 43 | TRINITY_DN10_c0_g1_i16 | 16150 | 12 | 10.764 | Disintegrin-like | | | ABN72537 | Bungarus multicinctus |
| 44 | TRINITY_DN10_c0_g1_i10 | 12234 | 2 | 10.764 | Disintegrin-like | | | ABN72537 | Bungarus multicinctus |
| 45 | TRINITY_DN10_c0_g1_i11 | 8426 | 0 | 10.764 | Disintegrin-like | | | LAA20517 | Micrurus lemniscatus carvalhoi |
| 46 | TRINITY_DN10_c0_g1_i4 | 8296 | 0 | 10.764 | Disintegrin-like | | | LAA20517 | Micrurus lemniscatus carvalhoi |
| 47 | TRINITY_DN10_c0_g1_i7 | 6015 | 0 | 10.764 | Disintegrin-like | | | AEH95531 | Drysdalia coronoides |
| 48 | TRINITY_DN10_c0_g1_i5 | 1540 | 6 | 10.764 | Disintegrin-like | | | ABN72537 | Bungarus multicinctus |
| 49 | TRINITY_DN10_c0_g1_i6 | 1298 | 0 | 10.764 | Disintegrin-like | | | ABN72537 | Bungarus multicinctus |

| | | | | | | | | | |
|---|---|---|---|---|---|---|---|---|---|
| 50 | TRINITY_DN10_c0_g1_i8 | 1245 | 0 | 10.764 | Disintegrin-like | | | ABN72537 | Bungarus multicinctus |
| 51 | TRINITY_DN10_c0_g1_i18 | 803 | 0 | 10.764 | Disintegrin-like | | | ABN72536 | Bungarus fasciatus |
| 52 | TRINITY_DN10_c0_g1_i1 | 132 | 0 | 10.764 | Disintegrin-like | | **1.13%** | ABN72537 | Bungarus multicinctus |
| 53 | TRINITY_DN8820_c0_g1_i2 | 860 | 19 | 4.6437 | Disintegrin-like | | | ETE68273 | Ophiophagus hannah |
| 54 | TRINITY_DN8043_c0_g1_i1 | 542 | 17 | 4.1723 | Disintegrin-like | | | XP_034279507 | Pantherophis guttatus |
| 55 | TRINITY_DN9584_c0_g1_i2 | 328 | 0 | 6.0593 | Disintegrin-like | | | XP_034280049 | Pantherophis guttatus |
| 56 | TRINITY_DN9584_c0_g1_i6 | 237 | 0 | 6.0593 | Disintegrin-like | | | XP_026577550 | Pseudonaja textilis |
| 57 | TRINITY_DN9584_c0_g1_i7 | 147 | 1 | 6.0593 | Disintegrin-like | | | XP_034280049 | Pantherophis guttatus |
| 58 | TRINITY_DN1565_c0_g1_i2 | 810 | 5 | 6.4951 | Disintegrin-like | | | ABN72537 | Bungarus multicinctus |
| 59 | TRINITY_DN1565_c0_g1_i1 | 170 | 1 | 6.4951 | Disintegrin-like | | | ABN72537 | Bungarus multicinctus |
| 60 | TRINITY_DN389_c1_g1_i1 | 395 | 2 | 6.868 | Disintegrin-like | | | ABQ01138 | Notechis scutatus |
| 61 | TRINITY_DN2189_c0_g2_i3 | 2691 | 0 | 12.296 | Disintegrin-like | | | ABK63559 | Demansia vestigiata |
| 62 | TRINITY_DN10_c4_g1_i1 | 2526 | 0 | 12.087 | Disintegrin-like | | | JAA74899 | Pseudonaja modesta |
| 63 | TRINITY_DN2873_c0_g1_i1 | 792 | 0 | 10.414 | Disintegrin-like | | | XP_026536946 | Notechis scutatus |
| 64 | TRINITY_DN5452_c0_g1_i1 | 543 | 0 | 9.869 | Disintegrin-like | | | ABN72537 | Bungarus multicinctus |
| 65 | TRINITY_DN5452_c0_g3_i1 | 399 | 0 | 9.4244 | Disintegrin-like | | | ABN72537 | Bungarus multicinctus |
| 66 | TRINITY_DN389_c0_g1_i2 | 248 | 0 | 8.8231 | Disintegrin-like | | | XP_013922279 | Thamnophis sirtalis |
| 67 | TRINITY_DN1030_c0_g1_i5 | 334 | 11 | 2.8671 | Hyaluronidase | | | ETE63480 | Ophiophagus hannah |
| 68 | TRINITY_DN3688_c2_g1_i4 | 9416 | 0 | 10.503 | Hyaluronidase | | | XP_026524834 | Notechis scutatus |
| 69 | TRINITY_DN3688_c2_g1_i7 | 473 | 0 | 10.503 | Hyaluronidase | | **0.14%** | XP_026524834 | Notechis scutatus |
| 70 | TRINITY_DN3688_c2_g1_i3 | 428 | 2 | 10.503 | Hyaluronidase | | | LAA86771 | Micrurus lemniscatus lemniscatus |
| 71 | TRINITY_DN3688_c2_g1_i6 | 182 | 2 | 10.503 | Hyaluronidase | | | XP_026524834 | Notechis scutatus |
| 72 | TRINITY_DN3_c0_g1_i38 | 819143 | 2 | 8.2948 | Kunitz | | | LAA53789 | Micrurus corallinus |
| 73 | TRINITY_DN3_c0_g1_i35 | 429 | 0 | 8.2948 | Kunitz | | | LAA53789 | Micrurus corallinus |
| 74 | TRINITY_DN3_c0_g1_i37 | 192 | 0 | 8.2948 | Kunitz | | | LAA53789 | Micrurus corallinus |
| 75 | TRINITY_DN855_c0_g1_i43 | 1239 | 323 | 3.3088 | Kunitz | | | XP_026543549 | Notechis scutatus |
| 76 | TRINITY_DN855_c0_g1_i76 | 1128 | 0 | 3.3088 | Kunitz | | | XP_026543549 | Notechis scutatus |
| 77 | TRINITY_DN855_c0_g1_i62 | 855 | 260 | 3.3088 | Kunitz | | | XP_026543549 | Notechis scutatus |
| 78 | TRINITY_DN855_c0_g1_i89 | 629 | 334 | 3.3088 | Kunitz | | **11.89%** | XP_026543549 | Notechis scutatus |
| 79 | TRINITY_DN8916_c1_g1_i4 | 92109 | 1 | 10.885 | Kunitz | | | ACR78501 | Drysdalia coronoides |
| 80 | TRINITY_DN2341_c0_g1_i1 | 2869 | 13 | 6.9313 | Kunitz | | | XP_026579404 | Pseudonaja textilis |
| 81 | TRINITY_DN2341_c1_g1_i1 | 170 | 1 | 6.5528 | Kunitz | | | XP_026581272 | Pseudonaja textilis |
| 82 | TRINITY_DN2565_c0_g1_i1 | 398 | 0 | 9.4208 | Kunitz | | | XP_026545967 | Notechis scutatus |
| 83 | TRINITY_DN1055_c0_g1_i1 | 132 | 0 | 8.5786 | Kunitz | | | XP_026579465 | Pseudonaja textilis |
| 84 | TRINITY_DN8381_c0_g1_i5 | 104 | 0 | 8.6911 | Kunitz | | | XP_032074676 | Thamnophis elegans |
| 85 | TRINITY_DN420_c0_g1_i9 | 18068 | 0 | 7.7758 | **LAAO** | | | ABN72539 | Bungarus multicinctus |
| 86 | TRINITY_DN420_c0_g1_i15 | 11506 | 0 | 7.7758 | **LAAO** | | **0.40%** | ABN72540 | Bungarus fasciatus |
| 87 | TRINITY_DN420_c0_g1_i14 | 688 | 5 | 7.7758 | **LAAO** | | | ABN72540 | Bungarus fasciatus |
| 88 | TRINITY_DN420_c0_g1_i8 | 605 | 0 | 7.7758 | **LAAO** | | | ABN72539 | Bungarus multicinctus |
| 89 | TRINITY_DN45_c1_g1_i2 | 120 | 0 | 9.9997 | Natriuretic peptide | | | ADK12002 | Ophiophagus hannah |
| 90 | TRINITY_DN45_c1_g1_i5 | 195105 | 0 | 9.9997 | Natriuretic peptide | | | ADK12002 | Ophiophagus hannah |
| 91 | TRINITY_DN45_c1_g1_i4 | 171746 | 10 | 9.9997 | Natriuretic peptide | | **4.85%** | ADK12002 | Ophiophagus hannah |
| 92 | TRINITY_DN45_c1_g1_i6 | 7494 | 192 | 9.9997 | Natriuretic peptide | | | JAS05144 | Micrurus tener |
| 93 | TRINITY_DN295_c0_g1_i2 | 614 | 1 | 11.107 | Natriuretic peptide | | | ADK12002 | Ophiophagus hannah |
| 94 | TRINITY_DN205_c0_g1_i8 | 8309 | 2 | 9.6574 | **NGF** | | | AAB25729 | Bungarus multicinctus |
| 95 | TRINITY_DN205_c0_g1_i15 | 1566 | 0 | 9.6574 | **NGF** | | | AAB25729 | Bungarus multicinctus |
| 96 | TRINITY_DN205_c0_g1_i11 | 1415 | 0 | 9.6574 | **NGF** | | | AAB25729 | Bungarus multicinctus |
| 97 | TRINITY_DN205_c0_g1_i3 | 1065 | 8 | 9.6574 | **NGF** | | **0.18%** | AAB25729 | Bungarus multicinctus |
| 98 | TRINITY_DN205_c0_g1_i7 | 627 | 0 | 9.6574 | **NGF** | | | AAB25729 | Bungarus multicinctus |
| 99 | TRINITY_DN205_c0_g1_i5 | 393 | 0 | 9.6574 | **NGF** | | | AAB25729 | Bungarus multicinctus |

| | | | | | | | | | |
|---|---|---|---|---|---|---|---|---|---|
| 100 | TRINITY_DN205_c0_g1_i13 | 388 | 0 | 9.6574 | NGF | | | AAB25729 | Bungarus multicinctus |
| 101 | TRINITY_DN205_c0_g1_i4 | 252 | 0 | 9.6574 | NGF | | | AAB25729 | Bungarus multicinctus |
| 102 | TRINITY_DN158_c0_g2_i21 | 416242 | 0 | 6.7164 | PLA2 | | | ACY68711 | Parasuta nigriceps |
| 103 | TRINITY_DN158_c0_g2_i5 | 19472 | 0 | 6.7164 | PLA2 | | | LAA18991 | Micrurus lemniscatus carvalhoi |
| 104 | TRINITY_DN158_c0_g1_i7 | 523798 | 0 | 8.1888 | PLA2 | | | AAB24834 | Bungarus fasciatus |
| 105 | TRINITY_DN158_c0_g1_i18 | 337319 | 0 | 8.1888 | PLA2 | | | ABG75909 | Bungarus fasciatus |
| 106 | TRINITY_DN158_c0_g1_i27 | 11669 | 0 | 8.1888 | PLA2 | | | LAA39992 | Micrurus corallinus |
| 107 | TRINITY_DN158_c0_g1_i36 | 3947 | 0 | 8.1888 | PLA2 | | | AAB24834 | Bungarus fasciatus |
| 108 | TRINITY_DN66_c3_g1_i2 | 3595 | 0 | 2.7671 | PLA2 | | | XP_034286728 | Pantherophis guttatus |
| 109 | TRINITY_DN66_c3_g1_i1 | 337 | 319 | 2.7671 | PLA2 | | | XP_034286728 | Pantherophis guttatus |
| 110 | TRINITY_DN4685_c0_g1_i9 | 340 | 0 | 2.6072 | PLA2 | | 23.33% | LAA68819 | Micrurus lemniscatus lemniscatus |
| 111 | TRINITY_DN4685_c0_g1_i20 | 208 | 0 | 2.6072 | PLA2 | | | LAA68819 | Micrurus lemniscatus lemniscatus |
| 112 | TRINITY_DN3_c4_g1_i16 | 1713 | 0 | 5.1741 | PLA2 | | | LAA68819 | Micrurus lemniscatus lemniscatus |
| 113 | TRINITY_DN3_c4_g1_i22 | 296 | 37 | 5.1741 | PLA2 | | | LAA68819 | Micrurus lemniscatus lemniscatus |
| 114 | TRINITY_DN241_c0_g1_i1 | 483423 | 4 | 16.026 | PLA2 | | | ABM88801 | Hoplocephalus stephensii |
| 115 | TRINITY_DN3025_c1_g2_i1 | 124 | 3 | 4.5127 | PLA2 | | | CAD24462 | Bungarus multicinctus |
| 116 | TRINITY_DN23151_c0_g1_i2 | 122 | 0 | 4.5243 | PLA2 | | | XP_026565791 | Pseudonaja textilis |
| 117 | TRINITY_DN5043_c1_g1_i8 | 1042 | 0 | 9.7582 | PLA2 | | | ADF50037 | Bungarus flaviceps |
| 118 | TRINITY_DN5043_c1_g1_i9 | 346 | 0 | 9.7582 | PLA2 | | | ADF50037 | Bungarus flaviceps |
| 119 | TRINITY_DN5043_c1_g1_i1 | 162 | 0 | 9.7582 | PLA2 | | | ADF50037 | Bungarus flaviceps |
| 120 | TRINITY_DN2464_c0_g1_i2 | 15349 | 74 | 5.1041 | PLB | | 0.20% | ACR78473 | Drysdalia coronoides |
| 121 | TRINITY_DN2464_c0_g1_i1 | 283 | 177 | 5.1041 | PLB | | | ACR78473 | Drysdalia coronoides |
| 122 | TRINITY_DN3093_c0_g1_i8 | 271 | 2 | 2.0355 | Serine Protease | | 0.02% | ABN72545 | Bungarus multicinctus |
| 123 | TRINITY_DN1_c16_g2_i4 | 1078 | 34 | 4.7938 | Serine Protease | | | XP_026562705 | Pseudonaja textilis |
| 124 | TRINITY_DN821_c0_g1_i7 | 57470 | 0 | 8.1587 | Type-I α-neurotoxin | | | CAA45882 | Bungarus multicinctus |
| 125 | TRINITY_DN821_c0_g1_i3 | 25406 | 0 | 8.1587 | Type-I α-neurotoxin | | | CAA45882 | Bungarus multicinctus |
| 126 | TRINITY_DN179_c0_g1_i11 | 85078 | 1 | 13.033 | Type-I α-neurotoxin | | | CAA06887 | Bungarus multicinctus multicinctus |
| 127 | TRINITY_DN179_c0_g1_i10 | 20468 | 0 | 13.033 | Type-I α-neurotoxin | | | CAA06887 | Bungarus multicinctus multicinctus |
| 128 | TRINITY_DN179_c0_g1_i3 | 19758 | 0 | 13.033 | Type-I α-neurotoxin | | | CAA06887 | Bungarus multicinctus multicinctus |
| 129 | TRINITY_DN4374_c0_g1_i2 | 165563 | 3 | 12.177 | Type-I α-neurotoxin | | | ETE58964 | Ophiophagus hannah |
| 130 | TRINITY_DN31129_c0_g1_i1 | 160874 | 29 | 11.095 | Type-II α-neurotoxin | | | AHZ08825 | Micropechis ikaheca |
| 131 | TRINITY_DN31129_c0_g1_i2 | 1544 | 7 | 11.095 | Type-II α-neurotoxin | | | CAJ77820 | Bungarus candidus |
| 132 | TRINITY_DN133_c4_g1_i5 | 896317 | 29 | 13.781 | Type-II α-neurotoxin | | | ETE56306 | Ophiophagus hannah |
| 133 | TRINITY_DN133_c4_g1_i1 | 3878 | 6 | 13.781 | Type-II α-neurotoxin | | | ETE56306 | Ophiophagus hannah |
| 134 | TRINITY_DN909_c0_g1_i13 | 3510 | 1 | 12.807 | Type-II α-neurotoxin | 64.44% | | CAA10359 | Bungarus multicinctus |
| 135 | TRINITY_DN909_c0_g1_i38 | 2921 | 0 | 12.807 | Type-II α-neurotoxin | | | CAA10359 | Bungarus multicinctus |
| 136 | TRINITY_DN909_c0_g1_i40 | 1653 | 0 | 12.807 | Type-II α-neurotoxin | | | LAA40709 | Micrurus corallinus |
| 137 | TRINITY_DN10768_c0_g1_i3 | 824 | 0 | 17.948 | Type-II α-neurotoxin | | | ETE56205 | Ophiophagus hannah |
| 138 | TRINITY_DN909_c0_g1_i11 | 564 | 0 | 12.807 | Type-II α-neurotoxin | | | LAA40709 | Micrurus corallinus |
| 139 | TRINITY_DN909_c0_g1_i17 | 548 | 0 | 12.807 | Type-II α-neurotoxin | | | ABK63538 | Tropidechis carinatus |
| 140 | TRINITY_DN909_c0_g1_i1 | 199 | 0 | 12.807 | Type-II α-neurotoxin | | 29.06% | LAA40709 | Micrurus corallinus |
| 141 | TRINITY_DN909_c0_g1_i37 | 177 | 0 | 12.807 | Type-II α-neurotoxin | | | ETE56521 | Ophiophagus hannah |
| 142 | TRINITY_DN2350_c0_g3_i1 | 854 | 0 | 10.522 | Type-II α-neurotoxin | | | LAA40709 | Micrurus corallinus |
| 143 | TRINITY_DN909_c1_g1_i2 | 389 | 0 | 9.9083 | Type-II α-neurotoxin | | | LAA40709 | Micrurus corallinus |
| 144 | TRINITY_DN3640_c0_g1_i5 | 147 | 0 | 11.252 | Type-II α-neurotoxin | | | ETE56205 | Ophiophagus hannah |
| 145 | TRINITY_DN3640_c0_g1_i10 | 120 | 0 | 11.252 | Type-II α-neurotoxin | | | LAA40709 | Micrurus corallinus |
| 146 | TRINITY_DN821_c0_g2_i2 | 187 | 0 | 12.673 | Unconventional 3FTx | | | CAD56381 | Bungarus multicinctus |
| 147 | TRINITY_DN179_c0_g1_i4 | 173811 | 3 | 13.033 | Unconventional 3FTx | | | AAL30060 | Bungarus candidus |
| 148 | TRINITY_DN179_c0_g1_i37 | 163976 | 4 | 13.033 | Unconventional 3FTx | | | CAB53358 | Bungarus multicinctus multicinctus |
| 149 | TRINITY_DN179_c0_g1_i15 | 131062 | 0 | 13.033 | Unconventional 3FTx | | | LAA65595 | Micrurus corallinus |

| | | | | | | | | | |
|---|---|---|---|---|---|---|---|---|---|
| 150 | TRINITY_DN179_c0_g1_i31 | 95299 | 2 | 13.033 | Unconventional 3FTx | | | AAL30059 | Bungarus candidus |
| 151 | TRINITY_DN1177_c0_g1_i30 | 126997 | 5 | 15.099 | Unconventional 3FTx | 35.56% | | CAA06885 | Bungarus multicinctus multicinctus |
| 152 | TRINITY_DN3640_c0_g2_i3 | 37781 | 1 | 11.648 | Unconventional 3FTx | | | CAB53358 | Bungarus multicinctus multicinctus |
| 153 | TRINITY_DN3640_c0_g2_i2 | 19948 | 7 | 11.648 | Unconventional 3FTx | | | CAB53358 | Bungarus multicinctus multicinctus |
| 154 | TRINITY_DN821_c0_g2_i1 | 47123 | 4 | 12.673 | Unconventional 3FTx | | | CAD56381 | Bungarus multicinctus |
| 155 | TRINITY_DN909_c0_g1_i25 | 2051 | 0 | 12.807 | Unconventional 3FTx | | | ETE56205 | Ophiophagus hannah |
| 156 | TRINITY_DN10768_c3_g1_i1 | 930 | 0 | 10.645 | Unconventional 3FTx | | | CAB53358 | Bungarus multicinctus multicinctus |
| 157 | TRINITY_DN21645_c1_g2_i1 | 104 | 0 | 7.4846 | Unconventional 3FTx | | | XP_034276006 | Pantherophis guttatus |
| 158 | TRINITY_DN596_c0_g1_i6 | 18639 | 0 | 7.6955 | VEGF | | | JAA74898 | Pseudonaja modesta |
| 159 | TRINITY_DN596_c0_g1_i4 | 6532 | 16 | 7.6955 | VEGF | | | XP_026551346 | Pseudonaja textilis |
| 160 | TRINITY_DN596_c0_g1_i2 | 5719 | 30 | 7.6955 | VEGF | | | XP_026551346 | Pseudonaja textilis |
| 161 | TRINITY_DN596_c0_g1_i1 | 3064 | 0 | 7.6955 | VEGF | | | XP_026551344 | Pseudonaja textilis |
| 162 | TRINITY_DN596_c0_g1_i10 | 744 | 0 | 7.6955 | VEGF | | 0.51% | XP_026551344 | Pseudonaja textilis |
| 163 | TRINITY_DN596_c0_g1_i5 | 667 | 0 | 7.6955 | VEGF | | | JAA74688 | Cacophis squamulosus |
| 164 | TRINITY_DN1555_c0_g1_i2 | 3490 | 40 | 5.0585 | VEGF | | | LAB17921 | Micrurus spixii |
| 165 | TRINITY_DN4014_c2_g2_i2 | 181 | 9 | 3.3181 | VEGF | | | XP_026542052 | Notechis scutatus |
| 166 | TRINITY_DN205_c0_g1_i9 | 607 | 0 | 9.6574 | VEGF | | | AAB25729 | Bungarus multicinctus |
| 167 | TRINITY_DN2536_c0_g1_i1 | 29714 | 4 | 12.002 | Vespryn | | **0.38%** | XP_026576508 | Pseudonaja textilis |
| 168 | TRINITY_DN158_c0_g2_i12 | 513486 | 17 | 6.7164 | β-bungarotoxin | | | BAD06267 | Bungarus candidus |
| 169 | TRINITY_DN158_c0_g1_i21 | 428465 | 178 | 8.1888 | β-bungarotoxin | | | CAD24462.1 | Micrurus corallinus |
| 170 | TRINITY_DN158_c0_g1_i11 | 128743 | 5 | 8.1888 | β-bungarotoxin | | | CAD24466 | Bungarus multicinctus |
| 171 | TRINITY_DN158_c0_g1_i31 | 31094 | 0 | 8.1888 | β-bungarotoxin | | | LAA39992 | Micrurus corallinus |
| 172 | TRINITY_DN158_c0_g1_i28 | 25784 | 0 | 8.1888 | β-bungarotoxin | | | LAA39992 | Micrurus corallinus |
| 173 | TRINITY_DN158_c0_g1_i9 | 4339 | 8 | 8.1888 | β-bungarotoxin | | | LAA39992 | Micrurus corallinus |
| 174 | TRINITY_DN9846_c0_g1_i3 | 57946 | 3 | 6.569 | β-bungarotoxin | | | JAA74785 | Furina ornata |
| 175 | TRINITY_DN4685_c0_g1_i23 | 284 | 0 | 2.6072 | β-bungarotoxin | | 20.09% | ABG90492 | Bungarus multicinctus |
| 176 | TRINITY_DN4189_c0_g1_i6 | 229956 | 22 | 10.269 | β-bungarotoxin | | | AAL87003 | Bungarus caeruleus |
| 177 | TRINITY_DN374_c0_g1_i8 | 70499 | 1 | 11.932 | β-bungarotoxin | | | JAA74785 | Furina ornata |
| 178 | TRINITY_DN374_c0_g1_i17 | 55732 | 0 | 11.932 | β-bungarotoxin | | | JAA74785 | Furina ornata |
| 179 | TRINITY_DN374_c0_g1_i22 | 3123 | 0 | 11.932 | β-bungarotoxin | | | AAL30065 | Bungarus candidus |
| 180 | TRINITY_DN374_c0_g1_i10 | 241 | 12 | 11.932 | β-bungarotoxin | | | JAA74785 | Furina ornata |
| 181 | TRINITY_DN2983_c0_g1_i2 | 197 | 2 | 5.9443 | β-bungarotoxin | | | AAL30063 | Bungarus candidus |
| 182 | TRINITY_DN370_c0_g1_i7 | 3912 | 0 | 13.603 | β-bungarotoxin | | | CAA72809 | Bungarus multicinctus |

| Figure S6B. *Bungarus caeruleus* | | | | | | | | | |
|---|---|---|---|---|---|---|---|---|---|
| S. No. | Isoform | VG | Int | M | Family | Subtype | Abundance | Accession | Scientific name |
| 1 | TRINITY_DN770_c0_g1 | 5199 | 8 | 6.12009511 | 5'-nucleotidase | | 0.04% | XP_026581084 | Pseudonaja textilis |
| 2 | TRINITY_DN4162_c0_g1 | 415 | 27 | 2.338480665 | 5'-nucleotidase | | | XP_026539324 | Notechis scutatus |
| 3 | TRINITY_DN4162_c0_g1 | 238 | 0 | 2.338480665 | 5'-nucleotidase | | | XP_026539324 | Notechis scutatus |
| 4 | TRINITY_DN4162_c0_g1 | 218 | 125 | 2.338480665 | 5'-nucleotidase | | | XP_026539324 | Notechis scutatus |
| 5 | TRINITY_DN5084_c1_g2 | 204 | 39 | 2.000383847 | 5'-nucleotidase | | | XP_026522489 | Notechis scutatus |
| 6 | TRINITY_DN4162_c0_g1 | 176 | 18 | 2.338480665 | 5'-nucleotidase | | | XP_026539324 | Notechis scutatus |
| 7 | TRINITY_DN846_c0_g1 | 107953 | 22 | 14.70189849 | Acetylcholinesterase | | 1.28% | AAC59905 | Bungarus fasciatus |
| 8 | TRINITY_DN846_c0_g1 | 68234 | 3 | 14.70189849 | Acetylcholinesterase | | | AAC59905 | Bungarus fasciatus |
| 9 | TRINITY_DN846_c0_g1 | 31465 | 0 | 14.70189849 | Acetylcholinesterase | | | Q92035 | Bungarus fasciatus |
| 10 | TRINITY_DN846_c0_g1 | 744 | 0 | 14.70189849 | Acetylcholinesterase | | | Q92035 | Bungarus fasciatus |
| 11 | TRINITY_DN389_c1_g1 | 1202248 | 48 | 17.10596803 | CRISP | | 13.07% | ACE73577 | Bungarus candidus |
| 12 | TRINITY_DN389_c1_g1 | 918054 | 0 | 17.10596803 | CRISP | | | ACE73577 | Bungarus candidus |
| 13 | TRINITY_DN2703_c0_g1 | 10221 | 669 | 4.223205459 | CRISP | | | JAB52844 | Micrurus fulvius |
| 14 | TRINITY_DN5802_c0_g1 | 743 | 0 | 11.38622429 | CRISP | | | ACE73578 | Bungarus candidus |
| 15 | TRINITY_DN1897_c0_g1 | 676 | 0 | 13.5002171 | CRISP | | | XP_015678374 | Protobothrops mucrosquamatus |
| 16 | TRINITY_DN11613_c0_g1 | 410 | 0 | 10.51880993 | CRISP | | | AAP20603 | Naja atra |
| 17 | TRINITY_DN14181_c1_g1 | 237 | 8 | 5.476371925 | CRISP | | | XP_026539804 | Notechis scutatus |
| 18 | TRINITY_DN2322_c0_g1 | 219 | 0 | 2.755185109 | CRISP | | | LAB38794 | Micrurus spixii |
| 19 | TRINITY_DN140247_c0_g1 | 188 | 0 | 9.393918685 | CRISP | | | XP_026568632 | Pseudonaja textilis |
| 20 | TRINITY_DN676_c1_g2 | 1541 | 3 | 10.5291807 | CTL | | 0.05% | AAK43586 | Bungarus multicinctus |
| 21 | TRINITY_DN1049_c0_g1 | 1528 | 20 | 8.369519998 | CTL | | | ABP94094 | Oxyuranus scutellatus |
| 22 | TRINITY_DN1049_c0_g1 | 1213 | 0 | 8.369519998 | CTL | | | JAA74667 | Brachyurophis roperi |
| 23 | TRINITY_DN676_c1_g2 | 1020 | 0 | 10.5291807 | CTL | | | ABH05181 | Bungarus multicinctus |
| 24 | TRINITY_DN676_c1_g2 | 629 | 0 | 10.5291807 | CTL | | | AAK43586 | Bungarus multicinctus |
| 25 | TRINITY_DN676_c1_g2 | 496 | 0 | 10.5291807 | CTL | | | AAK43586 | Bungarus multicinctus |
| 26 | TRINITY_DN1049_c0_g1 | 458 | 7 | 8.369519998 | CTL | | | ACC67942 | Oxyuranus microlepidotus |
| 27 | TRINITY_DN4221_c0_g1 | 350 | 44 | 4.666871011 | CTL | | | XP_026577709 | Pseudonaja textilis |
| 28 | TRINITY_DN1049_c0_g1 | 254 | 0 | 8.369519998 | CTL | | | ABU68499 | Leioheterodon madagascariensis |
| 29 | TRINITY_DN1049_c0_g1 | 128 | 0 | 8.369519998 | CTL | | | JAC95003 | Pantherophis guttatus |
| 30 | TRINITY_DN541_c0_g1 | 163938 | 0 | 16.49630974 | Disintegrin-like | | 2.52% | ABQ01132 | Tropidechis carinatus |
| 31 | TRINITY_DN541_c0_g1 | 67618 | 0 | 16.49630974 | Disintegrin-like | | | ABN72537 | Bungarus multicinctus |
| 32 | TRINITY_DN541_c0_g1 | 43227 | 7 | 16.49630974 | Disintegrin-like | | | ABN72537 | Bungarus multicinctus |
| 33 | TRINITY_DN541_c0_g1 | 40431 | 2 | 16.49630974 | Disintegrin-like | | | ABN72537 | Bungarus multicinctus |
| 34 | TRINITY_DN541_c0_g1 | 39001 | 2 | 16.49630974 | Disintegrin-like | | | ABN72537 | Bungarus multicinctus |
| 35 | TRINITY_DN1050_c0_g1 | 18795 | 3 | 12.89954547 | Disintegrin-like | | | ABN72537 | Bungarus multicinctus |
| 36 | TRINITY_DN541_c0_g1 | 8597 | 0 | 16.49630974 | Disintegrin-like | | | LAB50889 | Micrurus surinamensis |
| 37 | TRINITY_DN541_c0_g1 | 8254 | 0 | 16.49630974 | Disintegrin-like | | | ABN72537 | Bungarus multicinctus |
| 38 | TRINITY_DN17326_c0_g1 | 6890 | 0 | 14.64265375 | Disintegrin-like | | | JAA74814 | Hemiaspis signata |
| 39 | TRINITY_DN541_c0_g1 | 3901 | 0 | 16.49630974 | Disintegrin-like | | | ABN72537 | Bungarus multicinctus |
| 40 | TRINITY_DN995_c0_g1 | 2987 | 0 | 7.945553716 | Disintegrin-like | | | ADI47635 | Echis coloratus |
| 41 | TRINITY_DN34742_c0_g1 | 2967 | 0 | 13.47096164 | Disintegrin-like | | | XP_032084682 | Thamnophis elegans |
| 42 | TRINITY_DN541_c0_g1 | 1369 | 2 | 16.49630974 | Disintegrin-like | | | ABN72537 | Bungarus multicinctus |
| 43 | TRINITY_DN5558_c0_g1 | 863 | 7 | 7.021288627 | Disintegrin-like | | | XP_007430976 | Python bivittatus |
| 44 | TRINITY_DN36697_c0_g4 | 387 | 0 | 5.357901342 | Disintegrin-like | | | JAA74866 | Hoplocephalus bungaroides |
| 45 | TRINITY_DN17326_c0_g1 | 231 | 0 | 14.64265375 | Disintegrin-like | | | JAA74814 | Hemiaspis signata |
| 46 | TRINITY_DN6269_c0_g1 | 202 | 98 | 2.849120917 | Disintegrin-like | | | XP_034287530 | Pantherophis guttatus |
| 47 | TRINITY_DN11657_c0_g1 | 201 | 0 | 8.356915558 | Disintegrin-like | | | XP_034276822 | Pantherophis guttatus |
| 48 | TRINITY_DN1050_c0_g1 | 197 | 0 | 12.89954547 | Disintegrin-like | | | LAB61872 | Micrurus surinamensis |
| 49 | TRINITY_DN5558_c0_g1 | 159 | 8 | 7.021288627 | Disintegrin-like | | | XP_013912183 | Thamnophis sirtalis |

| | | | | | | | | | |
|---|---|---|---|---|---|---|---|---|---|
| 50 | TRINITY_DN995_c0_g1 | 142 | 0 | 7.945553716 | Disintegrin-like | | | JAS05092 | Micrurus tener |
| 51 | TRINITY_DN995_c0_g1 | 112 | 0 | 7.945553716 | Disintegrin-like | | | ABK63559 | Demansia vestigiata |
| 52 | TRINITY_DN6561_c0_g2 | 108 | 54 | 2.822845138 | Disintegrin-like | | | XP_026538843 | Notechis scutatus |
| 53 | TRINITY_DN5558_c0_g1 | 108 | 0 | 7.021288627 | Disintegrin-like | | | XP_013912183 | Thamnophis sirtalis |
| 54 | TRINITY_DN995_c0_g1 | 102 | 0 | 7.945553716 | Disintegrin-like | | | ADI47635 | Echis coloratus |
| 55 | TRINITY_DN7688_c0_g1 | 6890 | 0 | 2.192026844 | DPP | | | XP_026530199 | Notechis scutatus |
| 56 | TRINITY_DN7688_c0_g1 | 2576 | 0 | 2.192026844 | DPP | | 0.06% | ABQ63098 | Oxyuranus scutellatus |
| 57 | TRINITY_DN7688_c0_g1 | 603 | 90 | 2.192026844 | DPP | | | ABQ63103 | Pseudechis australis |
| 58 | TRINITY_DN7454_c0_g1 | 42758 | 0 | 14.67713735 | Hyaluronidase | | | XP_026524834 | Notechis scutatus |
| 59 | TRINITY_DN7454_c0_g1 | 19050 | 0 | 14.67713735 | Hyaluronidase | | | XP_034277309 | Pantherophis guttatus |
| 60 | TRINITY_DN7454_c0_g1 | 1270 | 0 | 14.67713735 | Hyaluronidase | | 0.40% | XP_034277309 | Pantherophis guttatus |
| 61 | TRINITY_DN7454_c0_g1 | 1187 | 0 | 14.67713735 | Hyaluronidase | | | XP_026524834 | Notechis scutatus |
| 62 | TRINITY_DN7454_c0_g1 | 551 | 0 | 14.67713735 | Hyaluronidase | | | XP_026524834 | Notechis scutatus |
| 63 | TRINITY_DN7454_c0_g1 | 496 | 0 | 14.67713735 | Hyaluronidase | | | LAA86771 | Micrurus lemniscatus |
| 64 | TRINITY_DN567_c0_g1 | 1282916 | 0 | 13.81824703 | Kunitz | | | ACR78501 | Drysdalia coronoides |
| 65 | TRINITY_DN9666_c0_g1 | 161168 | 0 | 19.13753563 | Kunitz | | | XP_026579466 | Pseudonaja textilis |
| 66 | TRINITY_DN1301_c0_g1 | 6801 | 0 | 2.848076319 | Kunitz | | | XP_026543549 | Notechis scutatus |
| 67 | TRINITY_DN389_c1_g2 | 3686 | 0 | 13.69341901 | Kunitz | | 8.94% | ABN72545 | Bungarus multicinctus |
| 68 | TRINITY_DN2477_c0_g1 | 2300 | 1513 | 2.201361066 | Kunitz | | | XP_026520368 | Notechis scutatus |
| 69 | TRINITY_DN567_c0_g1 | 892 | 0 | 13.81824703 | Kunitz | | | ACR78501 | Drysdalia coronoides |
| 70 | TRINITY_DN567_c0_g1 | 234 | 0 | 13.81824703 | Kunitz | | | LAA53789 | Micrurus corallinus |
| 71 | TRINITY_DN567_c0_g1 | 208 | 0 | 13.81824703 | Kunitz | | | LAA53789 | Micrurus corallinus |
| 72 | TRINITY_DN482_c0_g1 | 112710 | 0 | 11.91885276 | LAAO | | | ABN72539 | Bungarus multicinctus |
| 73 | TRINITY_DN482_c0_g1 | 51876 | 0 | 11.91885276 | LAAO | | | ABN72540 | Bungarus fasciatus |
| 74 | TRINITY_DN482_c0_g1 | 36169 | 0 | 11.91885276 | LAAO | | | ABN72539 | Bungarus multicinctus |
| 75 | TRINITY_DN482_c0_g1 | 31413 | 0 | 11.91885276 | LAAO | | | ABN72539 | Bungarus multicinctus |
| 76 | TRINITY_DN482_c0_g1 | 25309 | 0 | 11.91885276 | LAAO | | | ABN72539 | Bungarus multicinctus |
| 77 | TRINITY_DN482_c0_g1 | 15695 | 0 | 11.91885276 | LAAO | | | ABN72539 | Bungarus multicinctus |
| 78 | TRINITY_DN482_c0_g1 | 2605 | 6 | 11.91885276 | LAAO | | 1.72% | ABN72540 | Bungarus fasciatus |
| 79 | TRINITY_DN482_c0_g1 | 1701 | 0 | 11.91885276 | LAAO | | | ABN72539 | Bungarus multicinctus |
| 80 | TRINITY_DN8453_c0_g1 | 1018 | 0 | 12.2197909 | LAAO | | | JAA74636 | Acanthophis wellsi |
| 81 | TRINITY_DN5468_c0_g1 | 585 | 0 | 9.800504988 | LAAO | | | XP_026549343 | Notechis scutatus |
| 82 | TRINITY_DN5468_c0_g1 | 364 | 0 | 9.800504988 | LAAO | | | XP_026549343 | Notechis scutatus |
| 83 | TRINITY_DN8453_c0_g1 | 315 | 0 | 12.2197909 | LAAO | | | ABN72539 | Bungarus multicinctus |
| 84 | TRINITY_DN5468_c0_g1 | 168 | 0 | 9.800504988 | LAAO | | | XP_026549343 | Notechis scutatus |
| 85 | TRINITY_DN7590_c3_g1 | 105 | 0 | 8.553575351 | LAAO | | | ABN72540 | Bungarus fasciatus |
| 86 | TRINITY_DN450_c0_g1 | 441365 | 0 | 11.86948642 | Natriuretic peptide | | | ADK12002 | Ophiophagus hannah |
| 87 | TRINITY_DN450_c0_g1 | 123771 | 0 | 11.86948642 | Natriuretic peptide | | 3.59% | ADK12002 | Ophiophagus hannah |
| 88 | TRINITY_DN450_c0_g1 | 20717 | 500 | 11.86948642 | Natriuretic peptide | | | JAS05144 | Micrurus tener |
| 89 | TRINITY_DN14642_c0_g1 | 122 | 0 | 8.816609757 | Natriuretic peptide | | | ADK12002 | Ophiophagus hannah |
| 90 | TRINITY_DN676_c2_g1 | 52057 | 10 | 7.763338989 | NGF | | | AAB25729 | Bungarus multicinctus |
| 91 | TRINITY_DN676_c2_g1 | 2092 | 828 | 7.763338989 | NGF | | | ABA60118 | Pseudonaja textilis |
| 92 | TRINITY_DN676_c2_g1 | 1839 | 0 | 7.763338989 | NGF | | | AAB25729 | Bungarus multicinctus |
| 93 | TRINITY_DN676_c2_g1 | 611 | 0 | 7.763338989 | NGF | | 0.35% | AAB25729 | Bungarus multicinctus |
| 94 | TRINITY_DN676_c2_g1 | 296 | 0 | 7.763338989 | NGF | | | AAB25729 | Bungarus multicinctus |
| 95 | TRINITY_DN2739_c1_g1 | 238 | 9 | 4.460457436 | NGF | | | ETE72246 | Ophiophagus hannah |
| 96 | TRINITY_DN676_c2_g1 | 121 | 0 | 7.763338989 | NGF | | | AAB25729 | Bungarus multicinctus |
| 97 | TRINITY_DN117_c4_g1 | 1317114 | 0 | 13.23957567 | PLA2 | | | ADF50037 | Bungarus flaviceps |
| 98 | TRINITY_DN117_c4_g1 | 1210811 | 0 | 13.23957567 | PLA2 | | | ACY68711 | Parasuta nigriceps |
| 99 | TRINITY_DN117_c4_g1 | 821053 | 0 | 13.23957567 | PLA2 | | | AAB24834 | Bungarus fasciatus |

| | | | | | | | | | |
|---|---|---|---|---|---|---|---|---|---|
| 100 | TRINITY_DN491_c1_g1 | 625587 | 0 | 21.09418084 | PLA2 | | | BAC77655 | Bungarus flaviceps |
| 101 | TRINITY_DN117_c4_g1 | 31520 | 0 | 13.23957567 | PLA2 | | | ADF50037 | Bungarus flaviceps |
| 102 | TRINITY_DN117_c4_g1 | 18769 | 2 | 13.23957567 | PLA2 | | **24.71%** | ACY68711 | Parasuta nigriceps |
| 103 | TRINITY_DN117_c4_g1 | 5123 | 0 | 13.23957567 | PLA2 | | | ACY68711 | Parasuta nigriceps |
| 104 | TRINITY_DN117_c4_g1 | 1819 | 0 | 13.23957567 | PLA2 | | | ADF50037 | Bungarus flaviceps |
| 105 | TRINITY_DN117_c4_g1 | 721 | 0 | 13.23957567 | PLA2 | | | ADF50037 | Bungarus flaviceps |
| 106 | TRINITY_DN117_c4_g1 | 473 | 191 | 13.23957567 | PLA2 | | | AAB24834 | Bungarus fasciatus |
| 107 | TRINITY_DN7320_c0_g1 | 111 | 0 | 3.207965507 | PLA2 | | | LAA68819 | Micrurus lemniscatus |
| 108 | TRINITY_DN117_c4_g1 | 106 | 2 | 13.23957567 | PLA2 | | | ACY68711 | Parasuta nigriceps |
| 109 | TRINITY_DN839_c0_g1 | 22859 | 0 | 8.996722072 | PLB | | | ACR78473 | Drysdalia coronoides |
| 110 | TRINITY_DN839_c0_g1 | 19491 | 29 | 8.996722072 | PLB | | | XP_026553321 | Pseudonaja textilis |
| 111 | TRINITY_DN839_c0_g1 | 16903 | 289 | 8.996722072 | PLB | | **0.38%** | ACR78473 | Drysdalia coronoides |
| 112 | TRINITY_DN839_c0_g1 | 2787 | 6 | 8.996722072 | PLB | | | ACR78473 | Drysdalia coronoides |
| 113 | TRINITY_DN839_c0_g1 | 310 | 27 | 8.996722072 | PLB | | | ACR78473 | Drysdalia coronoides |
| 114 | TRINITY_DN839_c0_g1 | 152 | 0 | 8.996722072 | PLB | | | ACR78473 | Drysdalia coronoides |
| 115 | TRINITY_DN3739_c0_g1 | 23304 | 0 | 6.272974363 | Serine Protease | | **0.14%** | JAB54462 | Micrurus fulvius |
| 116 | TRINITY_DN9079_c4_g1 | 100 | 0 | 4.318947707 | Serine Protease | | | XP_026562705 | Pseudonaja textilis |
| 117 | TRINITY_DN7325_c0_g1 | 285459 | 0 | 22.11784508 | Type-I α-neurotoxin | | | CAA45882 | Bungarus multicinctus |
| 118 | TRINITY_DN7325_c0_g1 | 262 | 0 | 22.11784508 | Type-I α-neurotoxin | | | CAA45882 | Bungarus multicinctus |
| 119 | TRINITY_DN323_c0_g1 | 423609 | 0 | 18.75215495 | Type-I α-neurotoxin | | | CAA06887 | Bungarus multicinctus multicinctus |
| 120 | TRINITY_DN323_c0_g1 | 94604 | 0 | 18.75215495 | Type-I α-neurotoxin | | | CAA06887 | Bungarus multicinctus multicinctus |
| 121 | TRINITY_DN9940_c0_g1 | 39442 | 0 | 16.07062565 | Type-I α-neurotoxin | | | ETE58964 | Ophiophagus hannah |
| 122 | TRINITY_DN323_c0_g1 | 17856 | 0 | 18.75215495 | Type-I α-neurotoxin | | | CAA06887 | Bungarus multicinctus multicinctus |
| 123 | TRINITY_DN323_c0_g1 | 7722 | 0 | 18.75215495 | Type-I α-neurotoxin | | | CAA06887 | Bungarus multicinctus multicinctus |
| 124 | TRINITY_DN1005_c0_g1 | 1085 | 0 | 6.560991216 | Type-I α-neurotoxin | | | P82462 | Ophiophagus hannah |
| 125 | TRINITY_DN1005_c0_g1 | 691 | 0 | 6.560991216 | Type-I α-neurotoxin | | | P82462 | Ophiophagus hannah |
| 126 | TRINITY_DN1005_c0_g1 | 258 | 0 | 6.560991216 | Type-I α-neurotoxin | | | P82462 | Ophiophagus hannah |
| 127 | TRINITY_DN1005_c0_g1 | 140 | 0 | 6.560991216 | Type-I α-neurotoxin | | | P82462 | Ophiophagus hannah |
| 128 | TRINITY_DN323_c0_g1 | 1018919 | 0 | 18.75215495 | Type-II α-neurotoxin | | | ETE56306 | Ophiophagus hannah |
| 129 | TRINITY_DN1485_c0_g1 | 975145 | 0 | 21.29056926 | Type-II α-neurotoxin | | | CAJ77819 | Bungarus candidus |
| 130 | TRINITY_DN1364_c0_g1 | 367500 | 0 | 20.3818592 | Type-II α-neurotoxin | **61.82%** | | LAA95127 | Micrurus lemniscatus lemniscatus |
| 131 | TRINITY_DN1364_c0_g1 | 945 | 0 | 20.3818592 | Type-II α-neurotoxin | | | LAA95127 | Micrurus lemniscatus lemniscatus |
| 132 | TRINITY_DN45054_c0_g1 | 527 | 0 | 12.43272096 | Type-II α-neurotoxin | | | ABK63538 | Tropidechis carinatus |
| 133 | TRINITY_DN1364_c0_g1 | 423 | 0 | 20.3818592 | Type-II α-neurotoxin | | | LAA95127 | Micrurus lemniscatus lemniscatus |
| 134 | TRINITY_DN1364_c0_g1 | 3910 | 0 | 20.3818592 | Type-II α-neurotoxin | | | XP_026559448 | Pseudonaja textilis |
| 135 | TRINITY_DN1364_c0_g1 | 2291 | 0 | 20.3818592 | Type-II α-neurotoxin | | | ETE56205 | Ophiophagus hannah |
| 136 | TRINITY_DN2007_c2_g1 | 891 | 0 | 13.08488254 | Type-II α-neurotoxin | | **32.15%** | ETE56205 | Ophiophagus hannah |
| 137 | TRINITY_DN2007_c2_g1 | 581 | 0 | 13.08488254 | Type-II α-neurotoxin | | | ETE56205 | Ophiophagus hannah |
| 138 | TRINITY_DN1364_c0_g2 | 478 | 0 | 19.29174805 | Type-II α-neurotoxin | | | AAL30059 | Bungarus candidus |
| 139 | TRINITY_DN7325_c0_g1 | 330 | 0 | 22.11784508 | Type-II α-neurotoxin | | | ETE56205 | Ophiophagus hannah |
| 140 | TRINITY_DN2007_c2_g1 | 301 | 0 | 13.08488254 | Type-II α-neurotoxin | | | ETE56205 | Ophiophagus hannah |
| 141 | TRINITY_DN6859_c0_g1 | 236 | 0 | 10.77596777 | Type-II α-neurotoxin | | | ETE56205 | Ophiophagus hannah |
| 142 | TRINITY_DN7325_c0_g1 | 224 | 0 | 22.11784508 | Type-II α-neurotoxin | | | ETE56205 | Ophiophagus hannah |
| 143 | TRINITY_DN7325_c0_g1 | 148 | 0 | 22.11784508 | Type-II α-neurotoxin | | | ETE56205 | Ophiophagus hannah |
| 144 | TRINITY_DN1485_c0_g1 | 825527 | 0 | 21.29056926 | Unconventional 3FTx | | | CAA06885 | Bungarus multicinctus |
| 145 | TRINITY_DN323_c0_g1 | 109907 | 0 | 18.75215495 | Unconventional 3FTx | | | CAB53358 | Bungarus multicinctus |
| 146 | TRINITY_DN9940_c0_g1 | 9212 | 4 | 16.07062565 | Unconventional 3FTx | **18.01%** | | LAB55960 | Micrurus surinamensis |
| 147 | TRINITY_DN1485_c2_g2 | 277 | 0 | 10.10142468 | Unconventional 3FTx | | | XP_032078231 | Thamnophis elegans |
| 148 | TRINITY_DN24809_c0_g1 | 242 | 0 | 9.75819307 | Unconventional 3FTx | | | CAD56381 | Bungarus multicinctus |
| 149 | TRINITY_DN1485_c0_g1 | 1015208 | 0 | 21.29056926 | κ-bungarotoxin | | | CAA35775 | Bungarus multicinctus |

| | | | | | | | | | |
|---|---|---|---|---|---|---|---|---|---|
| 150 | TRINITY_DN9940_c0_g2 | 25651 | 0 | 16.48605728 | κ-bungarotoxin | | | AAL30054 | Bungarus candidus |
| 151 | TRINITY_DN1485_c0_g1 | 11382 | 0 | 21.29056926 | κ-bungarotoxin | | | CAB46659 | Bungarus multicinctus |
| 152 | TRINITY_DN1485_c0_g1 | 4741 | 0 | 21.29056926 | κ-bungarotoxin | **20.17%** | | CAB46659 | Bungarus multicinctus |
| 153 | TRINITY_DN1485_c0_g1 | 869 | 0 | 21.29056926 | κ-bungarotoxin | | | CAB46659 | Bungarus multicinctus |
| 154 | TRINITY_DN2007_c2_g1 | 376 | 0 | 13.08488254 | κ-bungarotoxin | | | CAA35775 | Bungarus multicinctus |
| 155 | TRINITY_DN2007_c2_g1 | 117 | 0 | 13.08488254 | κ-bungarotoxin | | | CAA35775 | Bungarus multicinctus |
| 156 | TRINITY_DN66_c0_g1 | 122728 | 16 | 9.076857398 | VEGF | | | JAA74898 | Pseudonaja modesta |
| 157 | TRINITY_DN66_c0_g1 | 33393 | 0 | 9.076857398 | VEGF | | | XP_026551346 | Pseudonaja textilis |
| 158 | TRINITY_DN66_c0_g1 | 12007 | 11 | 9.076857398 | VEGF | | | XP_026551344 | Pseudonaja textilis |
| 159 | TRINITY_DN14480_c0_g1 | 6882 | 56 | 6.555882396 | VEGF | | | LAB17921 | Micrurus spixii |
| 160 | TRINITY_DN66_c0_g1 | 3766 | 0 | 9.076857398 | VEGF | | | XP_026551346 | Pseudonaja textilis |
| 161 | TRINITY_DN66_c0_g1 | 1699 | 0 | 9.076857398 | VEGF | | | XP_026551346 | Pseudonaja textilis |
| 162 | TRINITY_DN66_c0_g1 | 1617 | 0 | 9.076857398 | VEGF | | **1.12%** | XP_026551346 | Pseudonaja textilis |
| 163 | TRINITY_DN12448_c0_g1 | 256 | 0 | 6.502128806 | VEGF | | | XP_026558913 | Pseudonaja textilis |
| 164 | TRINITY_DN14480_c0_g1 | 228 | 0 | 6.555882396 | VEGF | | | LAB17921 | Micrurus spixii |
| 165 | TRINITY_DN12959_c0_g4 | 161 | 14 | 5.198653474 | VEGF | | | LAA79011 | Micrurus lemniscatus |
| 166 | TRINITY_DN14480_c0_g1 | 143 | 0 | 6.555882396 | VEGF | | | XP_026555459 | Pseudonaja textilis |
| 167 | TRINITY_DN12448_c0_g1 | 113 | 13 | 6.502128806 | VEGF | | | XP_026558913 | Pseudonaja textilis |
| 168 | TRINITY_DN1325_c0_g1 | 110738 | 0 | 18.62453962 | Vespryn | | **0.68%** | XP_026576508 | Pseudonaja textilis |
| 169 | TRINITY_DN1076_c0_g2 | 2862 | 0 | 13.32867607 | β-bungarotoxin | | | AAL30067.1 | Furina ornata |
| 170 | TRINITY_DN768_c1_g1 | 710088 | 0 | 9.915452031 | β-bungarotoxin | | | BAD06267 | Bungarus candidus |
| 171 | TRINITY_DN768_c1_g1 | 389230 | 0 | 9.915452031 | β-bungarotoxin | | | BAD06267 | Bungarus candidus |
| 172 | TRINITY_DN768_c1_g1 | 200648 | 492 | 9.915452031 | β-bungarotoxin | | **8.80%** | BAD06267 | Bungarus candidus |
| 173 | TRINITY_DN567_c0_g1 | 112704 | 27 | 13.81824703 | β-bungarotoxin | | | AAL30065.1 | Furina ornata |
| 174 | TRINITY_DN11210_c1_g1 | 18577 | 0 | 11.61691136 | β-bungarotoxin | | | CAA72809 | Bungarus multicinctus |
| 175 | TRINITY_DN4436_c0_g1 | 1861 | 0 | 14.57255705 | β-bungarotoxin | | | CAB62503 | Bungarus multicinctus |

| Figure S6C. *Bungarus romulusi* | | | | | | | | | |
|---|---|---|---|---|---|---|---|---|---|
| S. No. | Isoform | VG | Int | M | Family | Subtype | Abundance | Accession | Scientific name |
| 1 | TRINITY_DN2049_c0_g2_i4 | 31860 | 304 | 5.235644987 | 5'-nucleotidase | | 0.44% | JAB52815 | Micrurus fulvius |
| 2 | TRINITY_DN19829_c0_g1_i3 | 135 | 23 | 5.235644987 | 5'-nucleotidase | | | XP_026522489 | Notechis scutatus |
| 3 | TRINITY_DN2049_c0_g2_i1 | 103 | 2 | 13.57174515 | 5'-nucleotidase | | | XP_026582322 | Pseudonaja textilis |
| 4 | TRINITY_DN359_c0_g1_i12 | 45074 | 0 | 13.57174515 | Acetylcholinesterase | | 0.96% | AAC59905 | Bungarus fasciatus |
| 5 | TRINITY_DN359_c0_g1_i3 | 20202 | 0 | 13.57174515 | Acetylcholinesterase | | | AAC59905 | Bungarus fasciatus |
| 6 | TRINITY_DN359_c0_g1_i9 | 2177 | 8 | 13.57174515 | Acetylcholinesterase | | | AAC59905 | Bungarus fasciatus |
| 7 | TRINITY_DN359_c0_g1_i7 | 1295 | 0 | 13.57174515 | Acetylcholinesterase | | | AAC59905 | Bungarus fasciatus |
| 8 | TRINITY_DN359_c0_g1_i1 | 1261 | 0 | 13.57174515 | Acetylcholinesterase | | | Q92035 | Bungarus fasciatus |
| 9 | TRINITY_DN359_c0_g1_i6 | 884 | 0 | 2.867876439 | Acetylcholinesterase | | | AAC59905 | Bungarus fasciatus |
| 10 | TRINITY_DN356_c3_g1_i2 | 42031 | 7828 | 3.367052312 | Calreticulin | | 0.57% | JAB54560 | Micrurus fulvius |
| 11 | TRINITY_DN49_c0_g1_i3 | 361934 | 10 | 3.367052312 | CRISP | | 9.67% | ACE73577 | Bungarus candidus |
| 12 | TRINITY_DN49_c0_g1_i4 | 344200 | 0 | 3.367052312 | CRISP | | | ACE73577 | Bungarus candidus |
| 13 | TRINITY_DN49_c0_g1_i6 | 5624 | 0 | 3.367052312 | CRISP | | | ACE73577 | Bungarus candidus |
| 14 | TRINITY_DN49_c0_g1_i7 | 567 | 0 | 3.367052312 | CRISP | | | ACE73577 | Bungarus candidus |
| 15 | TRINITY_DN49_c0_g1_i12 | 491 | 6 | 3.367052312 | CRISP | | | ACE73577 | Bungarus candidus |
| 16 | TRINITY_DN49_c0_g1_i1 | 236 | 0 | 10.74274075 | CRISP | | | ACE73577 | Bungarus candidus |
| 17 | TRINITY_DN755_c0_g2_i1 | 12517 | 10 | 15.13281705 | CTL | | 0.39% | XP_026579605 | Pseudonaja textilis |
| 18 | TRINITY_DN755_c0_g1_i2 | 10401 | 0 | 15.13281705 | CTL | | | ABP94094 | Oxyuranus scutellatus |
| 19 | TRINITY_DN755_c0_g1_i8 | 1665 | 0 | 15.13281705 | CTL | | | ABU68499 | Leioheterodon madagascariensis |
| 20 | TRINITY_DN755_c0_g1_i3 | 1340 | 0 | 15.13281705 | CTL | | | ABP94094 | Oxyuranus scutellatus |
| 21 | TRINITY_DN755_c0_g1_i13 | 1045 | 0 | 6.500581099 | CTL | | | JAC95003 | Pantherophis guttatus |
| 22 | TRINITY_DN1070_c0_g2_i2 | 450 | 7 | 9.111279121 | CTL | | | ADF50042 | Bungarus flaviceps |
| 23 | TRINITY_DN21017_c0_g1_i2 | 380 | 1 | 15.13281705 | CTL | | | XP_026527775 | Notechis scutatus |
| 24 | TRINITY_DN755_c0_g1_i11 | 271 | 0 | 15.13281705 | CTL | | | ABP94094 | Oxyuranus scutellatus |
| 25 | TRINITY_DN755_c0_g1_i5 | 260 | 0 | 2.575058162 | CTL | | | ABP94094 | Oxyuranus scutellatus |
| 26 | TRINITY_DN732_c0_g1_i8 | 181 | 74 | 8.410294102 | CTL | | | XP_026546618 | Notechis scutatus |
| 27 | TRINITY_DN1070_c0_g3_i2 | 144 | 0 | 15.13281705 | CTL | | | ABP94128 | Pseudechis porphyriacus |
| 28 | TRINITY_DN755_c0_g1_i7 | 132 | 0 | 10.61063388 | CTL | | | ABP94118 | Notechis scutatus |
| 29 | TRINITY_DN307_c0_g1_i10 | 96468 | 0 | 10.61063388 | Disintegrin-like | | 5.60% | ABN72537 | Bungarus multicinctus |
| 30 | TRINITY_DN307_c0_g1_i8 | 56832 | 0 | 16.94513734 | Disintegrin-like | | | ABN72537 | Bungarus multicinctus |
| 31 | TRINITY_DN863_c0_g1_i1 | 49210 | 0 | 10.61063388 | Disintegrin-like | | | ABN72537 | Bungarus multicinctus |
| 32 | TRINITY_DN307_c0_g1_i6 | 44473 | 0 | 10.61063388 | Disintegrin-like | | | ABN72537 | Bungarus multicinctus |
| 33 | TRINITY_DN307_c0_g1_i13 | 41782 | 0 | 10.61063388 | Disintegrin-like | | | ABN72537 | Bungarus multicinctus |
| 34 | TRINITY_DN307_c0_g1_i12 | 30232 | 0 | 10.61063388 | Disintegrin-like | | | ABQ01136 | Oxyuranus scutellatus |
| 35 | TRINITY_DN307_c0_g1_i9 | 27679 | 0 | 10.61063388 | Disintegrin-like | | | ABN72537 | Bungarus multicinctus |
| 36 | TRINITY_DN307_c0_g1_i5 | 26854 | 0 | 14.95171343 | Disintegrin-like | | | ABN72537 | Bungarus multicinctus |
| 37 | TRINITY_DN474_c0_g3_i1 | 10637 | 0 | 14.54386448 | Disintegrin-like | | | ABQ01138 | Notechis scutatus |
| 38 | TRINITY_DN2689_c0_g1_i2 | 8523 | 0 | 10.61063388 | Disintegrin-like | | | ABQ01138 | Notechis scutatus |
| 39 | TRINITY_DN307_c0_g1_i7 | 4275 | 287 | 16.94513734 | Disintegrin-like | | | ABQ01136 | Oxyuranus scutellatus |
| 40 | TRINITY_DN863_c0_g1_i3 | 4091 | 0 | 13.207235 | Disintegrin-like | | | ABN72536 | Bungarus fasciatus |
| 41 | TRINITY_DN1500_c0_g1_i1 | 4003 | 0 | 14.95171343 | Disintegrin-like | | | ABN72537 | Bungarus multicinctus |
| 42 | TRINITY_DN474_c0_g3_i2 | 2776 | 0 | 12.07484015 | Disintegrin-like | | | AAM51550 | Naja mossambica |
| 43 | TRINITY_DN16430_c0_g3_i1 | 1826 | 0 | 11.79591687 | Disintegrin-like | | | AXL96555 | Borikenophis portoricensis |
| 44 | TRINITY_DN16430_c0_g2_i1 | 1505 | 0 | 6.7682172 | Disintegrin-like | | | ABN72537 | Bungarus multicinctus |
| 45 | TRINITY_DN10280_c0_g1_i5 | 492 | 8 | 10.61063388 | Disintegrin-like | | | XP_026536946 | Notechis scutatus |
| 46 | TRINITY_DN307_c0_g1_i2 | 290 | 0 | 5.25304303 | Disintegrin-like | | | ABN72537 | Bungarus multicinctus |
| 47 | TRINITY_DN2668_c0_g1_i3 | 186 | 125 | 5.020752147 | Disintegrin-like | | | XP_013912183 | Thamnophis sirtalis |
| 48 | TRINITY_DN99187_c0_g1_i1 | 166 | 7 | 6.430347561 | Disintegrin-like | | | XP_029141633 | Protobothrops mucrosquamatus |
| 49 | TRINITY_DN863_c0_g3_i1 | 126 | 2 | 16.94513734 | Disintegrin-like | | | XP_013922279 | Thamnophis sirtalis |

| | | | | | | | | | |
|---|---|---|---|---|---|---|---|---|---|
| 50 | TRINITY_DN863_c0_g1_i2 | 107 | 0 | 7.940808819 | Disintegrin-like | | | ABN72537 | Bungarus multicinctus |
| 51 | TRINITY_DN9869_c0_g1_i1 | 104 | 0 | 17.41398768 | Disintegrin-like | | | XP_026527224 | Notechis scutatus |
| 52 | TRINITY_DN72_c0_g4_i7 | 72237 | 0 | 17.19726702 | Hyaluronidase | | **0.98%** | XP_026524834 | Notechis scutatus |
| 53 | TRINITY_DN57_c15_g1_i1 | 37682 | 0 | 17.19726702 | Kunitz | | | ACR78501 | Drysdalia coronoides |
| 54 | TRINITY_DN57_c15_g1_i2 | 24636 | 0 | 15.11285161 | Kunitz | | **1.05%** | ACR78501 | Drysdalia coronoides |
| 55 | TRINITY_DN7705_c0_g1_i1 | 14998 | 0 | 16.05485143 | Kunitz | | | ACR78501 | Drysdalia coronoides |
| 56 | TRINITY_DN95585_c0_g1_i1 | 165 | 153 | 13.78907201 | Kunitz | | | XP_026579404 | Pseudonaja textilis |
| 57 | TRINITY_DN830_c0_g1_i13 | 58549 | 2 | 13.78907201 | **LAAO** | | | ABN72539 | Bungarus multicinctus |
| 58 | TRINITY_DN830_c0_g1_i12 | 48369 | 0 | 13.78907201 | **LAAO** | | | ABN72539 | Bungarus multicinctus |
| 59 | TRINITY_DN830_c0_g1_i3 | 33498 | 0 | 13.78907201 | **LAAO** | | | ABN72539 | Bungarus multicinctus |
| 60 | TRINITY_DN830_c0_g1_i22 | 6188 | 0 | 13.78907201 | **LAAO** | | | ABN72539 | Bungarus multicinctus |
| 61 | TRINITY_DN830_c0_g1_i7 | 2554 | 0 | 13.78907201 | **LAAO** | | **2.10%** | ABN72539 | Bungarus multicinctus |
| 62 | TRINITY_DN830_c0_g1_i6 | 2385 | 0 | 13.78907201 | **LAAO** | | | ABN72539 | Bungarus multicinctus |
| 63 | TRINITY_DN830_c0_g1_i14 | 1797 | 0 | 13.78907201 | **LAAO** | | | ABN72539 | Bungarus multicinctus |
| 64 | TRINITY_DN830_c0_g1_i19 | 1420 | 0 | 13.78907201 | **LAAO** | | | ABN72539 | Bungarus multicinctus |
| 65 | TRINITY_DN830_c0_g1_i17 | 194 | 1 | 13.78907201 | **LAAO** | | | ABN72539 | Bungarus multicinctus |
| 66 | TRINITY_DN830_c0_g1_i9 | 119 | 0 | 7.118403555 | **LAAO** | | | ABN72539 | Bungarus multicinctus |
| 67 | TRINITY_DN397_c1_g4_i1 | 3451 | 34 | 8.104433595 | Natriuretic peptide | | | JAS05144 | Micrurus tener |
| 68 | TRINITY_DN57_c0_g1_i16 | 103484 | 0 | 8.104433595 | Natriuretic peptide | | **1.49%** | ADK12002 | Ophiophagus hannah |
| 69 | TRINITY_DN57_c0_g1_i7 | 2605 | 10 | 9.488296614 | Natriuretic peptide | | | ADK12002 | Ophiophagus hannah |
| 70 | TRINITY_DN8160_c1_g1_i1 | 304 | 0 | 12.42271112 | Natriuretic peptide | | | ADK12002 | Ophiophagus hannah |
| 71 | TRINITY_DN4045_c0_g3_i1 | 27478 | 7 | 6.136764092 | **NGF** | | **0.38%** | AAB25729 | Bungarus multicinctus |
| 72 | TRINITY_DN1317_c1_g3_i1 | 257 | 5 | 19.70949826 | **NGF** | | | ETE59033 | Ophiophagus hannah |
| 73 | TRINITY_DN744_c0_g1_i2 | 1.00E+06 | 1 | 21.27715236 | PLA2 | | | 0702209A | Bungarus multicinctus |
| 74 | TRINITY_DN134_c0_g1_i7 | 373525 | 1 | 19.70949826 | PLA2 | | | AAR19228 | Bungarus caeruleus |
| 75 | TRINITY_DN744_c0_g1_i7 | 295169 | 0 | 21.27715236 | PLA2 | | | 0702209A | Bungarus multicinctus |
| 76 | TRINITY_DN134_c0_g1_i3 | 219678 | 0 | 19.70949826 | PLA2 | | | ADF50037 | Bungarus flaviceps |
| 77 | TRINITY_DN744_c0_g1_i4 | 187297 | 0 | 21.27715236 | PLA2 | | | CAA37482 | Bungarus multicinctus |
| 78 | TRINITY_DN134_c0_g1_i13 | 28675 | 0 | 15.91273919 | PLA2 | | | P0C551 | Bungarus candidus |
| 79 | TRINITY_DN11387_c0_g2_i1 | 26111 | 0 | 21.27715236 | PLA2 | | | CAA37482 | Bungarus multicinctus |
| 80 | TRINITY_DN134_c0_g1_i12 | 13816 | 0 | 21.27715236 | PLA2 | | | ADF50037 | Bungarus flaviceps |
| 81 | TRINITY_DN134_c0_g1_i2 | 8977 | 0 | 8.865637485 | PLA2 | | **29.42%** | BAD06270 | Bungarus candidus |
| 82 | TRINITY_DN602_c0_g1_i3 | 4618 | 0 | 13.26377694 | PLA2 | | | BAD06270 | Bungarus candidus |
| 83 | TRINITY_DN42_c0_g2_i1 | 4163 | 0 | 12.82720889 | PLA2 | | | ETE56084 | Ophiophagus hannah |
| 84 | TRINITY_DN5510_c0_g1_i1 | 3076 | 0 | 12.60996645 | PLA2 | | | AAO84769 | Bungarus candidus |
| 85 | TRINITY_DN42_c5_g1_i2 | 2621 | 0 | 21.27715236 | PLA2 | | | LAA68819 | Micrurus lemniscatus lemniscatus |
| 86 | TRINITY_DN134_c0_g1_i4 | 431 | 0 | 10.37222606 | PLA2 | | | ADF50037 | Bungarus flaviceps |
| 87 | TRINITY_DN7462_c0_g2_i1 | 360 | 0 | 10.37222606 | PLA2 | | | LAA39990 | Micrurus corallinus |
| 88 | TRINITY_DN7630_c0_g1_i3 | 166 | 101 | 6.765950593 | PLA2 | | | XP_026554534 | Pseudonaja textilis |
| 89 | TRINITY_DN40744_c0_g1_i1 | 159 | 2 | 9.034784967 | PLA2 | | | ETE56084 | Ophiophagus hannah |
| 90 | TRINITY_DN3776_c0_g1_i2 | 141 | 0 | 11.57910548 | PLA2 | | | 0702209A | Bungarus multicinctus |
| 91 | TRINITY_DN462_c0_g1_i1 | 143 | 0 | 12.45306764 | PLB | | **0.00%** | BAN82156 | Ovophis okinavensis |
| 92 | TRINITY_DN1122_c0_g2_i1 | 3945 | 0 | 12.45306764 | Serine protease | | **0.06%** | ABN72545 | Bungarus multicinctus |
| 93 | TRINITY_DN1122_c0_g2_i4 | 119 | 1 | 13.29021765 | Serine protease | | | ETE72364 | Ophiophagus hannah |
| 94 | TRINITY_DN0_c1_g3_i2 | 4177 | 0 | 6.42194512 | Type-I α-neurotoxin | | | JAB52865 | Micrurus fulvius |
| 95 | TRINITY_DN608_c1_g1_i4 | 362 | 0 | 18.52072304 | Type-I α-neurotoxin | | | P82462 | Naja atra |
| 96 | TRINITY_DN203_c3_g1_i1 | 158749 | 0 | 7.88101313 | Type-I α-neurotoxin | | | CAA06887 | Bungarus multicinctus multicinctus |
| 97 | TRINITY_DN89_c5_g1_i2 | 734221 | 1 | 10.25102086 | Type-II α-neurotoxin | | | ETE56306 | Ophiophagus hannah |
| 98 | TRINITY_DN9485_c1_g1_i5 | 311337 | 0 | 17.84029405 | Type-II α-neurotoxin | | | ETE56306 | Ophiophagus hannah |
| 99 | TRINITY_DN0_c1_g1_i6 | 246022 | 2 | 10.25102086 | Type-II α-neurotoxin | | | CAJ77818 | Bungarus candidus |

| | | | | | | | | | |
|---|---|---|---|---|---|---|---|---|---|
| 100 | TRINITY_DN9485_c1_g1_i1 | 163857 | 0 | 7.88101313 | Type-II α-neurotoxin | | | ETE56306 | Ophiophagus hannah |
| 101 | TRINITY_DN89_c5_g1_i5 | 99045 | 5 | 17.84029405 | Type-II α-neurotoxin | | | AHZ08825 | Micropechis ikaheca |
| 102 | TRINITY_DN0_c1_g1_i4 | 35852 | 0 | 15.76405356 | Type-II α-neurotoxin | **86.67%** | | CAJ77818 | Bungarus candidus |
| 103 | TRINITY_DN8116_c0_g2_i2 | 23553 | 0 | 7.88101313 | Type-II α-neurotoxin | | | ETE56306 | Ophiophagus hannah |
| 104 | TRINITY_DN89_c5_g1_i8 | 11451 | 0 | 17.84029405 | Type-II α-neurotoxin | | **35.90%** | AHZ08825 | Micropechis ikaheca |
| 105 | TRINITY_DN0_c1_g1_i5 | 9023 | 0 | 10.25102086 | Type-II α-neurotoxin | | | CAJ77818 | Bungarus candidus |
| 106 | TRINITY_DN9485_c1_g1_i2 | 4474 | 0 | 17.84029405 | Type-II α-neurotoxin | | | ETE56306 | Ophiophagus hannah |
| 107 | TRINITY_DN0_c1_g1_i1 | 1897 | 0 | 18.09317133 | Type-II α-neurotoxin | | | CAJ77818 | Bungarus candidus |
| 108 | TRINITY_DN205_c0_g2_i4 | 939 | 0 | 8.955479842 | Type-II α-neurotoxin | | | AAB87417 | Naja sputatrix |
| 109 | TRINITY_DN0_c0_g1_i3 | 481357 | 0 | 14.64856614 | Type-II α-neurotoxin | | | AAL30059 | Bungarus candidus |
| 110 | TRINITY_DN525_c0_g1_i6 | 7708 | 0 | 8.955479842 | Type-II α-neurotoxin | | | ETE56205 | Ophiophagus hannah |
| 111 | TRINITY_DN0_c0_g1_i19 | 144778 | 0 | 17.85990133 | Unconventional 3FTx | | | AAD41806 | Bungarus multicinctus |
| 112 | TRINITY_DN93_c0_g1_i3 | 96349 | 0 | 8.955479842 | Unconventional 3FTx | **11.70%** | | ETE56933 | Ophiophagus hannah |
| 113 | TRINITY_DN0_c0_g1_i4 | 66221 | 0 | 17.85990133 | Unconventional 3FTx | | | AAD41806 | Bungarus multicinctus |
| 114 | TRINITY_DN93_c0_g1_i2 | 2317 | 0 | 17.66841843 | Unconventional 3FTx | | | ETE56933 | Ophiophagus hannah |
| 115 | TRINITY_DN430_c0_g1_i3 | 43017 | 0 | 8.173542529 | κ-bungarotoxin | **1.63%** | | CAB46659 | Bungarus multicinctus |
| 116 | TRINITY_DN397_c1_g1_i1 | 60867 | 227 | 8.173542529 | VEGF | | | JAA74898 | Pseudonaja modesta |
| 117 | TRINITY_DN397_c1_g1_i7 | 13513 | 0 | 8.173542529 | VEGF | | | XP_026551346 | Pseudonaja textilis |
| 118 | TRINITY_DN397_c1_g1_i4 | 11872 | 142 | 8.173542529 | VEGF | | | XP_026551346 | Pseudonaja textilis |
| 119 | TRINITY_DN397_c1_g1_i5 | 8991 | 50 | 8.173542529 | VEGF | | | JAA74898 | Pseudonaja modesta |
| 120 | TRINITY_DN397_c1_g1_i2 | 5797 | 0 | 5.543356523 | VEGF | | | XP_026551346 | Pseudonaja textilis |
| 121 | TRINITY_DN4427_c0_g1_i1 | 3442 | 139 | 5.543356523 | VEGF | | **1.45%** | LAB17921 | Micrurus spixii |
| 122 | TRINITY_DN4427_c0_g1_i4 | 763 | 0 | 8.173542529 | VEGF | | | JAB52939 | Micrurus fulvius |
| 123 | TRINITY_DN397_c1_g1_i3 | 617 | 8 | 5.543356523 | VEGF | | | XP_026551346 | Pseudonaja textilis |
| 124 | TRINITY_DN1402_c0_g1_i2 | 267 | 0 | 2.050467242 | VEGF | | | XP_015682072 | Protobothrops mucrosquamatus |
| 125 | TRINITY_DN47628_c0_g1_i1 | 201 | 73 | 5.543356523 | VEGF | | | LAB47472 | Micrurus surinamensis |
| 126 | TRINITY_DN5400_c0_g1_i1 | 153 | 71 | 5.543356523 | VEGF | | | LAA87755 | Micrurus lemniscatus lemniscatus |
| 127 | TRINITY_DN4427_c0_g1_i3 | 111 | 0 | 21.27715236 | VEGF | | | LAB17921 | Micrurus spixii |
| 128 | TRINITY_DN134_c0_g1_i14 | 302713 | 0 | 21.27715236 | β-bungarotoxin | | | CAA37482 | Bungarus multicinctus |
| 129 | TRINITY_DN134_c0_g1_i9 | 211831 | 0 | 21.27715236 | β-bungarotoxin | | | ABU63164 | Bungarus fasciatus |
| 130 | TRINITY_DN134_c0_g1_i8 | 107475 | 0 | 21.27715236 | β-bungarotoxin | | | AAL87003 | Bungarus caeruleus |
| 131 | TRINITY_DN134_c0_g1_i10 | 57814 | 0 | 16.05485143 | β-bungarotoxin | | | AAL87003 | Bungarus caeruleus |
| 132 | TRINITY_DN660_c0_g1_i3 | 18152 | 0 | 18.8570687 | β-bungarotoxin | | **9.55%** | ABG90493 | Bungarus multicinctus |
| 133 | TRINITY_DN485_c0_g1_i10 | 4243 | 0 | 18.8570687 | β-bungarotoxin | | | AAL30067 | Bungarus candidus |
| 134 | TRINITY_DN485_c0_g1_i9 | 1069 | 0 | 16.05485143 | β-bungarotoxin | | | AAL30067 | Bungarus candidus |
| 135 | TRINITY_DN660_c0_g1_i4 | 703 | 0 | 16.6518196 | β-bungarotoxin | | | ABG90493 | Bungarus multicinctus |
| 136 | TRINITY_DN5510_c0_g2_i2 | 358 | 0 | #N/A | β-bungarotoxin | | | BAD06267 | Bungarus candidus |

**Table S7A.** Toxicity profiles of *Bungarus* species from Western India.

| Sample ID | Venom Dose (µg) | | | | | Number of survivors | | | | | $LD_{50}$ (µg/mouse) | $LD_{50}$ (mg/Kg) |
|---|---|---|---|---|---|---|---|---|---|---|---|---|
| *B. sindanus* | 0.31 | 0.39 | 0.49 | 0.61 | 0.76 | 5 | 3 | 2 | 0 | 0 | 0.44<br>0.37- 0.52 | 0.022<br>0.018- 0.026 |
| *B. caeruleus* | 2.88 | 3.6 | 4.5 | 5.62 | 7.03 | 5 | 5 | 3 | 2 | 0 | 5.03<br>4.25- 5.95 | 0.251<br>0.213- 0.297 |
| *B. romulusi* | 1 | 1.28 | 1.6 | 2 | 2.5 | 2 | 1 | 0 | 0 | 0 | 0.90<br>0.734- 1.10 | 0.045<br>0.037- 0.055 |

This table contains the median lethal doses and survivorship data for venoms of *Bungarus* species.

**Table S7B.** Neutralisation potencies of commercial Indian polyvalent antivenoms against *Bungarus* venoms.

| Sample ID | **Antivenom used:** Indian polyvalent antivenom manufactured by Premium Serums and Vaccines Pvt. Ltd. (Batch No. ASVS-I Lyo.013) | | | | | | |
|---|---|---|---|---|---|---|---|
| | **Amount of antivenom injected in the venom-antivenom mixture** (μL) | | | | **$ED_{50}$** (μL) | **$ED_{50}$** (μL antivenom/mg venom) | **Potency of antivenom** (mg/mL) |
| *B. sindanus* | 14.63 | 9.76 | 6.51 | 4.34 | 11.62<br>8.60- 15.68 | 5,281.82<br>3909.09- 7127.27 | 0.152<br>0.11- 0.20 |
| *B. caeruleus* | 32.93 | 21.96 | 14.63 | 9.75 | 26.90<br>23.21- 31.16 | 1069.58<br>922.86- 1238.97 | 0.74<br>0.64- 0.86 |
| *B. romulusi* | 32.93 | 21.96 | 14.63 | 9.75 | 26.90<br>23.21- 31.16 | 5977.78<br>5157.78- 6924.44 | 0.13<br>0.11- 0.15 |

The neutralisation potencies of commercial Indian polyvalent antivenom against the venoms of *Bungarus* species are shown in the table above.
